# A computational study of the fold and stability of cytochrome c with implications for disease

**DOI:** 10.1101/2024.07.24.604913

**Authors:** Muhammad Abrar Yousaf, Massimiliano Meli, Giorgio Colombo, Anna Savoia, Annalisa Pastore

**Affiliations:** Department of Neurosciences, Biomedicine and Movement Sciences, University of Verona, Verona, Italy; Institute of Chemical Sciences and Technologies ‘’Giulio Natta’’ - SCITEC, National Research Council (CNR), Milan, Italy; Department of Chemistry, University of Pavia, Pavia, Italy; Department of Engineering for Innovation Medicine, University of Verona, Verona, Italy; Department of Clinical Neuroscience, King’s College London, Denmark Hill Campus, London, United Kingdom; Elettra Sincrotrone Trieste, s.s. 14 km 163,500 in Area Science Park, Basovizza, Trieste, Italy

**Keywords:** Cytochrome c, thrombocytopenia 4, mutations, peroxidase activity, molecular dynamics simulation

## Abstract

Cytochrome c (Cyt-c), encoded by the *CYCS* gene, is crucial for electron transport, peroxidase activity, and apoptosis. Mutations in *CYCS* causes thrombocytopenia 4, a disorder with low platelet counts. We have, for instance, recently described six Italian families with five different heterozygous missense *CYCS* variants. These mutations likely enhance peroxidase and apoptotic activities, yet the mechanisms causing reduced platelet production and increased apoptosis are unclear. This study investigates clinically-related Cyt-c variants using an integrated bioinformatics approach. Our findings reveal that all variants are at evolutionarily conserved sites, potentially disrupting Cyt-c function and contributing to disease phenotypes. Specific variants are predicted to affect phosphorylation (T20I, V21G, Y49H), and ubiquitination (G42S, A52T, A52V, T103I). Molecular dynamics simulations (500 ns) revealed significant structural differences from the wild-type protein, with mutants showing reduced stability and increased unfolding and flexibility, particularly in the Ω-loops. These changes result in the displacement of the Ω-loops away from the heme iron, weakening critical hydrogen bonds and consequently opening the heme active site. This open conformation may enhance accessibility to small molecules such as H₂O₂, thereby promoting peroxidase activity, which may enhance apoptosis and likely impact megakaryopoiesis and platelet homeostasis in THC4.

## 1 Introduction

Cytochrome C (Cyt-c) is an essential small heme-binding protein [1] that functions as an electron carrier in the electron transport chain by transferring electrons from complex III to cytochrome oxidase [2–4]. Cyt-c is a small globular protein of 105 residues in humans that contains a single covalently bound heme group attached to Cys15 and Cys18 of a CXXCH motif with two thioether bonds. His19 and Met81 are axial ligands of the six-coordinated heme iron. The protein fold comprises five α-helices connected by three so-called for historic reasons Ω-loop regions, namely distal (residues 72-86), central (residues 41-58) and proximal (residues 13-29 and 28-40) Ω-loops [5–7]. These are non-regular protein structural motifs that consist of loops of six or more amino acid residues (Fetrow, 1995). Heme ligation remains similar both in the ferric and ferrous forms of Cyt-c, apart from the so-called alkaline transition of ferric Cyt-c. This is a pH-dependent conformational change, first identified more than 80 years ago [8], in which loss of the axial ligand Met80 leads to a mixture of two species with chemically equivalent axial coordination pattern. Although the details of this conformational change remain elusive, the resulting structure is thought to be globular and similasr to that of the wild-type protein at neutral pH, with the most important deviations localized in the 70–85 loop region. Being highly conserved, Cyt-c is attached to the inner membrane of the mitochondrion to fulfil multiple important functions [9] which include a role in the intrinsic apoptotic pathway where Cyt-c initiates cell apoptosis as an antioxidant agent [10–12]. Because of its essential cellular role, it is not a surprise that Cyt-c is associated to pathology: heterozygous mutations of the *CYCS* gene on chromosome 7p15 are associated to autosomal dominant non-syndromic thrombocytopenia 4 (THC4, OMIM: 612004) [13], a disease originally described in a New Zealander family and, more recently, found also in a few other families all over the world [13–16]. Thrombocytopenia are a heterogeneous group of diseases caused by mutations in as many as at least twenty different genes [17]. Some of these diseases are rare, such as it is the case for THC4. Despite the important roles of Cyt-c, it is interesting to notice that THC4 patients present only a mild phenotype that usually consists of a decreased number of platelets in circulating blood [13,14] caused by ectopic platelet release from bone marrow [13]. Platelets from THC4 patients are otherwise normal in size and morphology, and no increased bleeding tendency is usually described. While the low severity of this condition could suggest that a better understanding of the mechanisms causing it may be uninteresting, the need of being able to distinguish this condition from other more serious pathologies is essential for a correct diagnosis which may avoid mistreatment of the patients. There have in fact been cases of THC4 that were incorrectly treated by steroids and/or splenectomy, without any significant benefit [18].

Various Cyt-c variants have been so far identified that cause THC4 [13–16,19–21]. Amongst them, G42S and Y49H have been extensively studied and the crystal structures of some variants have been solved [22,23]. More recently, we have reported new variants and characterized the six families of the carriers for their symptoms [18]. By a clinical characterization of the 22 affected individuals, the largest series of THC4 patients ever reported, we showed that this disorder is characterized by mild-to-moderate thrombocytopenia, normal platelet size, but no additional clinical phenotypes associated with reduced platelet count. We also identified a significant correlation between the region of Cyt-c affected by mutations and the extent of thrombocytopenia, which suggests different degrees of impairment of Cyt-c functions caused by different pathogenetic variants.

In the present study, we present a comprehensive bioinformatics analysis of all currently known variants of Cyt-c, examining the structural features these variants induce in the Cyt-c fold through various computational tools and molecular dynamics simulations. We also explore how the structural roles of certain residues correlate with observed phenotypes affecting protein function. Hence, this study aims to propose new hypotheses regarding the impact of different Cyt-c variants on protein folding and stability, with the goal of elucidating the molecular mechanisms underlying enhanced peroxidase activity, increased apoptosis, and their effects on platelet production in THC4. These findings could offer new insights into the role of Cyt-c in the context of disease.

## 2 Methodology

### 2.1 Data collection and multiple sequence alignment

This study describes the functional effects of fourteen single amino acid variants (T20I, V21G, H27Y, N32H, L33V, G42S, Y49H, A52T, A52V, R92G, A97D, Y98H, L99V, and T103I) in human Cyt-c using bioinformatics approaches. These variants have been reported in patients with or without clinical characterization or identified through sequencing analysis (Table 1). The three-dimensional structure of Cyt-c (PDB ID: 3ZCF) was retrieved from the RCSB PDB database (https://www.rcsb.org/) [24]. Notably, the amino acid numbering in our study is offset by +1 from that in the PDB file, reflecting different sequence annotation conventions. Protein amino acid sequences of *Homo sapiens* and 112 other species [25] were retrieved either from UniProt (https://www.uniprot.org/) [26] or NCBI [27], belonging to diverse taxonomic groups including mammals, birds, reptiles, amphibians, fish, insects, crustaceans, mollusks, annelids, echinoderms, nematodes, fungi, protozoa, green algae, flowering plants, and gymnosperms. This broad selection allows us for the thorough analysis of evolutionary lineages to identify the conserved residues. A multiple sequence alignment was performed using the ClustalW tool (https://www.genome.jp/tools-bin/clustalw) [28] (**Supplementary data, Table S1**) to facilitate the evolutionary conservation analysis of protein residues across different species. The visualization was performed using WebLogo 3 [29]. The residues are numbered according to the human Cyt-c.

**Table 1.** THC4-associated Cyt-c variants analyzed in the present study.

| Protein | References<br>(with molecular and/or clinical characterization) | References<br>(without clinical characterization) |
| --- | --- | --- |
| T20I |  | [30] |
| V21G |  | [30] |
| H27Y | [31] |  |
| N32H |  | [30] |
| L33V |  | [30] |
| G42S | [13,18] | [21,30,32] |
| Y49H | [14,18] |  |
| A52T | [18] | [30,33] |
| A52V | [15] |  |
| R92G | [18] | [34] |
| A97D |  | ClinVar (ISTH-SSC Genomics) |
| Y98H |  | [35] |
| L99V | [18] | [21] |
| T103I | [32] |  |

### 2.2 Prediction of functional effect and pathogenicity of variants

The functional impacts of Cyt-c variants were assessed using SIFT (https://sift.bii.a-star.edu.sg/) [36]. SIFT predicts the effects of amino acid substitutions on protein function based on their position and nature, with scores ≤ 0.05 indicating deleterious changes. For the association of variants with the disease, the SuSPect tool (http://www.sbg.bio.ic.ac.uk/~suspect/) [37] was utilized. This tool uses sequence, structural, and systems biology features to predict the phenotypic effects of genetic variants. Trained on seventy-seven distinct features, SuSPect distinguishes between neutral and disease-causing variants, assigning scores ranging from 0 (neutral) to 100 (disease), with a cutoff score of 50 to differentiate between the two types of variants.

### 2.3 Prediction of molecular effects of variants

The MutPred2 platform (http://mutpred.mutdb.org/) [38] predicted molecular effects for Cyt-c variants, utilizing structural, functional and evolutionary data. This tool integrates PSI-BLAST, SIFT and Pfam for profiling, with TMHMM, MARCOIL and DisProt algorithms for predicting transmembrane helices, coiled-coil and disordered regions, respectively. This integration aids in the assessment of structural damage.

### 2.4 Prediction of post-translational modification

Post-translational modifications (PTMs) are crucial for predicting protein function. Methylation sites were identified using GPS-MSP (https://msp.biocuckoo.org/) [39], whereas phosphorylation sites on Serine (S), Threonine (T), and Tyrosine (Y) were predicted using NetPhos 3.1 (https://services.healthtech.dtu.dk/services/NetPhos-3.1/) [40], with scores above 0.5 indicating likely phosphorylation. PhosphoSitePlus (https://www.phosphosite.org/homeAction) [41] facilitated the estimation of ubiquitination sites.

### 2.5 Analysis of gene and protein interactions

Cyt-c mutations may have effects on the associated genes and proteins. The *CYCS* gene interactions were analyzed using GeneMANIA (https://genemania.org/) [42] to predict and understand gene-gene relationships comprehensively. Further, the STRING database (version 12.0) (https://string-db.org/) [43] was utilized to predict protein-protein interactions for Cyt-c, assessing the impact of variants on specific interacting proteins. A high confidence threshold of 0.7 was applied in the analysis to minimize the likelihood of false positives and negatives, ensuring that only interactions with substantial supporting evidence were considered.

### 2.6 Prediction of protein stability

Predicting the stability of mutant proteins is crucial for understanding their activity, function, and regulation. The impact of variants on protein stability was assessed by calculating the difference in Gibbs Free Energy (ΔΔG, in kcal/mol) between the wild-type (ΔG_WT_) and mutant (ΔG_MT_) proteins, where:

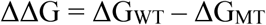

Positive ΔΔG values indicate an increase in stability due to mutations, whereas negative values denote destabilization. For these predictions, we employed both three-dimensional structural and sequence-based methods. DynaMut2 (https://biosig.lab.uq.edu.au/dynamut2/) [44] analyzed the impact of variants on protein dynamics and stability through normal mode analysis of protein three-dimensional structure. MUpro (https://mupro.proteomics.ics.uci.edu/) [45], a sequence-based predictor, utilized a support vector machine model to assess the effects of mutations. These complementary approaches offer a comprehensive understanding of the stability of mutant proteins.

### 2.7 Molecular dynamics simulation

The WT structure of Cyt-c was obtained from the RCSB PDB database (PDB ID: 3ZCF). The starting structures of the mutants T20I, V21G, H27Y, N32H, L33V, G42S, Y49H, A52T, A52V, R92G, A97D, Y98H, L99V, and T103I were generated from the WT structure using PyMOL [46]. The CHARMM36 force field, including HEME parameters derived by Riepl et al. [47] (https://github.com/KailaLab/ff_parameters), was used to model the WT protein and its mutants. Cubic simulation boxes were constructed with dimensions chosen to ensure a 12 Å distance between all protein atoms and the box edges. The boxes were filled with water using the TIP3P water model, and counterions were added to neutralize the system and achieve a final ionic concentration of 0.015 M [48]. All molecular dynamics (MD) simulations were performed using the AMBER 22 suite of programs [49], employing the CUDA implementation for GPUs. The CHARMM36 force field was made compatible with the AMBER MD engine via the Chamber (CHarmm↔AMBER) utility in ParmEd [50,51]. Following parametrization, the systems underwent unrestrained minimization: 5000 steps of steepest descent followed by 5000 steps of conjugate gradient minimization. The minimized systems were equilibrated at 300 K for 10 ns using Langevin dynamics with a collision frequency (γ) of 1 ps^-1^. Subsequently, the equilibrated systems were simulated in the NPT ensemble at 1 atm using weak coupling algorithms [52].

Electrostatic interactions were treated using the particle-mesh Ewald (PME) method [53], and all covalent bonds involving hydrogen atoms were constrained using the SHAKE algorithm [54]. A 2 fs time step and a 10 Å cutoff were applied for nonbonded van der Waals interactions. Production runs of 500 ns at 300 K were performed, with trajectory snapshots saved every 5 ps. For subsequent analysis, trajectory frames were extracted every 10 ps, yielding a total of 50000 frames. The trajectories were then analyzed using the CPPTRAJ module for different structural analyses. The statistics were performed using MS Excel and GraphPad Prism 8. To compare significant differences in MD simulation parameters between WT and mutant proteins, the Kruskal-Wallis test was applied (as the data were non-parametric, indicated by the Shapiro-Wilk test), followed by the Dunn test (p-value < 0.05).

### 2.8 Normal mode analysis

Normal mode analysis was conducted on average protein structures derived from molecular dynamics simulations using iMODS (http://imods.chaconlab.org/) [55]. This analysis utilized an elastic network model with a cutoff of 0.00 Å to establish the network of interacting residues. We investigated 20 low-frequency modes, disregarding clusters and deformations. This method assessed the intrinsic flexibility of the protein, potentially elucidating mechanisms essential to its biological function. Results were visualized through B-factor/mobility maps, eigenvalue plots, and the elastic network model. For covariance matrices, structural clustering was performed using the method developed by Daura et al., as implemented in GROMACS, with a cutoff of 0.2 nm [56,57]; followed by computation of covariance matrices via the ENM (Elastic Network Models) 1.0 server of Bahr group [58]. To assess the impact of mutations, a difference matrix was generated using the WT covariance matrix as a reference. Then plots were produced by replotting the difference matrices using a custom Python script.

## 3 Results

### 3.1 Cyt-c and evolutionary conserved residues

The variants analyzed in this study are present in the proximal and central Ω-loops and the C-terminus (**Fig. 1A**). These mutated residues were assessed for evolutionary conservation through multiple sequence alignment and displayed as sequence logos. Amino acid positions critical for protein structure and/or function tend to be more evolutionarily conserved. Consequently, mutations at these conserved sites are usually more deleterious [59–61]. The sequence logo shows the frequency of amino acids at each position in a sequence alignment (**Fig. 1B**). The height of each letter in the logo is proportional to its frequency, and the total height of letters at a position represents the sequence conservation in bits. Larger letters correspond to more conserved residues, while smaller letters indicate more variability. The analysis revealed that residues T20, V21, H27, N32, L33, G42, Y49, A52, R92, A97, Y98, L99, and T103 are conserved across species from diverse taxonomic groups (**Fig. 1B**). This conservation suggests that the described variants T20I, V21G, H27Y, N32H, L33V, G42S, Y49H, A52T/V, R92G, A97D, Y98H, L99V, and T103I are likely deleterious.

**Fig. 1.**
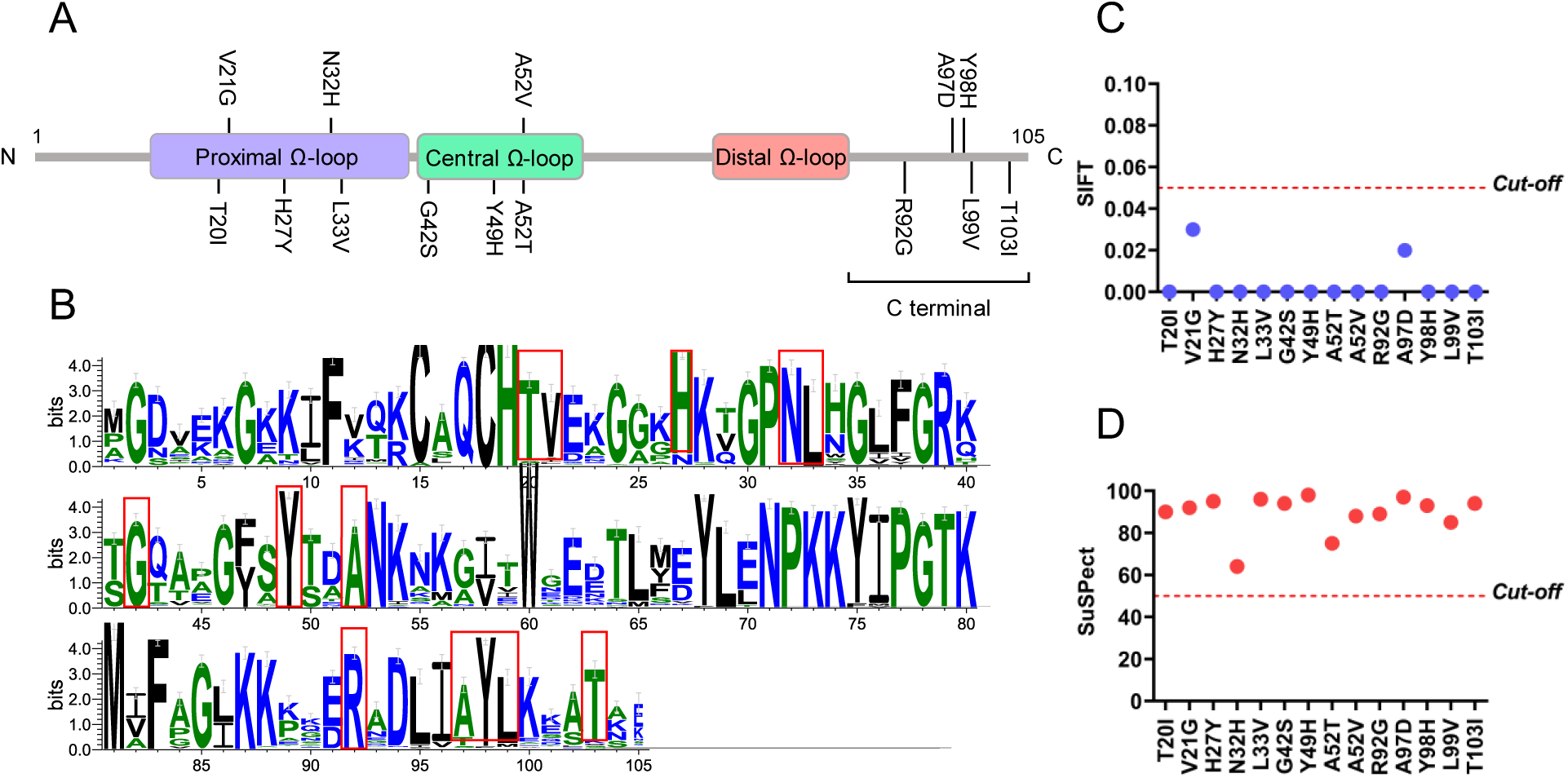
Overall structure of Cyt-c and information on the position and conservation of the clinically important variants. (A) The Cyt-c variants, T20I, V21G, H27Y, N32H and L33V are located in the proximal Ω-loop, G42S, Y49H, A52T and A52V in the central Ω-loop and R92G, A97D, Y98H, L99V and T103I in the protein C-terminus. (B) Sequence logo representing the conservation of amino acids in the protein sequence. The height of each letter at a given position corresponds to the level of conservation, measured in bits, where taller letters indicate higher conservation. Amino acids are color-coded based on their chemical properties: polar residues are green, basic residues are blue, acidic residues are red, and hydrophobic residues are black. Red boxes highlight the positions of the mutated residues: T20I, V21G, H27Y, N32H, L33V, G42S, Y49H, A52T, A52V, R92G, A97D, Y98H, L99V, and T103I. These mutations occur at moderately to highly conserved positions, suggesting potential functional significance. (C) SIFT analysis shows that all variants affect protein function (lower than the cut-off score < 0.05) and (D) SuSPect associates the variants with the disease (greater than the cut-off score > 50).

### 3.2 Prediction of the functional effect and pathogenicity of variants

Analysis of the variants and their predicted impacts on protein function and association with diseases was computed utilizing two widely used computational prediction tools, SIFT, and SuSPect. SIFT scores (<0.05) (**Fig. 1C**) indicate severe effects of all variants on protein function. SuSPect scores (**Fig. 1D**) further support these findings, categorizing each variant with a’Disease’ label, where high scores (>50) suggest a potential association with the disease. This agreement across different predictive platforms underscores the robustness of the predictions. Furthermore, the variants H27Y, G42S, Y49H, A52T, A52V, R92G, and L99V have been described in the literature and are associated with pathogenic effects [13,18,19,22,23,31,62–64], validating our predictions.

### 3.3 Mapping variants on protein structure

The three-dimensional structure of Cyt-c (**Fig. 2A**) shows the presence of Ω-loops with the heme, thereby providing insights into the protein function in electron transport and apoptosis within cells. All Cyt-c variants were mapped onto the protein three-dimensional structure (**Fig. 2B, C, D**). The mutant structures of Cyt-c were generated through the Mutagenesis Wizard in PyMOL [46] using the WT structure (PDB ID: 3ZCF). Ramachandran plots were generated to evaluate the conformational integrity of both WT and mutant structures. These plots assess the phi and psi dihedral angles of the amino acid residues, key indicators of structural feasibility. In all mutant structures, the Ramachandran plot did not reveal any significant differences in the steric hindrance at the mutated residues compared to the WT (**Supplementary data, Fig. S1**), except for G42S (**Fig. 2E**). How the clinically important mutations alter the affected amino acids is shown in **Supplementary data** (**Fig. S2**).

**Fig. 2.**
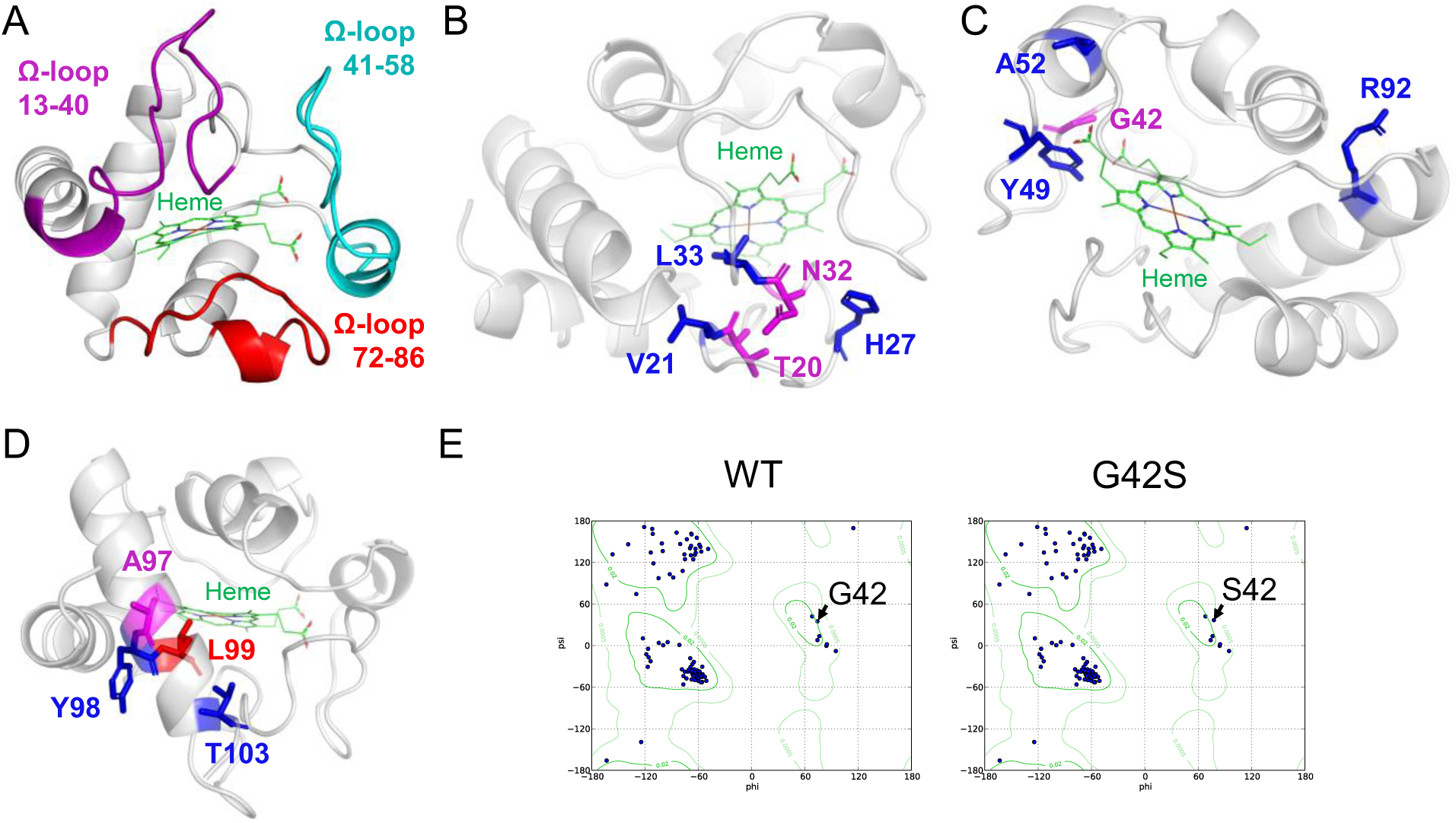
Mapping the currently known clinically important variants on the structure of Cyt-c, and Ramachandran plot analysis. The protein structure is depicted as a grey ribbon, with heme in green. (A) Human Cyt-c (PDB ID: 3ZCF) structure showing three Ω-loop regions: proximal (residues 13-40, magenta), central (residues 41-58, cyan) and distal (residues 72-86, red) Ω-loops, which make the large part of Cyt-c. These Ω-loops are important for both structural and functional reasons, including protein folding and conformational transition. Amino acids investigated for variants are highlighted as follows: (B) T20 (magenta), V21 (blue), H27 (blue), N32 (magenta), L33 (blue); (C) G42 (magenta), Y49 (blue), A52 (blue), R92 (blue); (D) A97 (magenta), Y98 (blue), L99 (red), T103 (blue). Note that the residue numbering is offset by +1 relative to the PDB entry 3ZCF. (E) In the Ramachandran plot, glycine at position 42 (G42) is an exception due to its small size and lack of side chains, which allows it to occupy even the unfavorable regions. In the G42S variant, replacing glycine with serine introduces steric hindrance due to the larger and less flexible residue, positioning it in a less favorable region of the plot.

Buried residues are crucial for the structural stability of proteins because they contribute to the hydrophobic core and are like pillars of the three-dimensional structure. Mutations that destabilize proteins can induce unfolding or misfolding, potentially affecting the functionality of Cyt-c. Therefore, the relative solvent accessibility of mutation sites was calculated using “get_sasa_relative” function of PyMOL [46] that calculates the relative per-residue solvent accessible surface area. Residue with a value of 0 is considered completely buried, and that with a value of 1 is completely exposed. All examined amino acids are buried (< 0.50) within the protein structure; however, the degree of burial varies. Residues like V21 (0.09), R92 (0.06), L99 (0.05), and T103 (0.03) are deeply buried with minimal solvent exposure. T20 (0.15), N32 (0.11), L33 (0.14), G42 (0.14), A97 (0.18), and Y98 (0.12) are also highly buried, though slightly more exposed. Y49 (0.2), and A52 (0.25) have moderate burials, while H27 (0.37) is the most exposed but still considered buried.

The T20I variant in the Cyt-c coil region (**Fig. 2B**), substituting a buried hydrophilic threonine with a larger hydrophobic isoleucine, likely compromises protein stability and disrupts the T20-N32 H-bonds [65]. This is due to the absence of the hydroxyl group in isoleucine, impeding the formation of the H-bond. The V21G variant in a coil region (**Fig. 2B**) introduces a cavity into the hydrophobic core by replacing a medium-sized valine with a smaller, more flexible glycine, potentially affecting protein stability (**Fig. 2B**). The critical role of H27 in a coil region (**Fig. 2B**) for maintaining protein stability is highlighted by the formation of a key H-bond with P45, which ensures the close association of the 20s and 40s Ω-loops [66]. Substituting histidine with the bulkier, uncharged, and hydrophobic tyrosine in the H27Y variant may induce structural alterations that significantly reduce protein stability and disrupt crucial interactions. The replacement of N32 (**Fig. 2B**) with a bulkier histidine in the N32H variant could impair the T20-N32 H-bonding [65] and is likely to affect the protein structural integrity, leading to instability. Additionally, the L33V variant at the turn (**Fig. 2B**) might create a cavity in the hydrophobic core due to the smaller valine size, and the steric differences introduced by the Val mutation could be an additional factor (this is true also for the mutation L99V as mentioned below).

G42, located in a protein loop, exhibits positive phi and psi values in the Ramachandran plot (Fig. 2E). Glycines, with their single hydrogen atom side chain, are symmetric and have a wider range of acceptable phi/psi backbone dihedral angles than other amino acids, allowing glycines to occupy also typically an unfavorable or disallowed region of the Ramachandran plot. Replacing glycine with the bulkier serine could thus lead to steric clashes with adjacent residues due to glycine unique conformational flexibility, significantly altering the loop conformation [67] (Fig. 2E). This could be the reason that G42S mutation displaces residues 42 and 43 about 0.4 Å away from Tyr49, which is part of the core structure of the third omega loop in the experimental structure [22]. Further, the G42S variant (in a coil region, **Fig. 2C**), within the hydrophobic core, introduces a polar and hydrophilic serine, potentially disrupting hydrophobic interactions, destabilizing the core, and affecting the structural integrity and protein function. Y49, located in a buried hydrophobic loop, is a large residue with a bulky phenyl ring and a hydroxyl group (**Fig. 2C**). Replacing Y49 with the medium-sized, imidazole-ring-containing H49 could destabilize the hydrophobic core by creating a cavity. The hydrophilic and basic nature of H49 might disrupt hydrophobic interactions and introduce pH-dependent changes, potentially impacting the protein structure and function. NMR data indicates that the Y49H mutation results in a loss of resonances, suggesting increased conformational heterogeneity, potentially due to partial unfolding of the protein [23,68]. The A52T variant, replacing the nonpolar alanine located in a helix within the hydrophobic core (**Fig. 2C**) with the polar and hydrophilic threonine, could disturb hydrophobic interactions, affecting structure. Similarly, substitution to a larger valine in the A52V variant might affect the structure. In an experimental structure, the A52V variant of causes decreased global stability and decreased stability of the native state relative to sub-global unfolding of Ω-loop [63]. R92, a large, polar and hydrophilic residue located in the core helix near the cytochrome c active site (**Fig. 2C**), is crucial for interactions with the heme and other buried hydrophilic residues. Mutating R92 to the much smaller glycine could disrupt these interactions and create a cavity, compromising the protein function and structure.

The A97D variant in the helix (**Fig. 2D**) may disrupt the protein hydrophobic interactions due to the introduction of the hydrophilic aspartate at position 97, replacing hydrophobic alanine. The Y98H (**Fig. 2D**) variant could compromise the protein structural integrity by creating a cavity and disrupting hydrophobic interactions within the core helix, as it involves replacing the large hydrophobic tyrosine with a medium-sized hydrophilic histidine. Similarly, the L99V variant, located in a helix within the hydrophobic core (**Fig. 2D**), is likely to form a cavity due to the substitution of leucine with the smaller valine. The T103I variant in a coil region (**Fig. 2D**) replaces the small, polar, hydrophilic threonine with a larger, nonpolar, hydrophobic isoleucine, potentially affecting local packing and hydrophilic interactions.

Collectively, these variants could significantly influence the protein by altering its structure and functionality.

### 3.4 Molecular effects of the mutations

We assessed the molecular effects of the mutations of the Cyt-c variants, which could lead to pathogenicity, by the MutPred2 software (**Supplementary data, Table S2**). Mutations in Cyt-c can change its structure and interactions, leading to various molecular effects that may alter its activity. The T20I variant was predicted to affect several aspects of Cyt-c. It could cause the loss of intrinsic disorder, reducing protein flexibility. It may also result in altered metal binding and changes in the protein interaction surface. There is a gain of a functional site at C18, which might influence its activity, but Cyt-c is not primarily known for classical catalytic functions. The V21G could lead to increased flexibility (gain of intrinsic disorder), which may destabilize the protein. It may also change metal binding and stability, both of which are essential for Cyt-c role in electron transfer. This mutation can result in the loss of a functional site at C18, likely affecting how the protein interacts with other molecules. The H27Y was predicted to cause changes in the interaction surface and reduce the protein flexibility. It can also introduce PTMs like acetylation and methylation, which may affect protein stability, although these modifications are not typical for Cyt-c regulation. The mutation may also impact metal binding, which is crucial for its electron transfer function. The N32H has the probability of altering the interaction surface and metal binding, both important for maintaining Cyt-c structure and function. It may also result in the loss of a functional region at K28, potentially affecting the protein interactions in the electron transport chain. The L33V could lead to the gain of an allosteric site, which may introduce new regulatory features. However, it can also alter the interaction surface, possibly disrupting its function in the electron transport chain.

The G42S variant was predicted to cause changes in the interaction surface and solvent accessibility, which may impact how Cyt-c interacts with other molecules in its environment. This mutation may also affect metal binding and the presence of methylation sites, which could influence its structural dynamics. The Y49H likely results in changes to the interaction surface and a loss of phosphorylation at Y49 [3]. It may also alter metal binding and solvent accessibility, which may impact its function in electron transport. The A52T could reduce the solvent accessibility of Cyt-c, potentially impacting key functional sites. It could also affect metal binding and interactions with membranes, which are important for the Cyt-c role in electron transport. Additionally, the variant may introduce acetylation at K56, though this is not a typical regulatory feature for Cyt-c. The R92G has the potential to alter the protein-coiled structure and result in the loss of an allosteric site, potentially affecting the protein shape and function. This mutation can also introduce acetylation and methylation at K87, which could influence protein stability, although these modifications are not commonly linked to Cyt-c function. The A97D, Y98H, and L99V mutations are all predicted to affect the protein-coiled structure and disordered regions, which are important for stability. These mutations can also introduce changes in PTMs and solvent accessibility, which may affect Cyt-c role in electron transport and apoptosis.

### 3.5 Predicting the effects of PTMs

PTMs refer to the chemical modifications that proteins undergo after translation, including methylation, phosphorylation, ubiquitylation, acetylation, and several others. PTMs play a pivotal role in the regulation of protein structures and functions, proving essential across various pathways, including cell signaling and protein-protein interactions, as well as numerous biological processes [69,70]. Mutations at or near the site of these modifications could impact their function, stability, localization, and interaction with other molecules. Our analysis utilized a combination of bioinformatics tools to evaluate the potential effects of Cyt-c mutations on its PTMs, which are crucial for its functionality and regulatory mechanisms. The GPS-MSP tool suggests that native Cyt-c lacks methylation sites, indicating a minimal role of methylation in its regulation. NetPhos 3.1 predicted phosphorylation sites at threonine 20 (T20), tyrosine 47 (Y47), threonine 64 (T64), and threonine 79 (T79). Variants such as T20I, V21G, and Y49H, located at or near these sites, could potentially modify the phosphorylation pattern, thereby affecting the protein activity or its interactions with other cellular molecules. Particularly, the T20I variant would abolish the possibility of phosphorylation at this site, potentially influencing the protein function or stability. Ubiquitination, a modification marking proteins for degradation, was predicted at lysine residues K28, K40, K54, K73, K74, K80, K87, and K100. Variants near these sites (G42S, A52T, A52V, and T103I) may affect the ubiquitination process, thus altering the protein degradation rate or its role in cellular signaling. For instance, previous research has found that Pirh2-mediated ubiquitination of cytochrome c, facilitated by an E3 ubiquitin ligase, plays a critical role in its translocation from mitochondria to the cytosol, impacting mitochondrial function and apoptosis [71].

### 3.6 Effects of variants on interactions

Gene network analysis using the GeneMANIA tool revealed that the *CYCS* gene, encoding Cyt-c, engages with twenty distinct genes (**Fig. 3A**). Mutations in the gene product of *CYCS* could potentially disrupt these interactions. Specifically, *CYCS* was found to co-express with CYC1, VDAC1, VDAC3, and EPS15L1, maintaining similar expression levels across various conditions. It shares a domain only with CYC1. Additionally, *CYCS* demonstrated co-localization with BCL2L1, VDAC1, VDAC2, and CASP9, suggesting that these genes are expressed together in the same cellular locations or tissues. All genes within the same pathway were associated with apoptosis. *CYCS* displayed genetic interactions with BAX, BAK1, and DAP3; all these genes are key players in apoptosis. *CYCS* encodes Cyt-c, which, upon mitochondrial release, initiates apoptosome formation with APAF1 and CASP9, subsequently activating caspases such as CASP3 to drive apoptosis. Genetic interactions between *CYCS* and BAX, BAK1, and DAP3 demonstrate that alterations in *CYCS* function or expression can modulate the apoptotic effects mediated by these genes, underscoring the interconnected nature of the apoptotic pathway and the pivotal role of *CYCS* in regulating apoptosis.

**Fig. 3.**
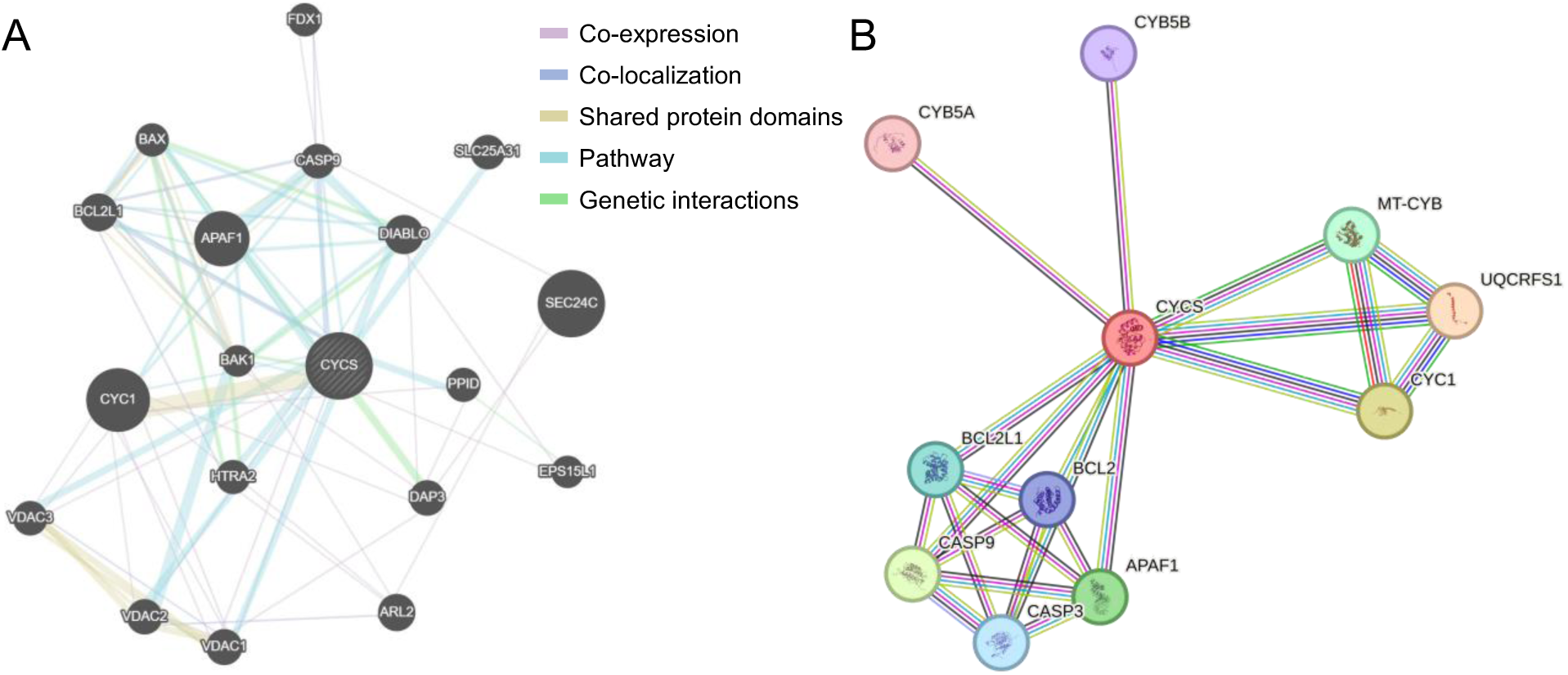
Visualization of gene and protein interactions. (A) Interactions of the *CYCS* gene with other genes, analyzed using the GeneMANIA tool. (B) The protein-protein interaction network of Cyt-c analyzed using the STRING tool. Variants in these networks may affect the functions of interacting genes or proteins.

On the protein level, analyses of protein-protein interactions elucidated the functional relationships among cellular proteins. The STRING database revealed that Cyt-c interacts with ten other proteins, suggesting that mutations in Cyt-c might disrupt their functions (**Fig. 3B**). Notably, Cyt-c was found to interact with QCRFS1, CYC1, CASP9, APAF1, MT-CYB, BCL2L1, CASP3, BCL2, CYB5B, and CYB5A, all with high confidence scores (**Supplementary data, Table S3**). These proteins are involved in the electron transport chain and apoptotic pathway. This analysis highlights the complex network of protein-protein interactions involving Cyt-c, reflecting its critical role in cellular processes.

### 3.7 Gibbs free energy-based protein stability analysis

We assessed the impacts of Cyt-c variants on protein stability by the DynaMut2 and MUpro software, using ΔΔG values which quantify the difference between the energy of the unfolded and folded states of a protein. A positive ΔΔG indicates that a mutant protein is thermodynamically more stable (requiring more energy to unfold) than the WT, whereas a negative ΔΔG suggests it is less stable (requiring less energy to unfold). V21G, H27Y, N32H, L33V, G42S, Y49H, A52T, A52V, R92G, A97D, Y98H, L99V and T103I variants were identified as destabilizing (negative ΔΔG) by both prediction tools (Table 2). Notably, variants carrying the V21G, Y49H, R92G, Y98H and L99V mutations showed the most pronounced destabilizing effects (larger negative ΔΔG), underscoring their significant impact on protein function.

**Table 2.**
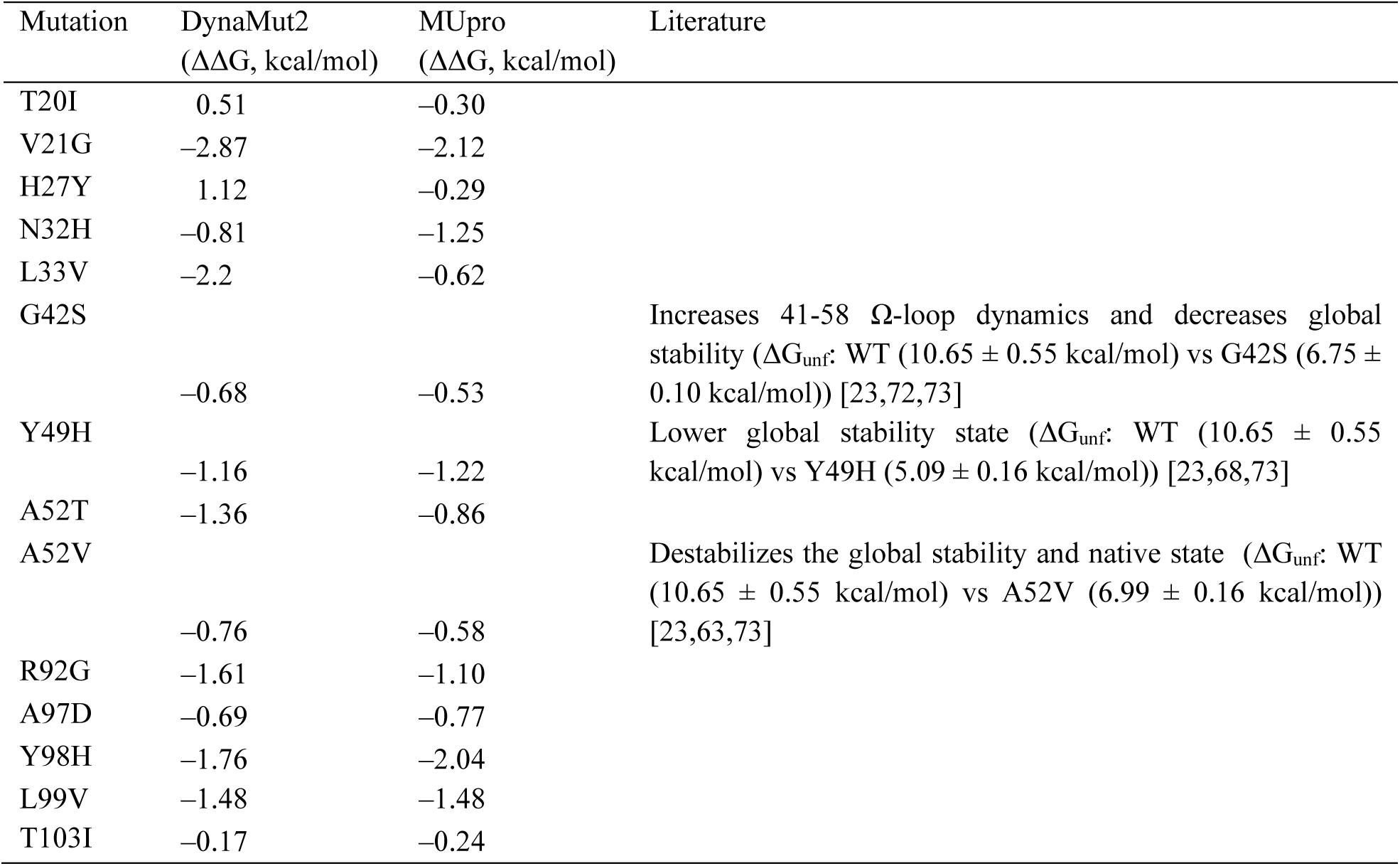
Gibbs free energy (ΔΔG) values for Cyt-c variants computed through DynaMut2 and MUpro servers. Negative ΔΔG indicates destabilizing variants, while positive values suggest the variant as stabilizing. In literature, the lower free energy of unfolding (ΔG_unf_) of G42S, Y49H, and A52V compared to the WT indicates their low stability, validating our predictions.

### 3.8 Dynamics of the wildtype and mutant proteins in aqueous solution

To obtain a more detailed understanding of the structural changes in the precatalytic native state (a six-coordinate heme) of WT and mutant Cyt-c at the molecular level, we carried out MD simulations for all proteins in their oxidized (Fe(III)) state within an aqueous environment. The structural and conformational modifications in the Cyt-c protein have been reported in the initial period during protein unfolding [74,75], and several previous studies conducted simulation analyses till or less than 40 ns [75–79]. However, in this study, we extended the simulations to 500 ns to observe the effects of each protein variant over a prolonged period, potentially providing a more comprehensive analysis.

During the MD simulation, several changes in secondary structures were observed in the mutant proteins compared to the WT. The average structures of the proteins and fluctuations of the secondary structures during the simulation were calculated (**Supplementary data, Fig. S3&S4**). An increase in stable secondary structures, such as β-sheets and α-helices, enhances protein stability due to strong hydrogen bonding and rigid conformations. In contrast, an increase in flexible or disordered structures, including coils, turns, bends, or π-helices, reduces stability by decreasing structural compactness and increasing susceptibility to unfolding. In the WT protein, the secondary structure composition comprises 0.78% parallel β-sheets, 2.32% anti-parallel β-sheets, 1.48% 3-10 helices, 42.67% α-helices, no π-helices, 13.78% turns, 10.42% bends, and 28.54% coils (**Fig. 4 & Supplementary data, Table S4**). Compared to the WT, the T20I mutant shows increases in parallel β-sheets and 3-10 helices, along with a noticeable increase in bends. Conversely, it exhibits decreases in anti-parallel β-sheets, α-helices, turns, and coils. The V21G mutant displays slight increases in parallel and anti-parallel β-sheets, 3-10 helices, α-helices, and coils, as well as a moderate increase in turns, while showing a decrease in bends. The H27Y mutant has increased parallel β-sheets, 3-10 helices, α-helices, and bends, and decreased anti-parallel β-sheets, turns, and coils. The N32H mutant shows a decrease in parallel and anti-parallel β-sheets and 3-10 helices but slight increases in α-helices, turns, bends, and coils. The L33V mutant exhibits decreases in parallel and anti-parallel β-sheets, 3-10 helices, α-helices, and turns, with an increase in bends and a slight increase in coils. The Y49H mutant shows increases in parallel β-sheets, 3-10 helices, and α-helices, while decreases in turns, bends, and coils; anti-parallel β-sheets remain unchanged. The G42S mutant displays increases in parallel and anti-parallel β-sheets and 3-10 helices, a slight increase in bends, and decreases in α-helices, turns, and coils. The A52T mutant shows increases in parallel β-sheets, 3-10 helices, and turns, and decreases in anti-parallel β-sheets, α-helices, and bends, with slightly higher coil content. The A52V mutant has increases in parallel β-sheets, 3-10 helices, α-helices, and bends, and decreases in anti-parallel β-sheets, turns, and coils. The R92G mutant exhibits decrease in parallel and anti-parallel β-sheets and 3-10 helices, with slight increases in α-helices, turns, bends, and coils. The A97D mutant shows a slight increase in parallel β-sheets and increases in turns, bends, and coils, while experiencing decreases in anti-parallel β-sheets, 3-10 helices, and α-helices compared to the WT. The Y98H mutant displays increases in parallel β-sheets, 3-10 helices, and turns, alongside decreases in anti-parallel β-sheets, α-helices, bends, and coils. The L99V mutant exhibits increases in parallel β-sheets, 3-10 helices, α-helices, and bends, while showing decreases in anti-parallel β-sheets, turns, and coils. Lastly, the T103I variant leads to increases in parallel β-sheets, 3-10 helices, α-helices, and turns, while exhibiting decreases in anti-parallel β-sheets, bends, and coils compared to the WT (**Fig. 4A & Supplementary data, Table S4**).

**Fig. 4.**
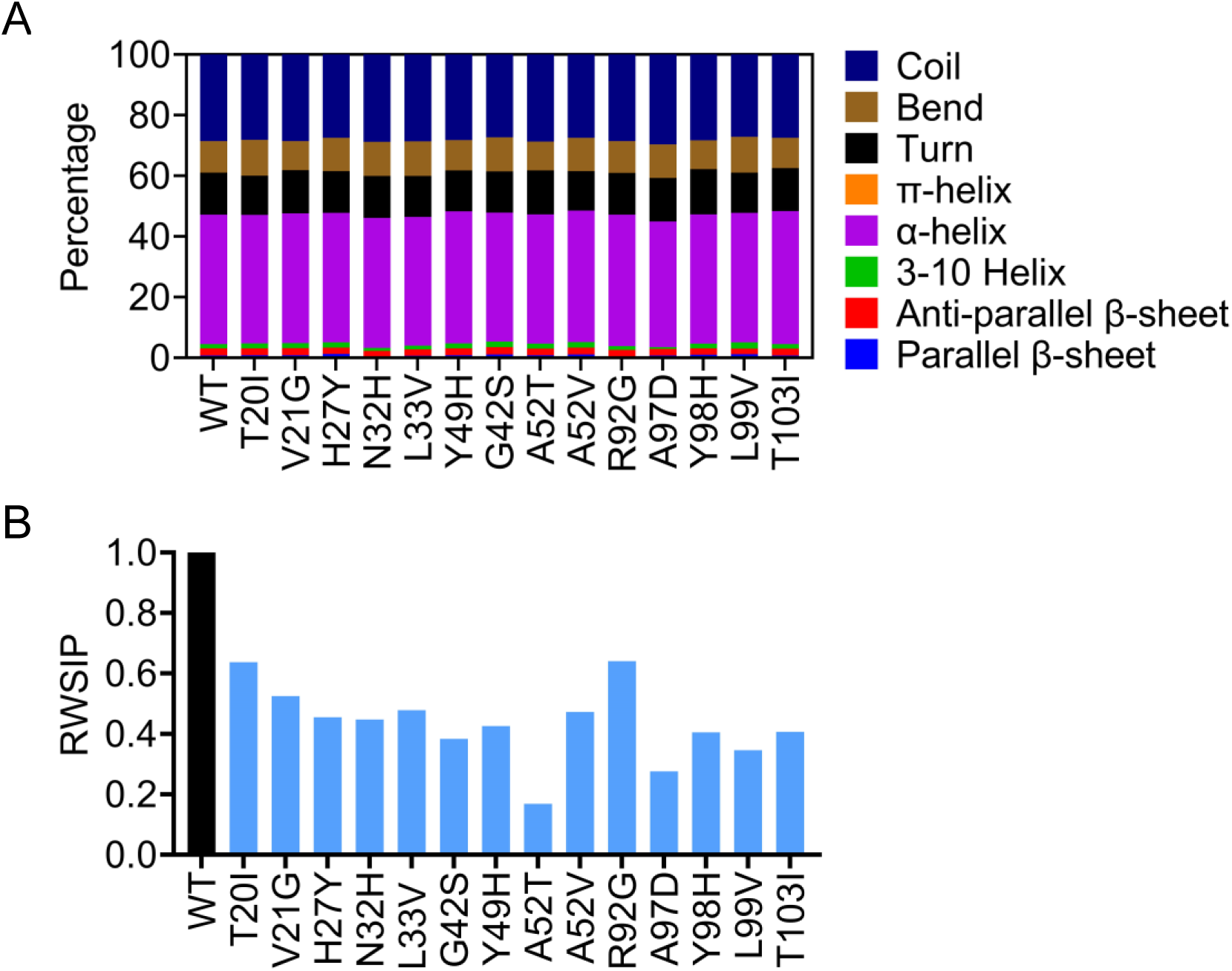
(A) The overall percentage distribution of secondary structure composition of the Cyt-c mutants over the 500 ns MD simulation. The variants display changes in the coil (dark blue), bend (brown), turn (black), π-helix (orange), α-helix (purple), 3-10 helix (green), anti-parallel β-sheet (red), and parallel β-sheet (light blue) during the simulation. (B) Comparison of the root-weighted square inner product (RWSIP) values with the WT, where mutants display significantly low values, compared to the WT.

We then analyzed the root-weighted square inner product (RWSIP) [80] to compare the essential dynamics between WT and mutant proteins. RWSIP accounts for the similarity of essential dynamics and the relative importance (weight) of individual principal components in protein motion. This makes RWSIP sensitive to even subtle shifts in how mutations affect protein dynamics. An RWSIP value of 1 indicates that two simulations span identical conformational space, whereas 0 shows the absence of any superposition in the dynamic evolution of the systems. In our analysis, the WT has an RMSIP of 1, serving as the reference. The RMSIP values (**Fig. 4B)** show that all mutants differ significantly with low values from that of the WT protein, suggesting potential alterations in the conformational dynamic evolution of the native structural fold.

Collectively, these findings demonstrate that these variants can lead to significant variations in protein stability by altering the prevalence and type of secondary structure elements.

### 3.9 Effect of the mutations on protein conformational stability

The root mean square deviation (RMSD) along an MD trajectory is a common quantitative measure of how much a structure deviates from a reference conformation, typically its starting or average structure. It is usually taken as an indirect measure of the fold stability and/or protein flexibility [81]. Among the analyzed structures, the WT Cyt-c Cα displays low RMSD, indicating a stable conformation similar to its initial structure (**Fig. 5A-C**). Conversely, statistical analysis (**Supplementary data, Table S5**) shows that most variants significantly increase RMSD compared to the WT, indicating decreased Cyt-c stability. Mutants T20I, V21G, N32H, L33V, G42S, Y49H, A52V, R92G, A97D, L99V, and T103I exhibit significantly higher RMSD values across the entire protein structure, suggesting increased conformational fluctuations and decreased stability (**Fig. 5A-C**, Cα). However, H27Y, A52T, and Y98H displayed overall better stability compared to the WT based on lower RMSD values of the whole structure.

**Fig. 5.**
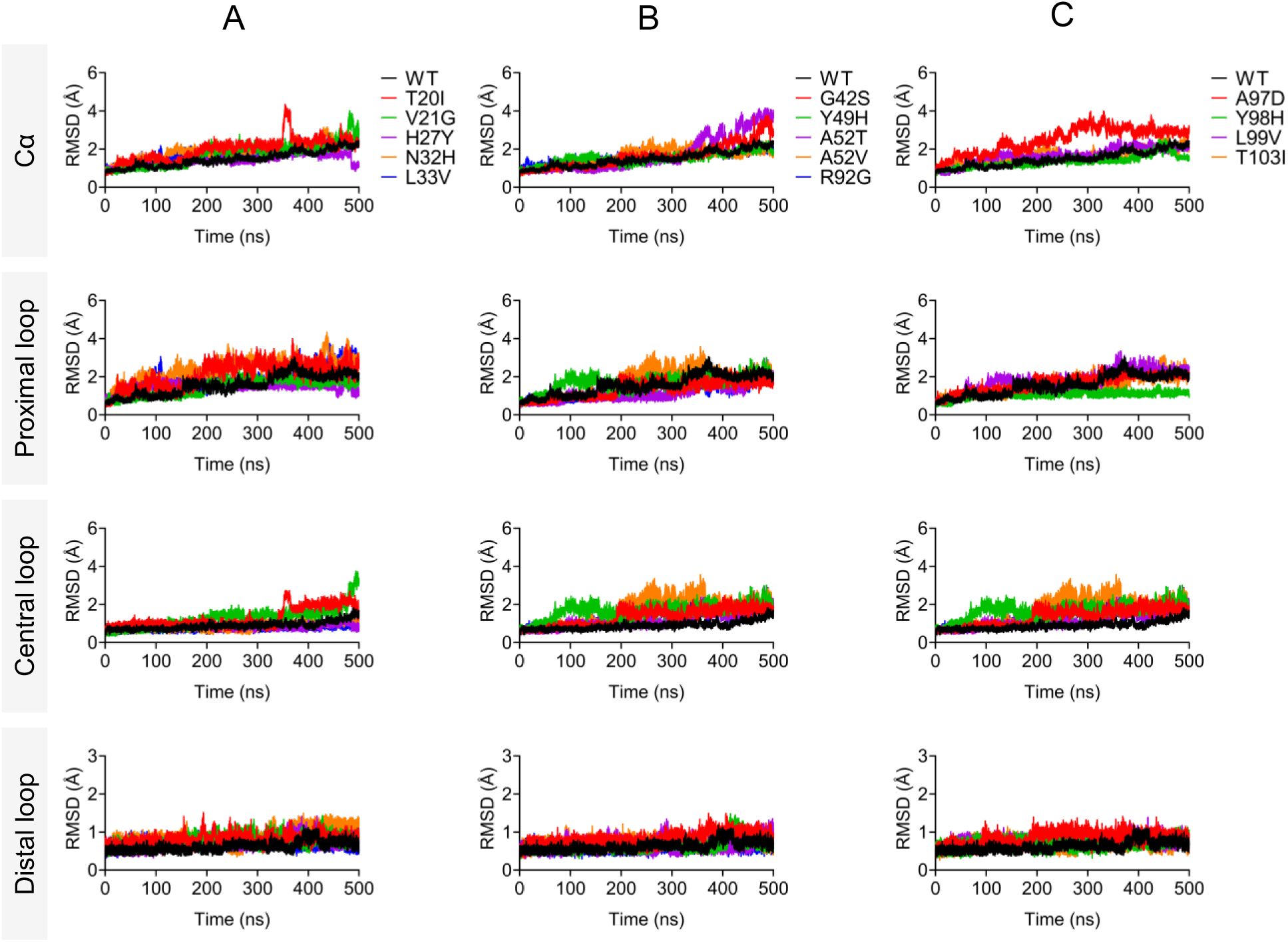
Root Mean Square Deviation (RMSD) profiles of the entire protein backbone (Cα atoms) and specific regions (proximal, central, and distal Ω-loops) in the WT and mutant Cyt-c, illustrating structural fluctuations and stability over the 500 ns MD simulation. (A) RMSD variations of the whole protein and Ω-loops for WT (black) and mutants T20I (red), V21G (green), H27Y (purple), N32H (orange), and L33V (blue). (B) Comparison of RMSD values for whole protein (Cα atoms) and Ω-loops of WT (black), G42S (red), Y49H (green), A52T (purple), A52V (orange), and R92G (blue). (C) RMSD values for the entire protein (Cα atoms) and localized Ω-loops in WT (black) and mutants A97D (red), Y98H (green), L99V (purple), and T103I (orange). The plots highlight the effects of specific mutations on the overall stability and localized structural variations in the protein.

Region-specific analysis reveals how variants variably affect the stability of the Ω-loops within a protein structure, focusing on the proximal, central, and distal Ω-loops. In the proximal Ω-loop, variants at T20I, V21G, N32H, L33V, Y49H, A52V, A97D, and L99V significantly destabilize this region (**Fig. 5A-C**, Proximal loop). In the central Ω-loop, variants T20I, V21G, G42S, Y49H, A52T, A52V, R92G, A97D, Y98H, L99V, and T103I lead to considerable destabilization (Fig. 5A-C, Central loop). The distal Ω-loop is significantly destabilized by all the analyzed variants except R92G, whose effect is not statistically significant (**Fig. 5A-C**, Distal loop).

### 3.10 Effect of the mutations on protein compactness and folding

The radius of gyration (Rg) is a critical parameter in MD simulations, widely used to evaluate protein compactness and structural stability. Lower Rg values represent tightly packed and stable protein conformations, while higher values indicate extended or unfolded structures. In this study, the WT Cyt-c consistently shows reduced Rg for the entire structure, as well as within its functional Ω-loops (**Fig. 6A-C**). This uniform Rg decrease suggests an overall trend toward greater compactness and stability in WT Cyt-c.

**Fig. 6.**
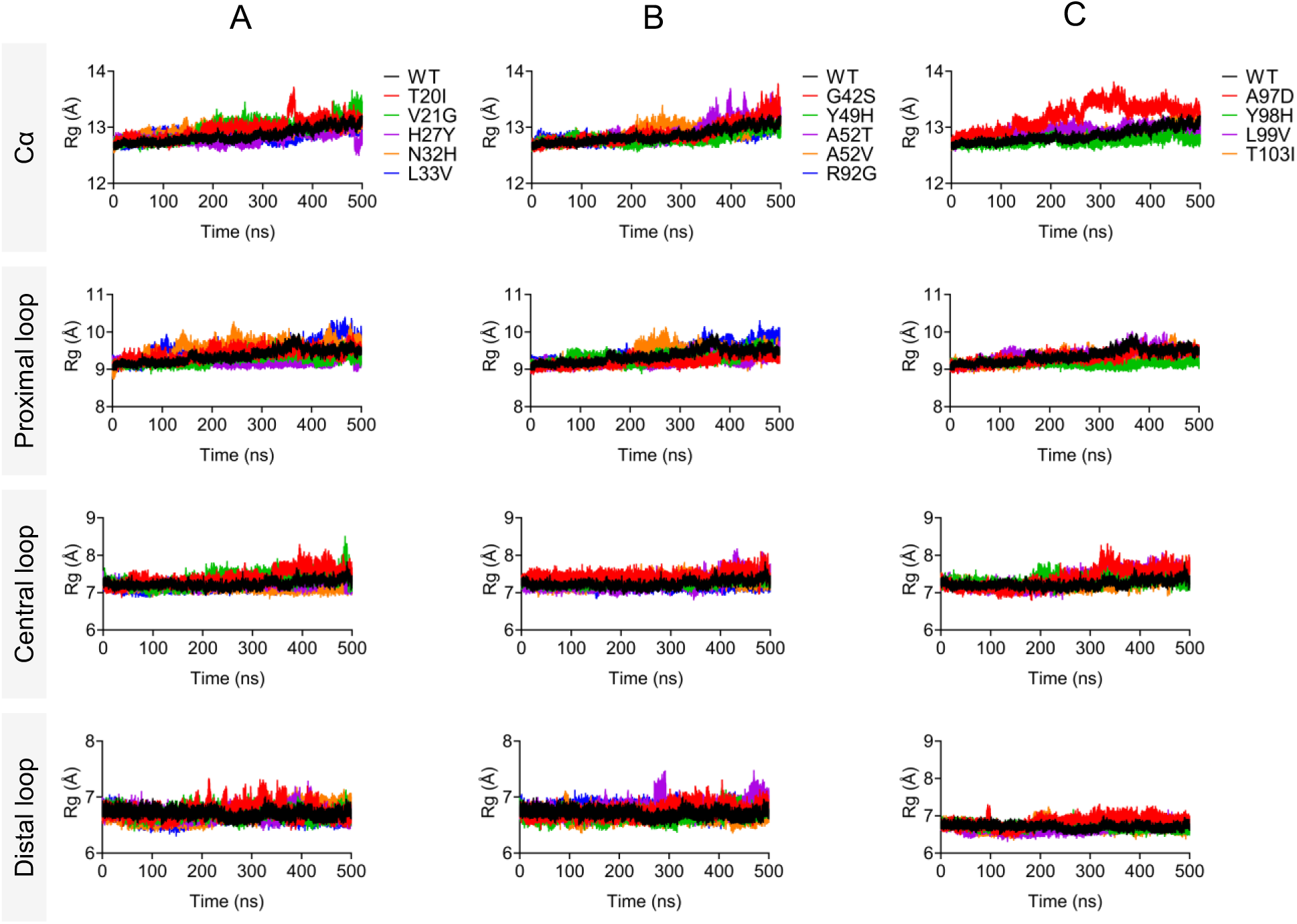
Radius of gyration (Rg) profiles of the entire protein backbone (Cα atoms) and specific regions (proximal, central, and distal Ω-loops) in the WT and mutant Cyt-c proteins, highlighting the effect of variants on protein compactness over the 500 ns MD simulation. (A) Rg variations for the whole protein (Cα atoms) and Ω-loops in WT (black) and mutants T20I (red), V21G (green), H27Y (purple), N32H (orange), and L33V (blue). (B) Rg profiles for the whole protein (Cα atoms) and Ω-loops in WT (black) and mutants G42S (red), Y49H (green), A52T (purple), A52V (orange), and R92G (blue). (C) Rg variations for the whole protein (Cα atoms) and Ω-loops in WT (black) and mutants A97D (red), Y98H (green), L99V (purple), and T103I (orange). The data highlights how each variant impacts protein compactness and Ω-loop stability, revealing structural changes associated with specific mutations.

In contrast, Rg analysis of several Cyt-c variants reveals significant structural deviations that reflect reduced compactness and increased unfolding compared to the WT protein (**Supplementary data, Table S6)**. Mutants including T20I, V21G, N32H, L33V, G42S, A52T, A52V, R92G, A97D, L99V, and T103I display higher Rg values compared to the WT, indicating structural unfolding; except H27Y, Y49H, and Y98H that show lower Rg values (**Fig. 6A-C**, Cα). These variants influence both the global protein structure and specific regions within the functional Ω-loops, disrupting the overall stability of Cyt-c. The proximal loop is particularly affected by T20I, N32H, L33V, A52V, and R92G, which are associated with significant unfolding in this region (**Fig. 6A-C**, Proximal loop). Most of the variants, T20I, V21G, G42S, Y49H, A52T, A52V, A97D, Y98H, and L99V lead to reduced compactness and structural destabilization of the central loop (**Fig. 6A-C**, Central loop). In the distal loop, N32H, G42S, A52T, R92G, A97D, Y98H, and T103I lead to increased unfolding and decreased compactness, contributing to the destabilized protein conformation (**Fig. 6A-C**, Distal loop). The findings demonstrate that the specific variants in Cyt-c alter local loop dynamics and overall compactness, affecting its stability and functionality and providing insights into the impact of sequence variations on protein structure.

### 3.11 Effect of the mutations on protein flexibility

Root mean square fluctuation (RMSF) analysis is a key tool for quantifying the flexibility of amino acid residues within a protein structure during simulations. Higher RMSF values correspond to greater residue mobility, often linked to reduced protein stability and potential functional alterations. Our analysis reveals that the T20I, V21G, N32H, G42S, A52T, A97D, and L99V variants exhibit increased flexibility compared to the WT protein, as indicated by elevated RMSF values (**Fig. 7A-C**, Cα), though, all the higher values were not statistically significant (**Supplementary data, Table S7**). In contrast, L33V, Y49H, and Y98H depict lower flexibility for the entire structure. The detailed examination of specific protein Ω-loops shows that flexibility is increased across most variants, with notable impacts in the proximal, central, and distal loops. The proximal Ω-loop demonstrates enhanced mobility in the T20I, V21G, L33V, G42S, A52T, A52V, R92G, A97D, Y98H, and L99V mutants, suggesting that these variants in this region can destabilize local structural interactions (**Fig. 7A-C**, Proximal loop). Similarly, the central loop exhibits heightened flexibility caused by the T20I, V21G, G42S, A52T, and A97D variants (**Fig. 7A-C**, Central loop). In the distal loop, increased residue flexibility is observed in the T20I, V21G, G42S, A52T, A97D, and L99V variants, further contributing to the destabilization of the protein functional regions (**Fig. 7A-C**, Distal loop). These findings underscore the impact of specific mutations on residue flexibility, particularly in critical Ω-loops, where increased RMSF values suggest reduced structural stability and potential functional impairment. This analysis sheds light on how mutations affect protein dynamics, revealing their structural and functional consequences.

**Fig. 7.**
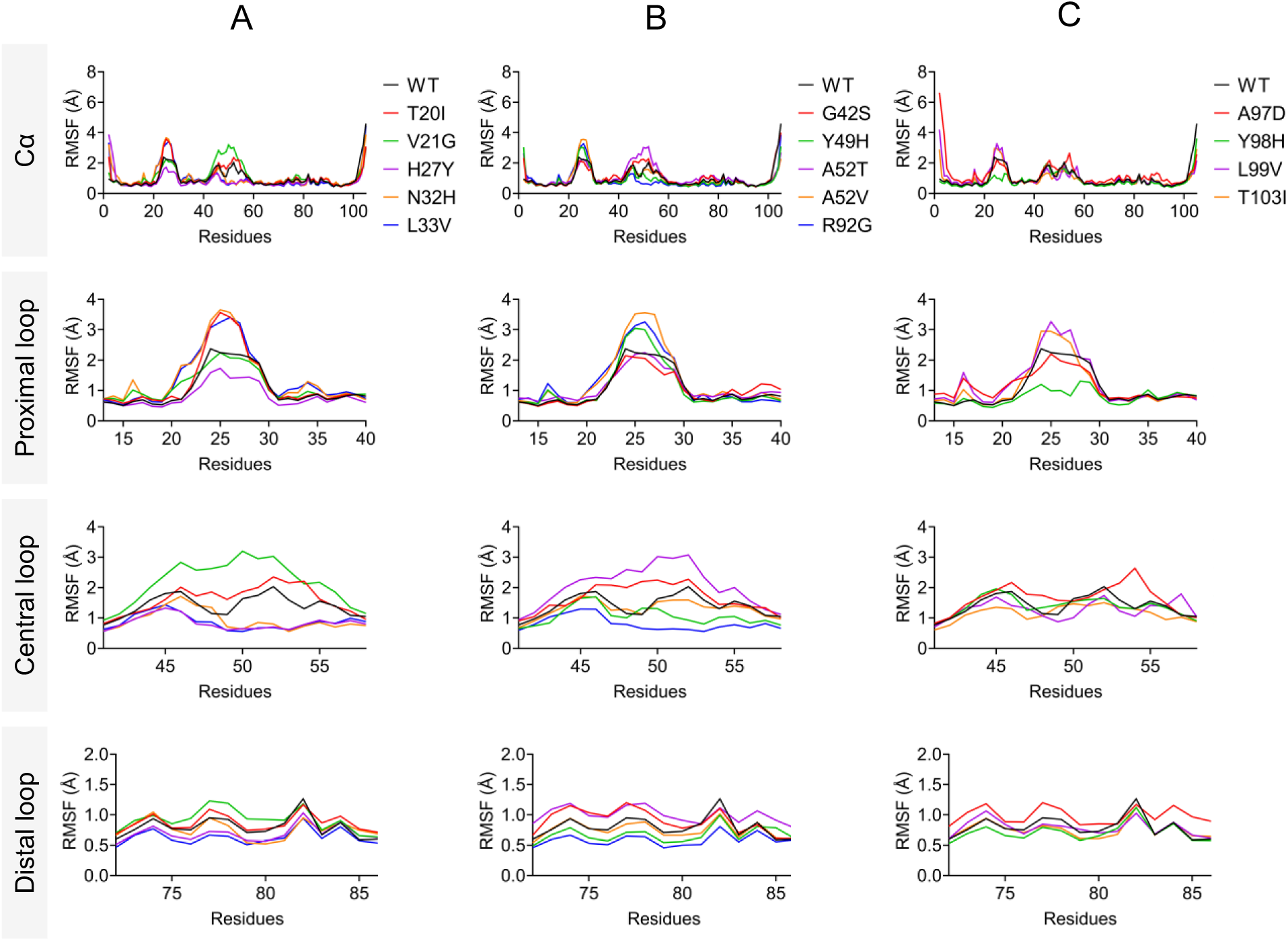
Root mean square fluctuation (RMSF) profiles comparing the structural flexibility of the Cyt-c WT and mutant proteins over the 500 ns MD simulation. The analysis includes the entire protein backbone (Cα atoms) and the proximal, central, and distal Ω-loops. (A) RMSF variations for the WT (black) and mutants T20I (red), V21G (green), H27Y (purple), N32H (orange), and L33V (blue) showing variations in backbone and Ω-loop flexibility. (B) RMSF profiles for the WT (black) and mutants G42S (red), Y49H (green), A52T (purple), A52V (orange), and R92G (blue), highlighting differences in flexibility across the protein and Ω-loops. (C) RMSF differences for the WT (black) and mutants A97D (red), Y98H (green), L99V (purple), and T103I (orange) illustrate distinct flexibility patterns in the backbone and Ω-loops for each variant.

### 3.12 Distance analysis suggests enhanced peroxidase activity of Cyt-c

Here, we analyze if the structural dynamics of the Ω-loops in the hexacoordinated form are associated with the peroxidase activity observed in Cyt-c mutants. For that, we measured the distances between the center of mass of each Ω-loop and the ferric iron in the heme (**Fig. 8 & Supplementary data, Table S8**). The distance evolution between each of the Ω-loop and heme Fe indicates that V21G, H27Y, N32H, G42S, Y49H, A52V, R92G, A97D, and L99V variants cause the movement farther of proximal Ω-loop from the heme iron (**Fig. 8A-C**, Proximal loop-Fe), while T20I, V21G, G42S, A52T, A52V, A97D, Y98H, and L99V increase the distance between central Ω-loop and heme Fe (**Fig. 8A-C**, Central loop-Fe). While the T20I, V21G, N32H, L33V, G42S, Y49H, A52V, L99V, and T103I variants increase the distal Ω-loop-Fe distance (**Fig. 8A-C**, Distal loop-Fe). In short, all the analyzed variants lead to the increased distance of at least one Ω-loop and the heme Fe (**Fig. 8D-F**).

**Fig. 8.**
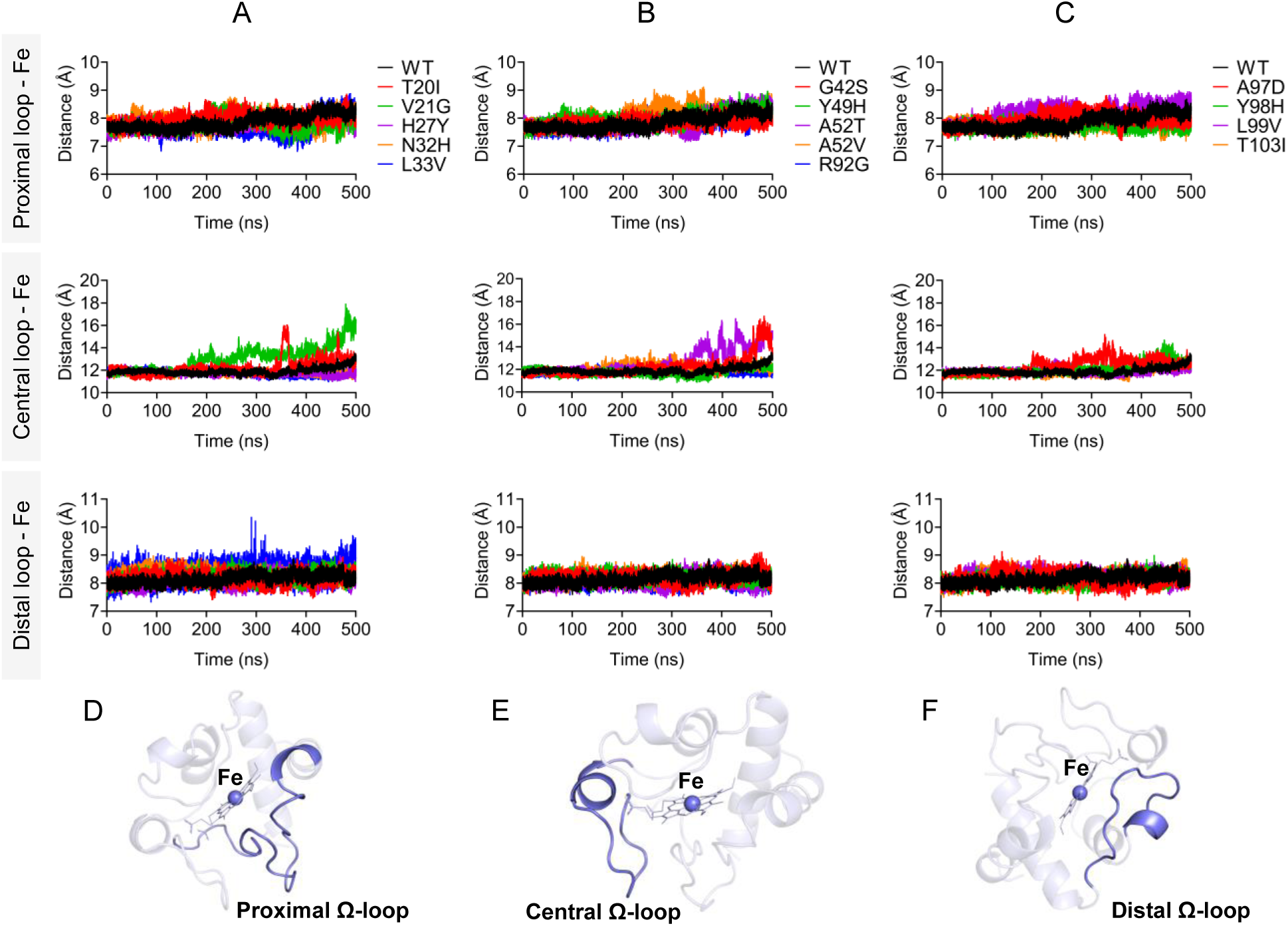
Distance analysis of the center of mass of Ω-loops to the heme Fe for WT and mutant Cyt-c over the 500 ns MD simulation. (A) The distance variations between center of mass of proximal Ω-loop and heme Fe for WT (black) and mutants T20I (red), V21G (green), H27Y (purple), N32H (orange), and L33V (blue). (B) Central Ω-loop (center of mass) to heme Fe distance for the WT (black) and mutants G42S (red), Y49H (green), A52T (purple), A52V (orange), and R92G (blue). (C) Distance from center of mass of distal Ω-loop to heme Fe analyzed for the WT (black) and mutants A97D (red), Y98H (green), L99V (purple), and T103I (orange). Presentation of (D) Proximal, (E), central, and (F) distal Ω-loops along with heme Fe.

In Cyt-c, Tyr68, a well-conserved residue near the heme site, has been shown to play a key role in regulating the structural characteristics of the heme cavity and influencing the peroxidase function of Cyt-c [82,83]. Tyr68 forms an H-bond network with the axial ligand Met81 and evolutionary conserved residues Asn53 and Thr79. This network involving Tyr68 is crucial for the conformational switch, orienting the protein toward the apoptotic pathway [84]. Consequently, distances between these residues were calculated (**Fig. 9 & Supplementary data, Table S9**). The results indicate the weakening of Tyr68 and Asn53 H-bonds in T20I, V21G, L33V, G42S, A52T, A52V, A97D, Y98H, and L99V mutants with increased distance (**Fig. 9A-C**, Tyr68-Asn53), possibly due to the displacement of the central Ω-loop away from the heme Fe (**Fig. 8A-C**, Central loop-Fe). V21G, L33V, G42S, A52T, and L99V variants lead to the loss of the Tyr68 and Met81 H-bond (**Fig. 9A-C**, Tyr68-Met81) mainly due to the rearrangement of distal Ω-loop (**Fig. 8A-C**, Distal loop-Fe). However, the variants do not significantly affect the Tyr68-Thr79 H-bond, except H27Y, L33V, G42S, A52T, A97D where a slight increase in the distances was observed (**Fig. 9F**). This disruption of specific interactions, Tyr68-Asn53 (**Fig. 9D**) and Tyr68-Met81 (**Fig. 9E**) may destabilize the active heme configuration by altering the positions of Asn53 and Met81.

**Fig. 9.**
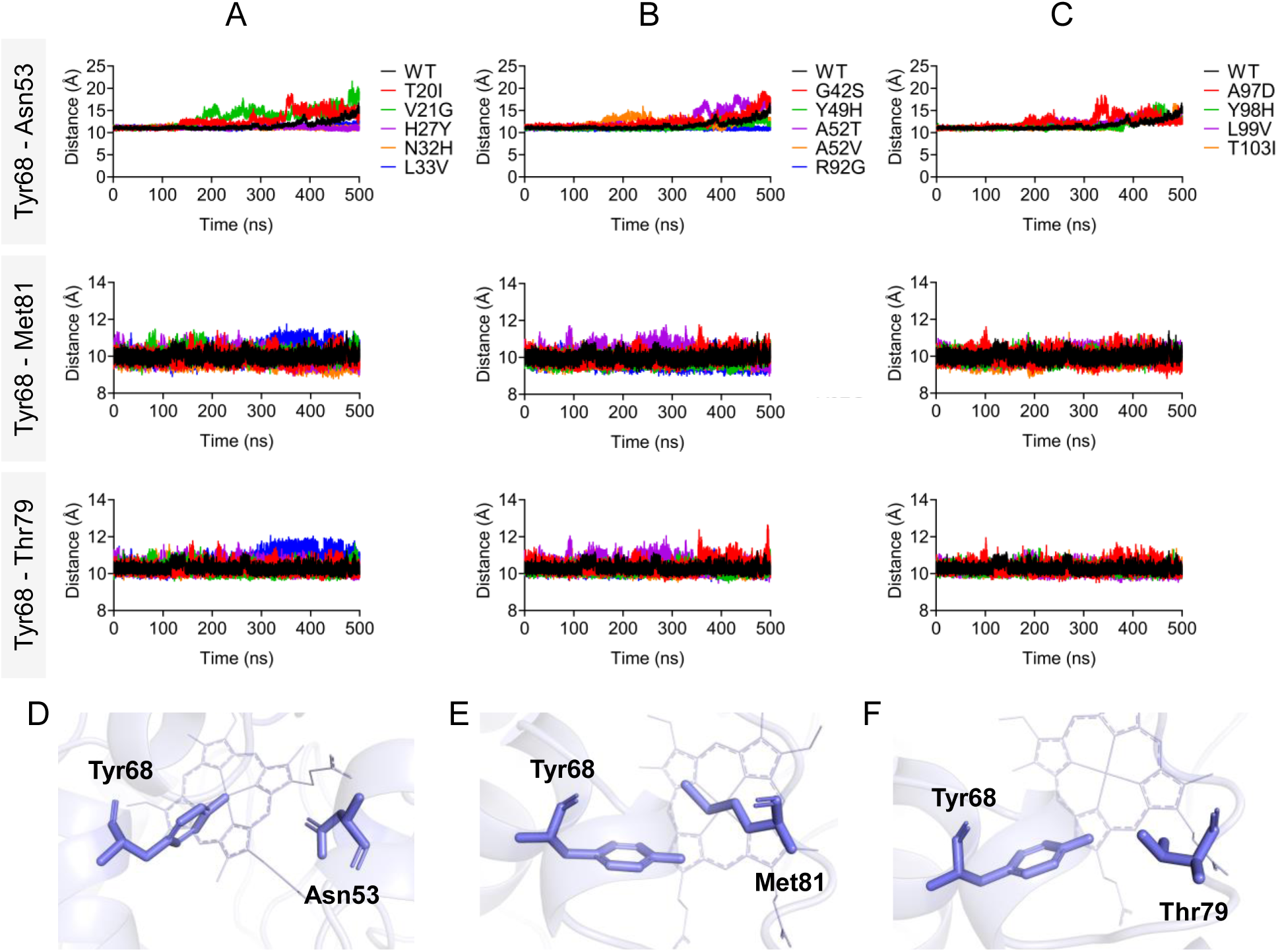
Evolution of distances between H-bonded residues between Tyr68-Asn53, Tyr68-Met81, and Tyr68-Thr79 for WT and mutant Cyt-c over 500 ns MD simulation. (A) The distance variations for WT (black) and mutants T20I (red), V21G (green), H27Y (purple), N32H (orange), and L33V (blue). (B) Distance for the WT (black) and mutants G42S (red), Y49H (green), A52T (purple), A52V (orange), and R92G (blue). (C) Distance analyzed for the WT (black) and mutants A97D (red), Y98H (green), L99V (purple), and T103I (orange). Residues, (D) Tyr68-Asn53, (E) Tyr68-Met81, and (F) Tyr68-Thr79 have been visualized as stick on protein structure. Note that amino acid numbering differs by +1 from that of the PDB file (3ZCF).

We further performed the distance calculations to analyze the opening of specific cavities, since this phenomenon has been associated with the Cyt-c peroxidase activity by regulating the access of substrate to the heme [23,72,85,86]. The opening of cavities analysis was performed following Bortolotti et al. [86], where the opening of the first cavity, for instance, cavity A, was defined by the distance between Ala51–Gly78 Cα while the opening of the second cavity (cavity B) was calculated by the distance Asn32–Ala44 C_α_. Cavity A consists of residues 50–53 on one side and 77-80 on the other; likewise, cavity B comprises residues 32–36 and 42–44. The movement of these cavities is considered to involve relative shifts between the central and distal Ω-loops for Cavity A, and between the proximal and central Ω-loops for Cavity B. It has been proposed that, in yeast iso-1-Cytc, the open state of both cavities facilitates substrate access to the heme crevice [86]. So, it is proposed that the reversible cavity openings permit more water molecules to reach the heme cavity, also facilitating H₂O₂ access to the heme. The analysis of cavity A for all mutant structures during the simulation suggests that T20I, V21G, G42S, A52T, A52V, A97D, and L99V variants cause an increase in Ala51–Gly78 C_α_ distance compared to the WT, leading to the opening of cavity A. The median distance of cavity A for these mutants is in the order V21G > A52V > T20I > L99V > G42S > A52T > A97D (**Fig. 10A-C**, Cavity A). The cavity B analysis indicates that H27Y, L33V, A52T, R92G, A97D, and T103I variants cause the opening of Cavity B when compared with the WT. The median distance for these variants is in the following order: L33V > T103I > H27Y > R92G > A52T > A97D (**Fig. 10A-C**, Cavity B). So, the results suggest that all the variants, except N32H, Y49H, and Y98H, cause an opening of either cavity A or cavity B, when compared to WT (**Fig. 10D & Supplementary data**, **Table S10**). Collectively, all such structural changes could increase the accessibility of the heme catalytic site to small molecules like H₂O₂, thereby enhancing peroxidase activity and facilitating the initiation of the apoptosis pathway.

**Fig. 10.**
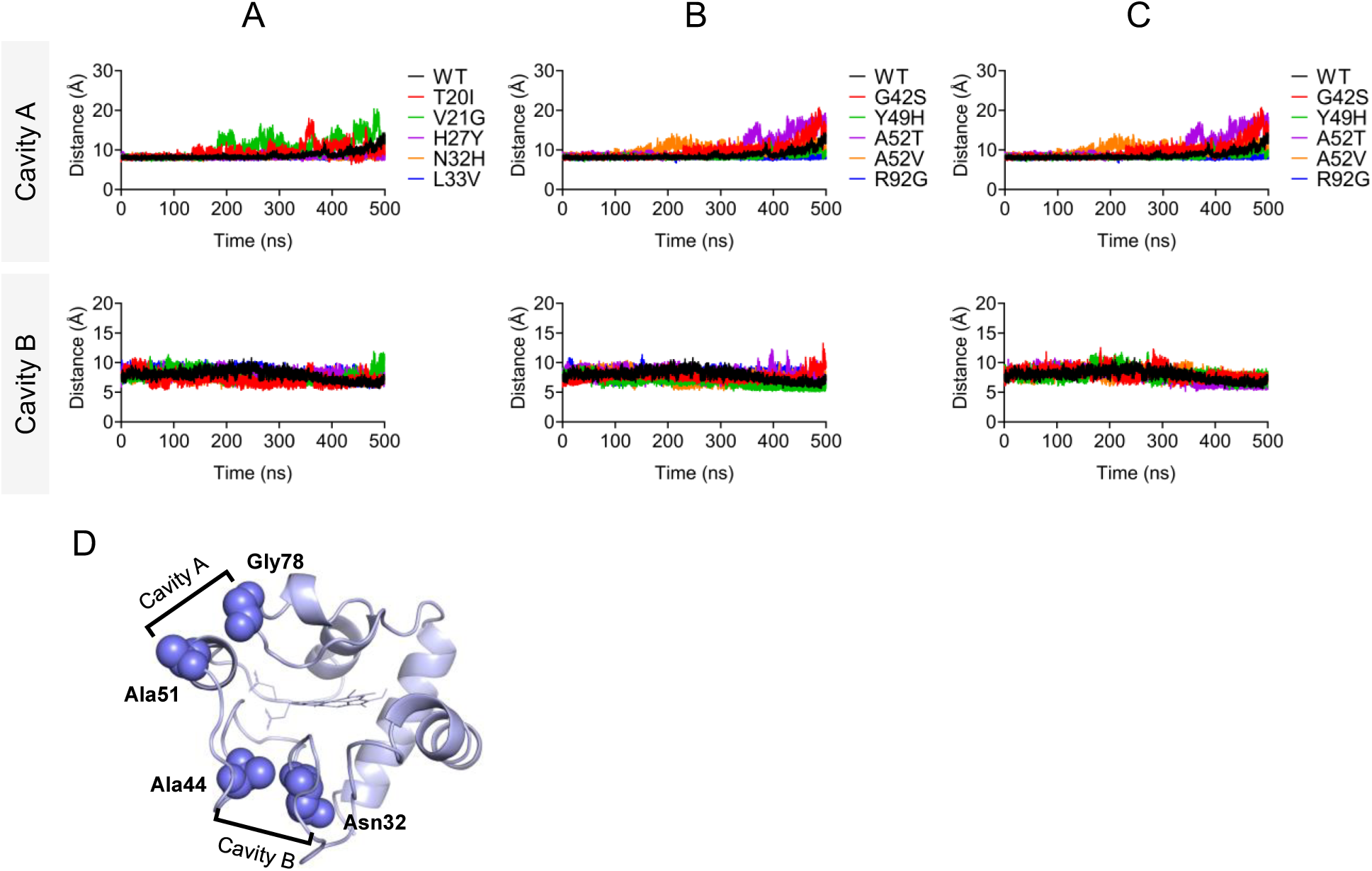
Distance analysis of WT and mutant Cyt-c during a 500 ns MD simulation to evaluate cavity opening dynamics. Distances between Asn51-Gly78 and Asn32-Ala44 were measured to assess the structural changes in cavity A and cavity B, respectively. (A) Comparison of WT (black) and mutants T20I (red), V21G (green), H27Y (purple), N32H (orange), and L33V (blue) for both cavities. (B) Analysis for WT (black) and mutants G42S (red), Y49H (green), A52T (purple), A52V (orange), and R92G (blue) for cavities A and B. (C) Data for WT (black) and mutants A97D (red), Y98H (green), L99V (purple), and T103I (orange) for cavities A and B. (D) Visualization of Asn51-Gly78 and Asn32-Ala44 residues to display the analyzed cavity A and cavity B, respectively on protein structure. Note that amino acid numbering differs by +1 from that of the PDB file (3ZCF).

In native Cyt-c, the hexacoordinate heme iron is axially ligated by the nitrogen of His19 and the sulfur of Met81. Therefore, the distances between the heme iron and the nitrogen (NE2) of His19, as well as the sulfur (SD) of Met81, were calculated for each mutant (**Fig. 11, Supplementary Data, Table S11**). The analysis shows that the V21G, N32H, L33V, R92G, A97D, and L99V variants cause an increase in the distance from His19 to the heme iron (**Fig. 11A-C**, His19-Fe). Similarly, the N32H, L33V, A52T, and R92G variants show an increased distance from Met81 to the heme iron compared to the WT (**Fig. 11A-C**, Met81-Fe). Such displacement (**Fig. 11D**) is associated with peroxidase activity [85,87]. These expanded distances in many mutants lead to major structural alterations, creating an open conformation of the active site that enhances the entry of small molecules like H₂O₂ and other peroxides to the heme center, increasing peroxidase activity and, thereby, apoptosis [82].

**Fig. 11.**
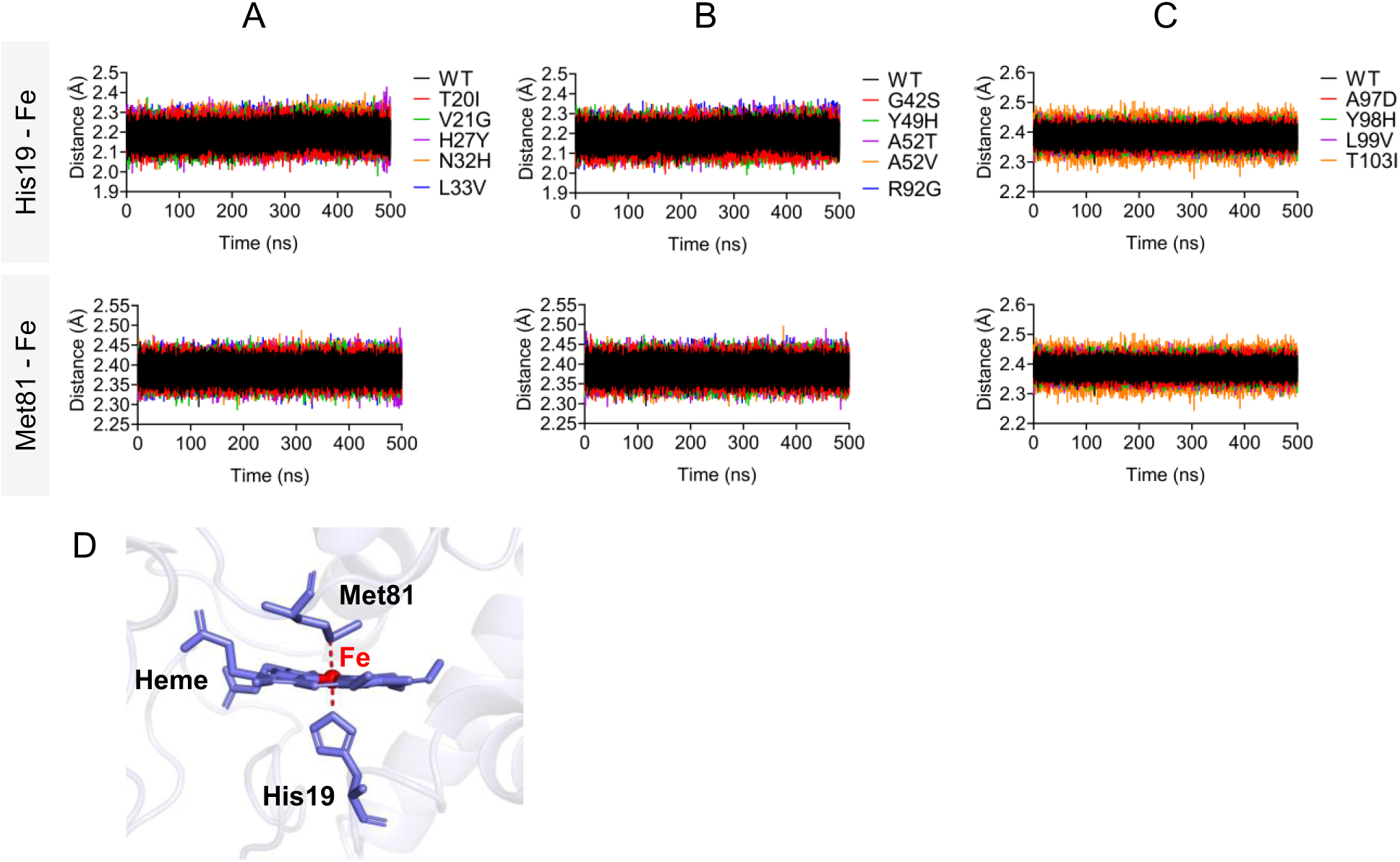
Distance analysis of WT and mutant Cyt-c during a 500 ns MD simulation to evaluate His19 nitrogen (NE2) and Met81 sulfur (SD) axial ligands of hexacoordinate heme iron (Fe). (A) Comparison of WT (black) and mutants T20I (red), V21G (green), H27Y (purple), N32H (orange), and L33V (blue) for His19-Fe and Met81-Fe distances. (B) Analysis for WT (black) and mutants G42S (red), Y49H (green), A52T (purple), A52V (orange), and R92G (blue) for His19(N)-Fe and Met81(S)-Fe distances. (C) Data for WT (black) and mutants A97D (red), Y98H (green), L99V (purple), and T103I (orange) for His19-Fe and Met81-Fe distances.

### 3.13 Normal mode analysis

The functional flexibility and mobility of WT and mutant proteins were assessed using the iMODS server. Deformability analysis reveals notable differences in flexibility across various regions of their structures. Each plot shows peaks that highlight areas of high deformability (**Supplementary data, Fig. S5**). Except for the A52T, all mutants showed lower eigenvalues than the WT, indicating increased flexibility and decreased stability (**Supplementary data, Fig. S6**). These lower eigenvalues suggest that less energy is needed for structural changes, allowing these proteins to alter their conformation more easily. Covariance matrices, used to illustrate correlations in motion between pairs of residues, are shown in maps where red regions denote residues moving in a correlated manner, white indicates uncorrelated movements, and blue represents anti-correlated movements. In the WT, most residues demonstrate strongly correlated movements (shown in red), which indicates coordinated movement. In contrast, most of the variants show an increase in anti-correlated movements (shown in blue), reducing the correlated movements and suggesting that these residues move more independently or in opposite directions. This could lead to increased flexibility and reduced stability. However, Y49H mainly displays uncorrelated movements while A52V exhibits both positive and negative correlations (**Supplementary data, Fig. S7**). The elastic network model provides insights into the stiffness or flexibility of the protein structure, with darker dots representing stronger, stiffer connections (indicating a more rigid structure) and lighter dots suggesting weaker, more flexible connections. The WT maintains a stable structure with a balanced distribution of stiff (dark) and flexible (light) connections. However, variants such as H27Y, N32H, G42S, Y49H, A52V, A97D, and L99V show more flexible connections, as indicated by lighter dots, suggesting they confer increased flexibility to the structure. The rest of the mutants display patterns similar to the WT, indicating only minor changes in flexibility (**Supplementary data, Fig. S8**). These variations affect the dynamic properties of Cyt-c, enhancing either flexibility or stiffness.

## 4 Discussion

The interplay between increased peroxidase activity and apoptosis in Cyt-c mutations presents a complex mechanism contributing to the pathogenesis of THC4. Cyt-c, besides the mitochondrial respiratory chain, is crucial in the intrinsic apoptosis pathway [7,88]. For apoptosis, Cyt-c moves across the outer mitochondrial membrane into the cytosol to activate the caspase cascade and initiate the apoptotic activity. This process could be initiated through several stimuli, including oxidative stress. It is believed that permeabilization of the outer mitochondrial membrane for Cyt-c release to the cytosol is induced by the peroxidase activity of Cyt-c in complex with cardiolipin, a phospholipid found in the mitochondrial inner membrane [89,90].

Post-translational modifications and mutations in Cyt-c can alter its structure and apoptotic function [73,91]. In this study, we propose that Cyt-c variants cause structural destabilization and increased flexibility, specifically through the loss of the intra-protein H-bond network and increased distance between important residues in the heme catalytic site. These changes increase H₂O₂ access, causing oxidative stress and enhancing peroxidase activity, which stimulates apoptosis. This increase in apoptotic activity, particularly within megakaryocytes, the progenitor cells for platelets, could lead to a reduction in platelet count, manifesting as THC4 [23,31]. The precise molecular mechanisms linking these enhanced activities to the clinical phenotype of THC4 remain an area of active investigation, with current hypotheses suggesting a disruption in the balance between platelet production and apoptosis. Our study shows that all variants, except H27Y, A52T, and Y98H, display higher RMSD values for the entire protein during MD simulation, indicating protein destabilization. The formation of the apoptosome depends on the interaction between Cyt-c and Apaf-1. Higher RMSD values are associated with increased apoptosome activation; however, the mechanism underlying increased RMSD could differ between the pathogenic variants [23]. In agreement with our results, a previous study reporting 200 ns simulation for human Cyt-c displayed higher RMSD values for G42S compared with the WT [92]. Similarly, a study of the G42S, Y49H, and A52V structures [23] showed that in a ferric state, all these mutant structures displayed higher RMSD values than those of the WT. This instability in the G42S structure is primarily due to increased RMSF in the central Ω-loop, while our study indicates that G42S increases flexibility in all loops. However, Y49H and A52V are major contributors to the flexibility of the proximal Ω-loop [23], which completely aligns with our findings. Our analysis further revealed that most variants cause protein unfolding and higher flexibility of each Ω-loop. The increased Ω-loop flexibility, specifically the central Ω-loop, also impacts Cyt-c interaction with cardiolipin, facilitating Cyt-c translocation to the cytosol during apoptosis [75]. This flexibility further enhances apoptosome activation, suggesting that Cyt-c and Apaf-1 interactions may follow an “induced fit” mechanism, particularly in variants with greater conformational flexibility [23]. Further, higher RMSF values of loops, specifically in the proximal and central Ω-loops are associated with the increased apoptosome activation [23].

Next, we analyzed how different variants might lead to structural rearrangements in the protein structure that could be associated with higher peroxidase activity of the protein. For that, we performed distance analyses between critical residues involved in enhancing accessibility to the heme. Most variants cause a widening of the heme active site, as indicated by increased distances between each of the Ω-loop and heme Fe. Such movements of the Ω-loops away from the heme also lead to the weakening of the Tyr68-Asn53, and Tyr68-Met81 hydrogen bonds. These residues are highly conserved and are key to the structure, function, folding, and stability of Cyt-c [83,93]. For instance, Tyr68 in the 60-70 helix extends its side chain into the heme pocket and forms hydrogen bonds with Asn53, and Met81 [78], which are important for electron transfer, apoptosis initiation, and heme pocket stability [75,94–98]. Disruption of these Tyr68 hydrogen bonds with Asn53, and Met81 increases caspase activation, impacting peroxidase activity and apoptosis [78,95].

We further evaluated the opening of two cavities, cavity A (Asn51-Gly78) and cavity B (Asn32-Ala44), which result from structural rearrangements caused by various variants. All variants, except N32H, Y49H, and Y98H, lead to the opening of at least one of these cavities. It is suggested that the reversible opening of the cavities facilitates an increased influx of water molecules into the heme crevice, thereby enhancing the accessibility of H_2_O_2_ to the heme, essential for initiating peroxidase activity [2,23,99]. Additionally, we analyzed the distances from the axial ligands His19 nitrogen and Met81 sulfur to the heme iron. The analysis indicates that the V21G, N32H, L33V, R92G, A97D, and L99V variants increase the His19-heme iron distance, while N32H, L33V, A52T, and R92G variants increase the Met81-heme iron distance relative to WT. The weakening of the His19 and Met81 ligation with the heme iron is associated with enhanced peroxidase activity [85,87]. These structural changes, including increased distances in the heme active site and the opening of various cavities, would facilitate the access of small molecules like H₂O₂ to the heme catalytic site, thereby promoting peroxidase activity and apoptosis [78].

We can then wonder how do our predictions correlate with what we know already experimentally from previous clinical and/or biochemical studies. Variants H27Y, G42S, Y49H, A52T, A52V, R92G, and L99V have been previously described in the literature. H27Y, found in a Chinese family with mild THC4 and low platelet counts, can destabilize protein structure and impair function as displayed by bioinformatics analysis [31]. Normally, H27 forms a crucial H-bond with P45 for maintaining heme orientation and α-helices, affecting cardiolipin binding to Cyt-c and apoptosis [62]. Replacing histidine with tyrosine may affect electron transport in the mitochondrial respiratory chain [31]. G42S shows enhanced peroxidase activity and proapoptotic effects by altering heme electronic structure and accelerating electron exchange [13,22,63]. G42S and Y49H variants reduce respiratory levels and increase apoptosis in yeast and mouse models [14]. A52V increases peroxidase activity by destabilizing Cyt-c native conformation. G42S, Y49H, and A52V reduce overall and local stability compared to WT Cyt-c [63], and cause an increase in apoptosome activation and peroxidase activity [23]. Another study suggests that G42S increases apoptosis by altering the central Ω loop and its heme interactions, both in human and mouse Cyt-c [64]. Y49H has been shown to exhibit higher peroxidase activity than the WT, associated with increased flexibility and unfolding of the central and distal loops as demonstrated by NMR. X-ray analysis of Y49H reveals minimal structural differences compared to WT [68]. Additionally, this study finds that G42S displays higher peroxidase activity than WT, though it is lower than that of Y49H [68]. Similarly, another study reports an increased peroxidase activity by G42S, A52V, Y49H [73]. This increase may be attributed to the displacement of all Ω-loops from the heme iron, facilitating the opening of cavity A and increasing the distances between the residues Tyr68 - Asn53, Tyr68 - Met81, and Tyr68 - Thr79 in the G42S mutant. Similarly, the A52V variant causes the movement of all Ω-loops away from heme iron, leading to the opening of cavity A and weakening the interactions between Tyr68 - Asn53. Y49H primarily results in the displacement of both proximal and distal loops away from heme iron. Furthermore, the three variants (G42S, Y49H, and A52V) have been found to affect the stability of the central loop, which is associated with peroxidase activity [63]. This association is supported by significantly higher RMSD values of the central loop as observed in our study.

From a clinical perspective, we know already that patients with G42S and Y49H variants have reduced mean platelet counts but normal platelet volume and function, while H27Y is associated with mild THC4 and low platelet counts [31]. The A52V family shows lower mean platelet counts with a platelet function defect [19]. In our previous research [18], G42S, Y49H, A52T, R92G, and L99V variants have been described in Italian families with low platelet counts, where A52T, and R92G variants showed effects on oxidative growth and respiratory activity in a yeast model. Further, there was a correlation between the spatial region of Cyt-c affected by mutations and the extent of THC4. For instance, patients with mutations (G42S, Y49H, A52T) in the central loop displayed higher platelet counts compared to those with substitutions (R92G, L99V) in the C-terminal near the distal loop of the protein. Notably, the L99V mutation caused a significant decrease in platelet counts, which could be the instability of all loops (with higher RMSD), resulting in all loops moving away from the central heme iron and significant widening of both cavities A and B, as demonstrated in this study. Further, the R92G variant primarily compromises the stability of the central loop, induces movement of the proximal loop away from the heme iron, and leads to the opening of cavity B. In comparison, G42S mainly affects the stability of the central loop, causing an increase in the distance of the loops from the heme iron and the opening of cavity A. Y49H destabilizes the proximal and distal loops, with a slight increase in their distances from the heme iron without causing the opening of any cavities. Lastly, A52T destabilizes and increases the distance of the central loop from heme iron and opens both cavities.

These experimental and clinical data together also validate our predictions of the pathogenic effects caused by all reported mutations, as evidenced by a SIFT score of 0.00 and a SuSPect score greater than 80. However, the SuSPect score for A52T was the lowest, at 75, which is still considered pathogenic.

While providing a powerful tool to interpret the role of mutations and their effect, our study retains nevertheless some inherent limitations. First of all, our analyses allow us to predict the fold stability, that is, how much the protein will retain its folded conformation, but they do not tell us anything about whether a given mutant will also retain the folding pathway of the wild-type protein. Yjis means that, in the cell, the nascent chain of the protein (that is the chain produced by translation at the ribosome) could not even reach the native structure. It is a reasonable inference to assume that, if not significantly affecting the stability of the final folded structure, the protein may fold, but several aspects would still need to be considered before having the certainty of the effects of a mutation (e.g., RNA stability, folding pathway, higher proteolytic susceptibility, etc.). Second, while some mutations might appear not to affect appreciably the fold, they could well have a strong effect on interactions with other cellular partners which cannot be accounted for in our analyses. Another important limitation is that most of the analyses performed in this work do not and cannot take into explicit account the complexity of the *in vivo* environment, often affected by molecular crowding. Finally, it should be mentioned that MD simulations strongly depend on the force field used: while force fields become increasingly more sophisticated, they inherently depend on a number of assumptions and can never be expected to fully capture the complexity of the molecular internal forces. We thus expect that it will be necessary to validate our bioinformatics predictions experimentally.

Future studies will have to employ biochemical and molecular techniques to examine the effects of Cyt-c mutations on peroxidase activity and apoptosis. This process includes introducing mutations via site-directed mutagenesis, expressing and purifying the mutated Cyt-c protein, and measuring peroxidase activity using substrate oxidation assays. Apoptosis can be assessed through caspase activation and/or Cyt-c release assays, and by cell viability tests. Additionally, structural changes can be investigated by X-ray crystallography or NMR spectroscopy, whereas temperature scans followed by circular dichroism and/or differential scanning calorimetry may give a quantitative measurement of the protein stability. These experiments may confirm our computational predictions and enhance our understanding of the role of clinically-relevant Cyt-c mutations in THC4.

In conclusion, this study offers a comprehensive guide to THC4 mutations by systematically evaluating how each Cyt-c variant impacts protein structure and function. The computational analysis identifies key features, such as decreased protein stability, increased loop flexibility, and altered heme active site accessibility that correlate with enhanced peroxidase activity and apoptosis, hallmarks of THC4 pathology. These insights provide predictive markers to access the pathogenic potential of known and novel mutations, guiding future experimental validations and potentially informing diagnostic and therapeutic strategies for THC4.

## Acknowledgments

We acknowledge the CINECA award under the ISCRA initiative, for the availability of high-performance computing resources and support (project HP10BKFH8P).

