## Supplementary data for "A computational study of the fold and stability of cytochrome c with implications for disease"

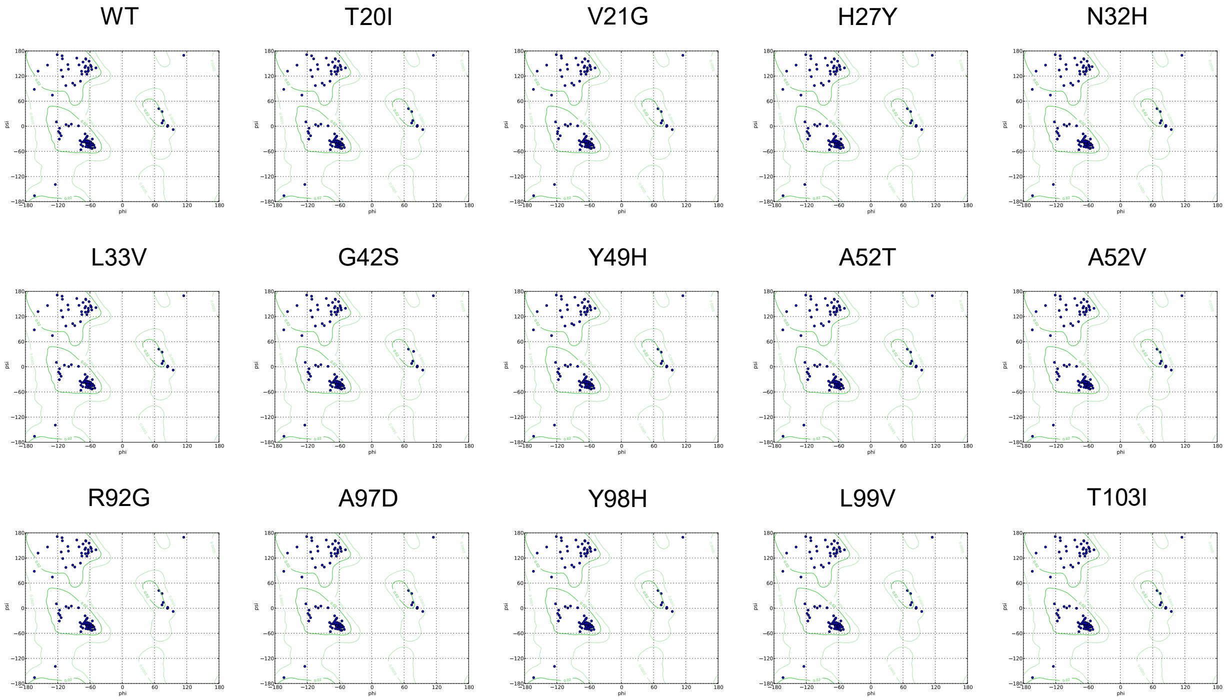


**Fig. S1.** Ramachandran plots for various residues, contrasting WT with mutant residues. Areas inside the green region on each plot denote more favorable regions. Notably, the positions of residues remained unchanged in the plots following mutations (except G42S).

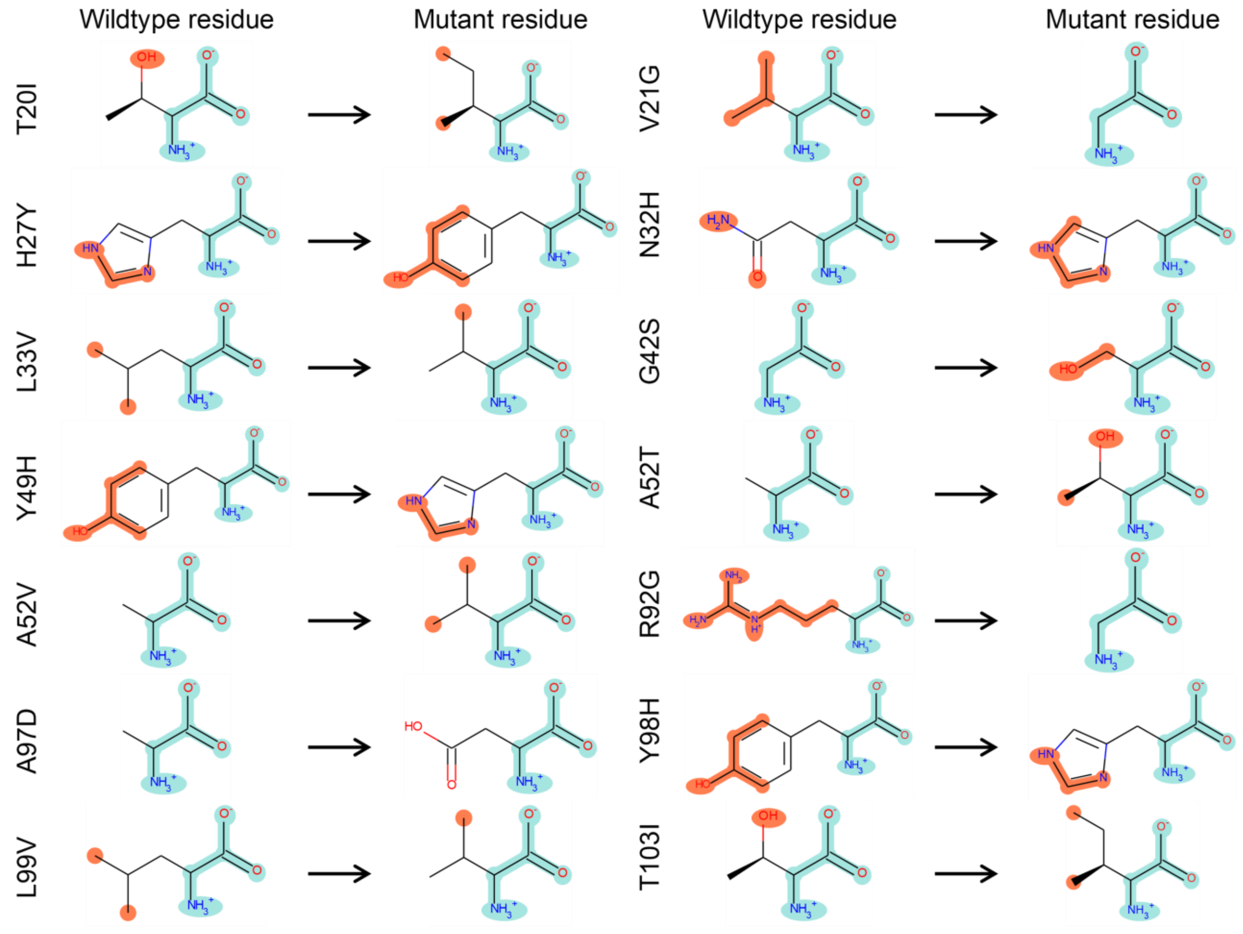


**Fig. S2.** Comparison of two-dimensional structures of wildtype and mutant amino acid residues in Cyt-c. Differing atoms in amino acids are highlighted in red. These representations indicate the impact of clinically important mutations on the affected amino acids.

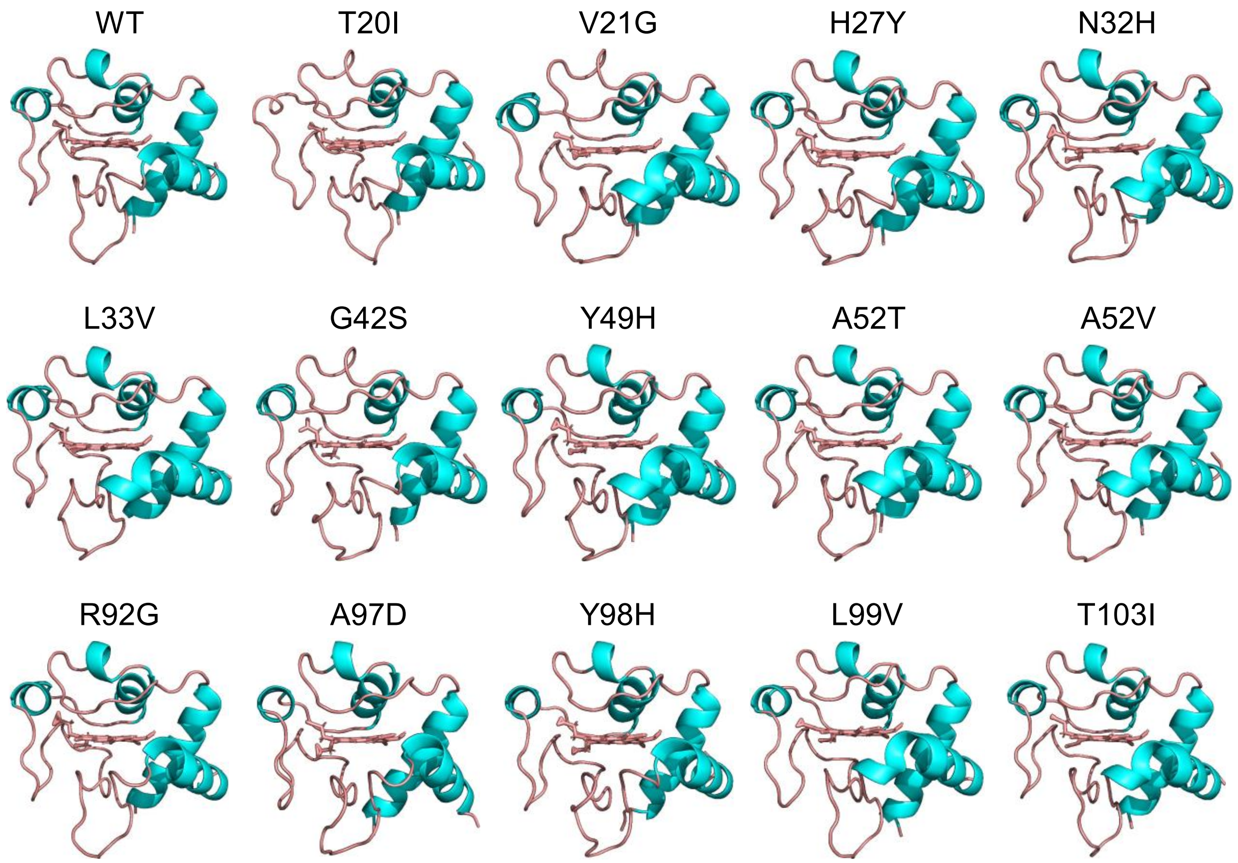


**Fig. S3.** The average Cyt-c structure (WT and mutants) during the 500 ns MD simulation.

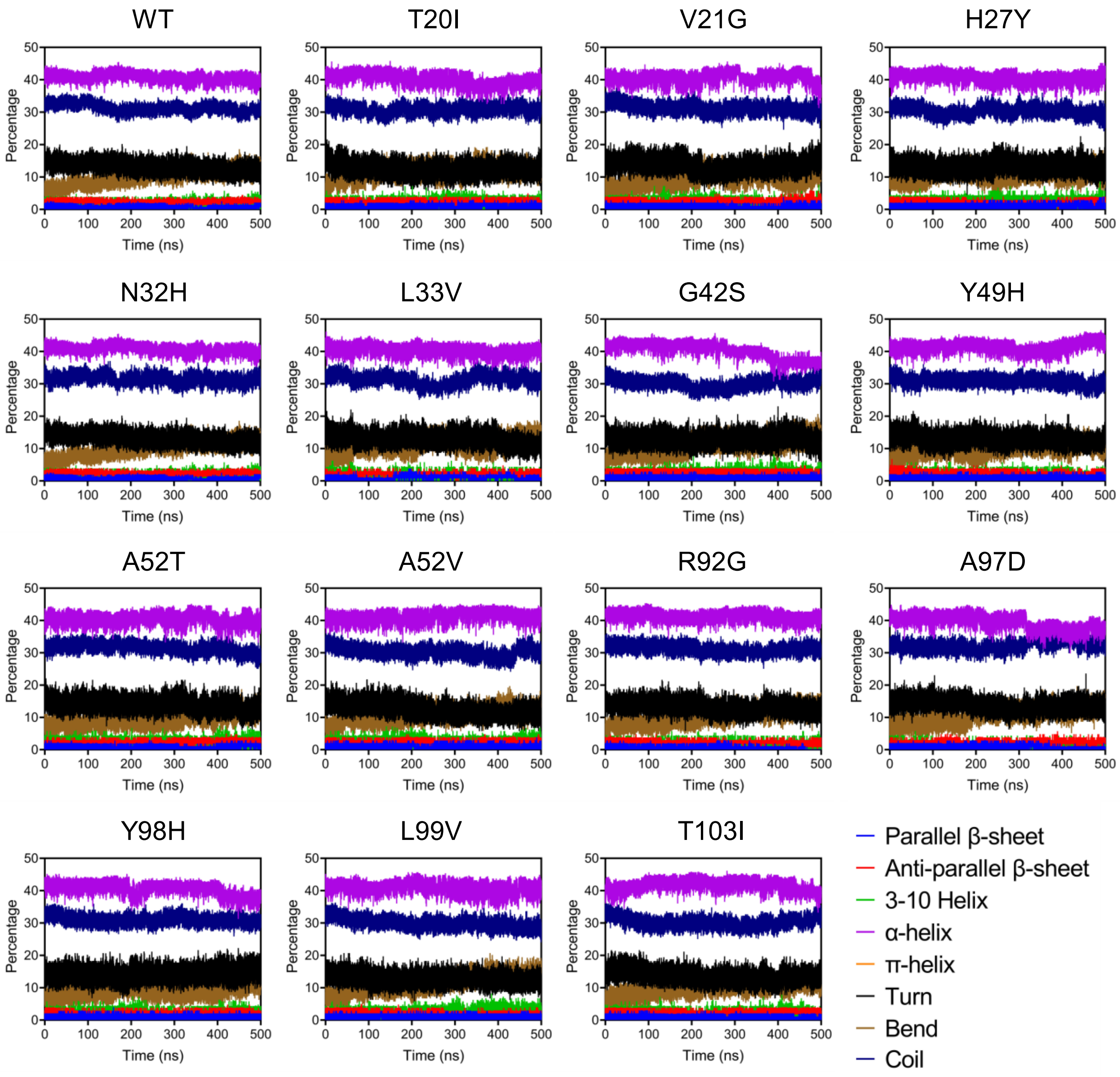


**Fig. S4.** Secondary structure percentages for WT and mutant Cyt-c over the 500 ns MD simulation. The color scheme is shown as: parallel β-sheet (blue), anti-parallel β-sheet (red), 3-10 helix (green), α-helix (purple), π-helix (orange), turn (black), bend (brown), and coil (dark blue).

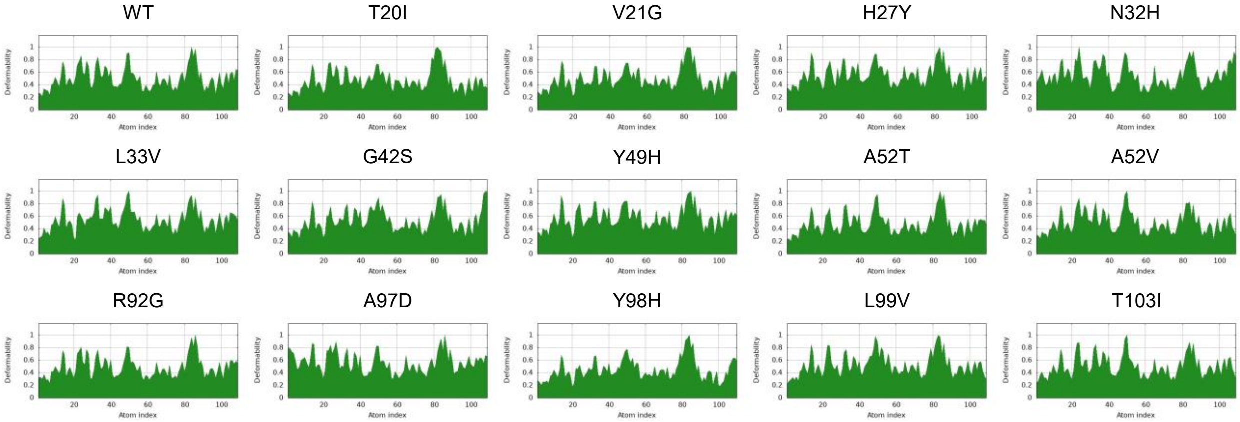


**Fig. S5.** Plots for B-factor/mobility for WT and mutant Cyt-c proteins. Main-chain deformability reflects the ability of a molecule to flex at each residue. Regions with high deformability indicate potential locations of chain 'hinges'.

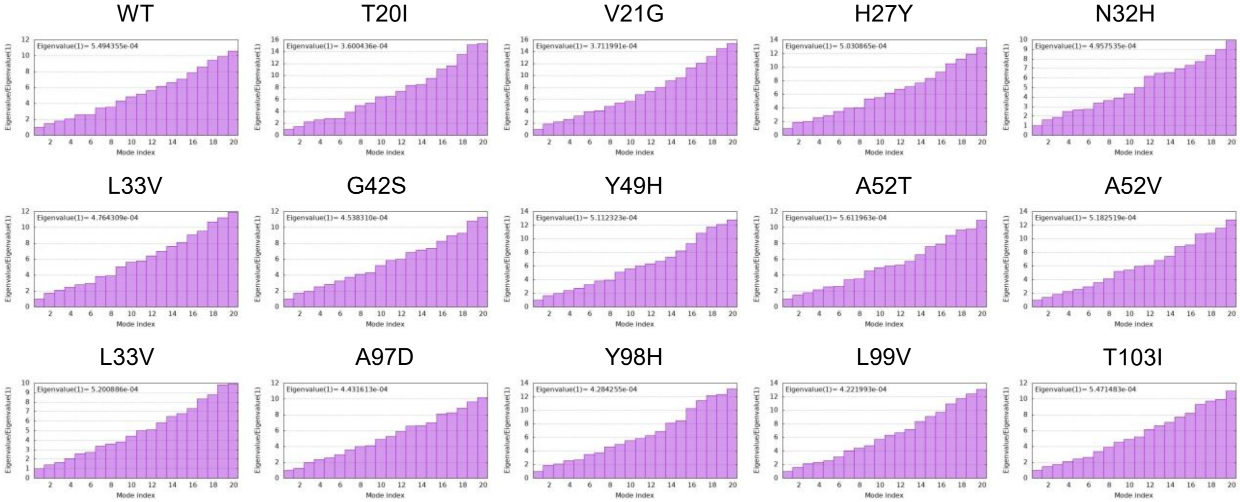


**Fig. S6.** Eigenvalues for WT and mutant Cyt-c proteins. The eigenvalue for each normal mode reflects how rigid the motion is. It is directly linked to the amount of energy needed to deform the structure. A lower eigenvalue means the structure deforms more easily.

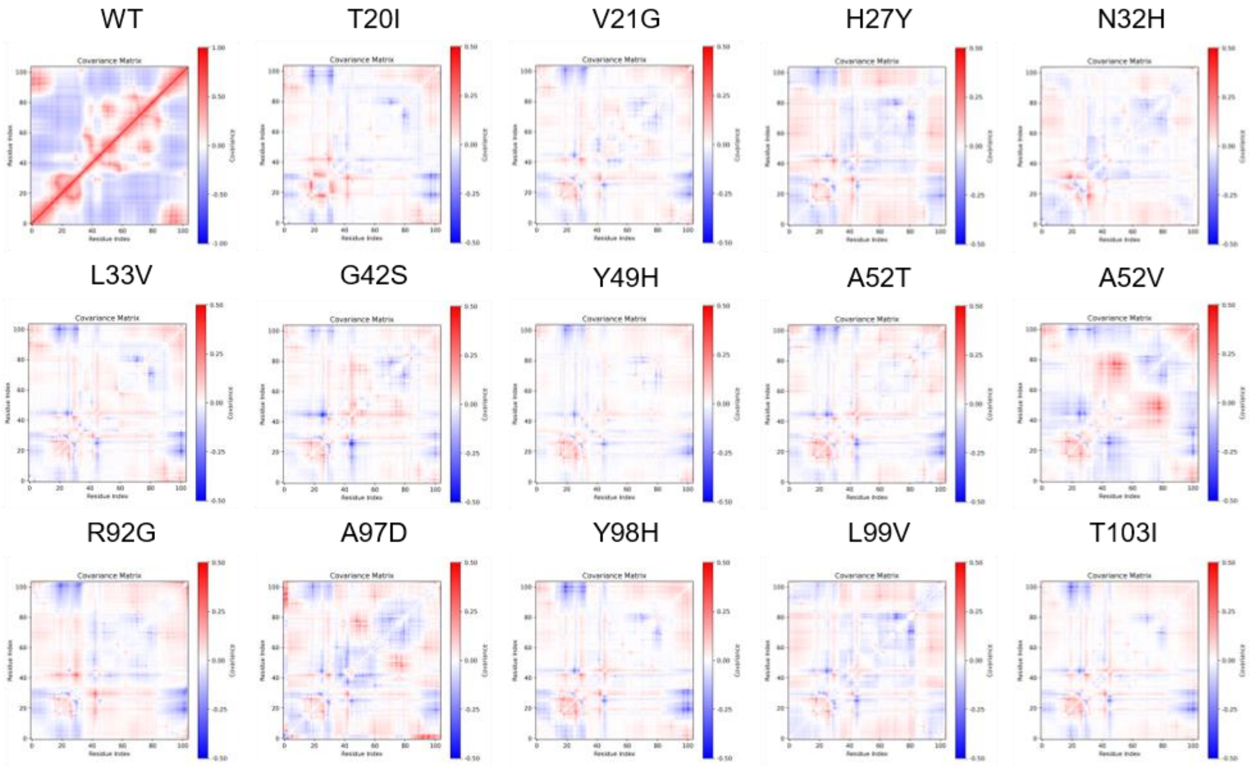


**Fig. S7.** Covariance matrices for WT and mutant proteins. For mutant variants, each matrix represents differences relative to the WT covariance matrix. Red areas indicate correlated residue motions, blue areas indicate anti-correlated motions, and white areas indicate no correlation. Changes in these patterns suggest altered residue interactions and flexibility due to mutations.

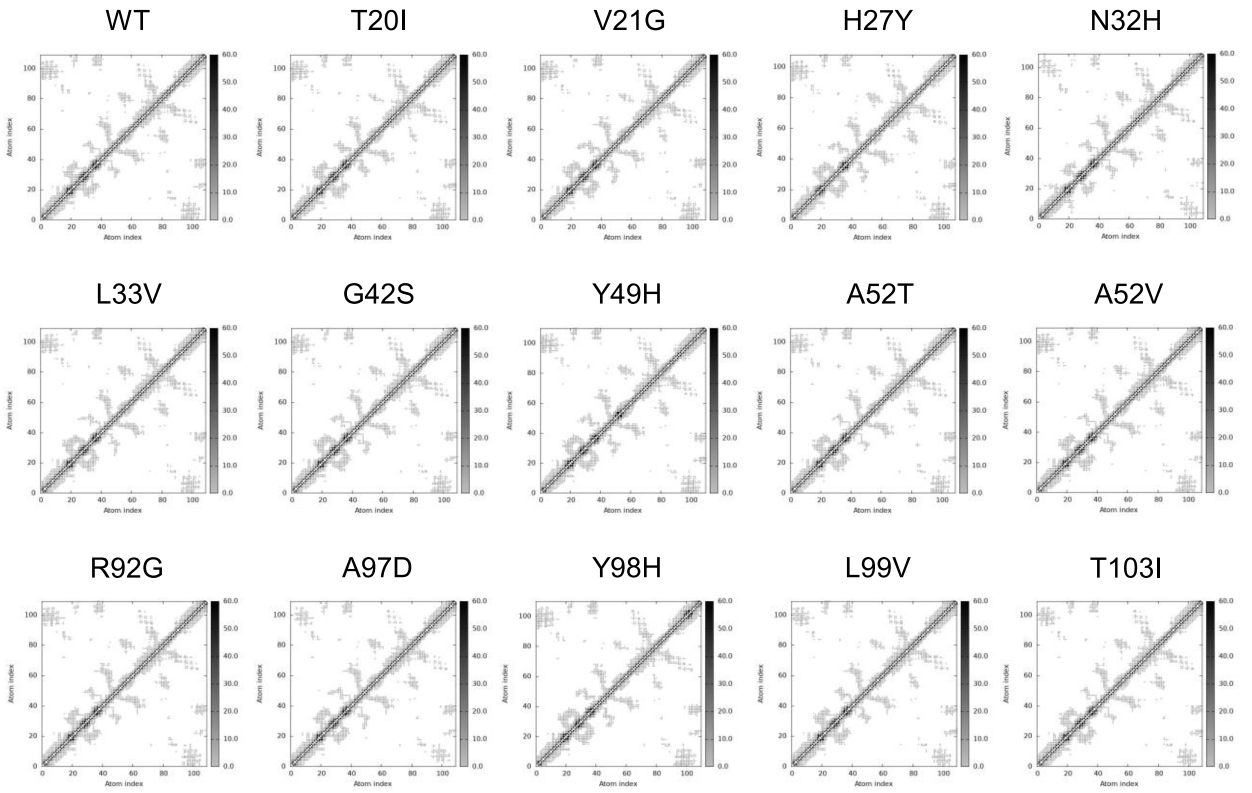


**Fig. S8.** The elastic network for WT and mutant Cyt-c proteins. The elastic network model represents pairs of atoms connected by springs. Each dot on the graph corresponds to a spring between two atoms. The color of the dots reflects the stiffness of the springs, with darker gray indicating stiffer springs and lighter gray representing more flexible ones.

**Table S1.** Multiple sequence alignment of Cyt-c protein across different species.

| ID (Species) | Sequence alignment | |
|---|---|---|
| P99999_[Homo_sapiens] | MGDVEKGKKIFIMKCSQCHTVEKGGKHKTGPNLHGLFGRKTGQAPGYSYTAANKNKGIIW | 60 |
| NP_001157486.1_[Equus_caballus] | MGDVEKGKKIFVQKCAQCHTVEKGGKHKTGPNLHGLFGRKTGQAPGFSYTDANKNKGITW | 60 |
| KAF4003837.1_[Saccharomyces_cerevisiae_iso-1] | AGSAKKGATLFKTRCLQCHTVEKGGPHKVGPNLHGIFGRHSGQAEGYSYTDANIKKNVLW | 60 |
| KAF4000258.1_[Saccharomyces_cerevisiae_iso-2] | PGSAKKGATLFKTRCQQCHTIEEGGPNKVGPNLHGIFGRHSGQVKGYSYTDANINKNVKW | 60 |
| P81459.1_[Thunnus_alalunga] | -GDVAKGKKTFVQKCAQCHTVENGGKHKVGPNLWGLFGRKTGQAEGYSYTDANKSKGIVW | 59 |
| XP_015635265.1_[Oryza_sativa] | VGNAASGKKIFRTKCAQCHTVERGGAHKQGPNLHGLFGRQSGTTPGYAYSTANKNMAVVW | 60 |
| XP_059584883.1_[Alligator_mississippiensis] | MGDVEKGKKIFVQKCAQCHTVEKGGKHKTGPNLHGLIGRKTGQAPGFSYTEANKNKGITW | 60 |
| XP_004553572.1_[Maylandia_zebra] | MGDAEKGKKVFVQKCAQCHTVEQGGKHKVGPNLWGLFGRKTGQAEGFTYTDANKSKGIIW | 60 |
| NP_001385227.1_[Gallus_gallus] | MGDIEKGKKIFVQKCSQCHTVEKGGKHKTGPNLHGLFGRKTGQAEGFSYTDANKNKGITW | 60 |
| AAB86817.3_[Scheffersomyces_stipitis] | KGSEKKGATLFKTRCLQCHTVEEGGPHKVGPNLHGIMGRKSGQAVGYSYTDANKKKGVEW | 60 |
| AAB86850.1_[Fritillaria_agrestis] | PGDFKSGEKIFKTKCAQCHTVDKGAGHKQGPNLNGLFGRQSGTTAGYSYSAANKNKAVNW | 60 |
| XP_014694487.1_[Equus_asinus] | MGDVEKGKKIFVQKCAQCHTVEKGGKHKTGPNLHGLFGRKTGQAPGFSYTDANKNKGITW | 60 |
| C04604_[Cavia_porcellus] | -GDVEKGKKIFVQKCAQCHTVEKGGKHKTGPNLHGLFGRKTGQAAGFSYTDANKNKGITW | 59 |
| XP_042229284.1_[Homarus_americanus] | MGDANKGKKIFVQRCAQCHTVEAGGKHKTGPNLSGLVGRQTGQAPGYVYTDANKAKGITW | 60 |
| NP_001065289.1_[Pan_troglodytes] | MGDVEKGKKIFIMKCSQCHTVEKGGKHKTGPNLHGLFGRKTGQAPGYSYTAANKNKGIIW | 60 |
| XP_058977350.1_[Musca_domestica] | AGDVEKGKKIFVQRCAQCHTVEAGGKHKVGPNLHGLFGRKTGQAAGFAYTDANKAKGITW | 60 |
| XP_030040484.1_[Manduca_sexta] | AGNADNGKKIFVQRCAQCHTVEAGGKHKVGPNLHGFFGRKTGQAPGFSYSDANKAKGITW | 60 |
| XP_008663765.1_[Zea_mays] | GGDAGSGEKIFRTKCAQCHTVERGGAHRQGPNLHGLFGRQSGTTLGYAYSTANKNMAVVW | 60 |
| CAB41053.1_[Schizosaccharomyces_pombe] | PGDEKKGASLFKTRCAQCHTVEKGGANKVGPNLHGVFGRKTGQAEGFSYTEANRDKGITW | 60 |
| XP_006027096.1_[Alligator_sinensis] | MGDVEKGKKIFVQKCAQCHTVEKGGKHKTGPNLHGLIGRKTGQAPGFSYTEANKNKGITW | 60 |
| AAB50255.1_[Aspergillus_nidulans] | PGDSTKGAKLFETRCKQCHTVENGGGHKVGPNLHGLFGRKTGQAGGYAYTDANKQADVTW | 60 |
| XP_004418964.1_[Ceratotherium_simum] | MGDVEKGKKIFVQKCAQCHTVEKGGKHKTGPNLHGLFGRKTGQAPGFSYTDANKNKGITW | 60 |
| XP_054971771.1_[Pan_paniscus] | MGDVEKGKKIFIMKCSQCHTVEKGGKHKTGPNLHGLFGRKTGQAPGYSYTAANKNKGIIW | 60 |
| XP_028701782.1_[Macaca_mulatta] | MGDVEKGKKIFIMKCSQCHTVEKGGKHKTGPNLHGLFGRKTGQAPGYSYTAANKNKGITW | 60 |
| AEP27214.1_[Ateles_paniscus] | MGDVEKGKRIFIQKCSQCHTVEKGGKHKTGPNLHGLFGRKTGQASGFTYTEANKNKGIIW | 60 |
| XP_046526693.1_[Equus_quagga] | MGDVEKGKKIFVQKCAQCHTVEKGGKHKTGPNLHGLFGRKTGQAPGFSYTDANKNKGITW | 60 |
| XP_027828510.2_[_[Ovis_aries] | MGDVEKGKKIFVQKCAQCHTVEKGGKHKTGPNLHGLFGRKTGQAPGFSYTDANKNKGITW | 60 |
| XP_057589467.1_[Hippopotamus_amphibius] | MGDVEKGKKIFVQKCAQCHTVEKGGKHKTGPNLHGLFGRKTGQSPGFSYTDANKNKGITW | 60 |
| XP_017205665.2_[Oryctolagus_cuniculus] | MGDVEKGKKIFVQKCAQCHTVEKGGKHKTGPNLHGLFGRKTGQAVGFSYTDANKNKGITW | 60 |
| AAA21711.1_[Rattus_norvegicus_somatic] | MGDVEKGKKIFVQKCAQCHTVEKGGKHKTGPNLHGLFGRKTGQAAGFSYTDANKNKGITW | 60 |

| | | |
|---|---|---|
| P68098.2_[Lama_guanicoe] | MGDVEKGKKIFVQKCAQCHTVEKGGKHKTGPNLHGLFGRKTGQAVGFSYTDANKNKGITW | 60 |
| NP_001183974.1_[Canis_lupus_familiaris] | MGDVEKGKKIFVQKCAQCHTVEKGGKHKTGPNLHGLFGRKTGQAPGFSYTDANKNKGITW | 60 |
| XP_034876658.1_[Mirounga_leonina] | MGDVEKGKKIFVQKCAQCHTVEKGGKHKTGPNLHGLFGRKTGQAPGFSYTDANKNKGITW | 60 |
| P00013.2_[Miniopterus_schreibersii] | MGDVEKGKKIFVQKCAQCHTVEKGGKHKTGPNLHGLFGRKTGQAPGFSYTDANKNKGITW | 60 |
| P00014.2_[Macropus_giganteus] | MGDVEKGKKIFVQKCAQCHTVEKGGKHKTGPNLNGIFGRKTGQAPGFTYTDANKNKGIIW | 60 |
| CAA25899.1_[Mus_musculus] | MGDVEKGKKIFVQKCAQCHTVEKGGKHKTGPNLHGLFGRKTGQAAGFSYTDANKNKGITW | 60 |
| XP_010711091.1_[Meleagris_gallopavo] | MGDIEKGKKIFVQKCSQCHTVEKGGKHKTGPNLHGLFGRKTGQAEGFSYTDANKNKGITW | 60 |
| P00017.2_[Aptenodytes_patagonicus] | MGDIEKGKKIFVQKCSQCHTVEKGGKHKTGPNLHGIFGRKTGQAEGFSYTDANKNKGITW | 60 |
| XP_025959630.2_[Dromaius_novaehollandiae] | MGDIEKGKKIFVQKCSQCHTVEKGGKHKTGPNLNGLFGRKTGQAEGFSYTDANKNKGITW | 60 |
| XP_068791281.1_[Struthio_camelus] | MGDIEKGKKIFVQKCSQCHTVEKGGKHKTGPNLNGLFGRKTGQAEGFSYTDANKNKGITW | 60 |
| XP_027303674.1_[Anas_platyrhynchos] | MGDVEKGKKIFVQKCSQCHTVEKGGKHKTGPNLHGLFGRKTGQAEGFSYTDANKNKGITW | 60 |
| XP_064907422.1_[Columba_livia] | MGDIEKGKKIFVQKCSQCHTVEKGGKHKTGPNLHGLFGRKTGQAEGFSYTDANKNKGITW | 60 |
| P00022_[Chelydra_serpentina] | MGDVEKGKKIFVQKCAQCHTVEKGGKHKTGPNLNGLIGRKTGQAEGFSYTEANKNKGITW | 60 |
| AFJ49828.1_[Crotalus_adamanteus] | MGDVEKGKKIFSMKCGTCHTVEEGGKHKTGPNLHGLFGRKTGQAVGYSYTAANKNKGIIW | 60 |
| ACO51922.1_[Aquarana_catesbeiana] | MGDVEKGKKIFVQKCAQCHTCEKGGKHKVGPNLYGLIGRKTGQAAGFSYTDANKNKGITW | 60 |
| P00025_[Katsuwonusa_pelamis] | MGDVAKGKKTFVQKCAQCHTVENGGKHKVGPNLWGLFGRKTGQAEGYSYTDANKSKGIVW | 60 |
| XP_042613797.1_[Cyprinus_carpio] | MGDVDKGKKVFVQKCAQCHTVENGGKHKVGPNLWGLFGRKTGQAEGFSYTDANKSKGIVW | 60 |
| P00027_[Squalus_suckleyi] | MGDVEKGKKVFVQKCAQCHTVENGGKHKTGPNLSGLFGRKTGQAQGFSYTDANKSKGITW | 60 |
| P00028_[Entosphenus_tridentatus] | MGDVEKGKKVFVQKCSQCHTVEKAGKHKTGPNLSGLFGRKTGQAPGFSYTDANKSKGIVW | 60 |
| P00029_[Asterias_rubens] | MGQVEKGKKIFVQRCAQCHTVEKAGKHKTGPNLNGILGRKTGQAAGFSYTDANRNKGITW | 60 |
| P00030_[Eisenia_fetida] | AGDVEKGKTIFKQRCAQCHTVDKGGPHKTGPNLHGIFGRATGQAAGFAYTDANKSKGITW | 60 |
| P00031_[Macrobrachium_malcolmsonii] | MGDVEKGKKIFVQRCAQCHSAQANLKHKTGPNLNGLFGRQTGQASGYVYTDANKAKGITW | 60 |
| P00032_[Cornu_aspersum] | -GZAZKGKKIFTQKCLQCHTVEAGGKHKTGPNLSGLFGRKQGQAPGFAYTDANKGKGITW | 59 |
| XP_004536252.1_[Ceratitis_capitata] | TGNAENGKKIFIQKCAQCHPIEKGARHKVGPNLFGVVGRKSGSFADYKYTDANKNKGVTW | 60 |
| P00035_[Haematobia_irritans] | AGDVEKGKKIFVQRCAQCHTVEAGGKHKVGPNLHGLFGRKTGQAAGFAYTNANKAKGITW | 60 |
| XP_023306578.1_[Lucilia_cuprina] | AGDVEKGKKIFVQRCAQCHTVEAGGKHKVGPNLHGLFGRKTGQAPGFAYTDANKAKGITW | 60 |
| P00037_[Samia_cynthia] | AGNAENGKKIFVQRCAQCHTVEAGGKHKVGPNLHGFYGRKTGQAPGFSYSNANKAKGITW | 60 |
| NP_001170961.1_[Apis_mellifera] | AGDPEKGKKIFVQKCAQCHTIESGGKHKVGPNLYGVYGRKTGQAPGYSYTDANKGKGITW | 60 |
| XP_030040484_[Manduca_sexta] | AGNADNGKKIFVQRCAQCHTVEAGGKHKVGPNLHGFFGRKTGQAPGFSYSDANKAKGITW | 60 |
| XP_049857057.1_[Schistocerca_gregaria] | SGDVEKGKKIFVQRCAQCHTVEAGGKHKTGPNLHGLFGRKTGQAPGFSYTDANKSKGITW | 60 |
| P00042_[Wickerhamomyces_anomalus] | KGSEKKGATLFKTRCLQCHTVEKGGPHKVGPNLHGIFGRQSGKAEGYSYTDANIKKAVEW | 60 |
| P00043_[Debaryomyces_hansenii] | KGSEKKGANLFKTRCLQCHTVEEGGPHKVGPNLHGVVGRTSGQAQGFSYTDANKKKGVEW | 60 |

```
P22342_[Euglena_viridis]                         -GDAERGKKLFESRAGQCHSSQKG-VNSTGPALYGVYGRTSGTVPGYAYSNANKNAAIVW    58
P00047_[Thermomyces_lanuginosus]                 PGDASKGANLFKTRCAQCHSVEQGGANKIGPNLHGLFGRKTGSVEGYSYTDANKQAGITW    60
KAK3488365.1_[Neurospora_crassa]                 AGDSKKGANLFKTRCAQCHTLEEGGGNKIGPALHGLFGRKTGSVDGYAYTDANKQKGITW    60
P00049_[Ustilago_sphaerogena]                    DGDAKKGARIFKTRCAQCHTLGAGEPNKVGPNLHGLFGRKSGTVEGFSYTDANKKAGQVW    60
XP_013707149.1_[Brassica_napus]                  PGNSKAGEKIFRTKCAQCHTVDKGAGHKQGPNLNGLFGRQSGTTAGYSYSAANKNKAVEW    60
XP_022994612.1_[Cucurbita_maxima]                PGNSKAGEKIFKTKCAQCHTVDKGAGHKQGPNLNGLFGRQSGTTPGYSYSAANKNMAVIW    60
XP_011095631.1_[Sesamum_indicum]                 AGNKAAGEKIFKTKCAQCHTVEKGAGHKQGPNLNGLFGRQSGTTPGYSYSAANKNMAVIW    60
XP_002528842.1_[Ricinus_communis]                PGDVKAGEKIFKTKCAQCHTVEKGAGHKQGPNLNGLFGRQSGTTAGYSYSAANKNMAVQW    60
P00059_[Abutilon_theophrasti]                    PGBAKAGEKIFKTKCAQCHTVEKGAGHKQGPNLNGLFGRQSGTTPGYSYSAANKNMAVNW    60
P00064_[Allium_porrum]                           PGBZKAGQKIFKLKCAQCHTVEKGAGHKQGPNLNGLFGRQSGTAAGYSYSAANKNMAVVW    60
KAE9612850.1_[Lupinus_albus]                     PGDAKVGEKIFKTKCAQCHTVDKGAGHKQGPNLNGLFGRQSGTTAGYSYSTANKNMAVNW    60
P00069_[Guizotia_abyssinica]                     AGDAKAGEKIFKTKCAZCHTVZKGAGHKQGPNLNGLFGRQSGTTAGYSYSAANKNKAVAW    60
P00072_[Fagopyrum_esculentum]                    PGNIKSGEKIFKTKCAQCHTVEKGAGHKQGPNLNGLFGRQSGTTAGYSYSAANKNKAVTW    60
P00074_[Ginkgo_biloba]                           PGDPKAGEKIFKTKCAZCHTVZKGAGHKQGPNLHGLFGRQSGTTAGYSYSTGNKNKAVNW    60
AAA28437.1_[Drosophila_melanogaster]             AGDVEKGKKLFVQRCAQCHTVEAGGKHKVGPNLHGLIGRKTGQAAGFAYTDANKAKGITW    60
NP_036972.1_[Rattus_norvegicus_testis-specific]  MGDAEAGKKIFIQKCAQCHTVEKGGKHKTGPNLWGLFGRKTGQAPGFSYTDANKNKGVIW    60
P12831_[Sarcophaga_peregrina]                    AGDVEKGKKIFVQRCAQCHTVEAGGKHKVGPNLHGLFGRKTGQAPGFAYTDANKAKGITW    60
CAB16954.1_[Chlamydomonas_reinhardtii]           AGDLARGEKIFKTKCAQCHVAEKGGGHKQGPNLGGLFGRVSGTAAGFAYSKANKEAAVTW    60
P19681.2_[Schwanniomyces_occidentalis]           KGSEKKDANLFKTRCLQCHTVEKGGPHKVGPNLHGIFGRKSGQAAGYSYTDANKKKGVEW    60
P21665_[Varanus_varius]                          MGDVEKGKKIFVQKCSQCHTVEKGGKHKTGPNLHQLFGRKTGEAEGFSYTAANKNKGITW    60
XDT51687.1_[Nakaseomyces_glabratus]              -MSEKKGATLFKTRCLQCHTVEKGGPNKVGPNLHGIFGRKSGQAAGYSYTDANIKKNVTW    59
AAB50255_[Aspergillus_nidulans]                  PGDSTKGAKLFETRCKQCHTVENGGGHKVGPNLHGLFGRKTGQAGGYAYTDANKQADVTW    60
KAL1579288.1_[Candida_albicans]                  KGSEKKGATLFKTRCLQCHTVEKGGPHKVGPNLHGVFGRKSGLAEGYSYTDANKKKGVEW    60
P56205_[Aspergillus_niger]                       PGDSAKGAKLFQTRCAQCHTVEAGGPHKVGPNLHGLFGRKTGQSEGYAYTDANKQAGVTW    60
NP_001039526.1_[Bos_taurus]                      MGDVEKGKKIFVQKCAQCHTVEKGGKHKTGPNLHGLFGRKTGQAPGFSYTDANKNKGITW    60
NP_001123442.1_[Sus_scrofa]                      MGDVEKGKKIFVQKCAQCHTVEKGGKHKTGPNLHGLFGRKTGQAPGFSYTDANKNKGITW    60
XP_068381675.1_[Eschrichtius_robustus]           MCDVEKGKKIFVQKCAQCHTVEKGGKHKTGPNLHGLFGRKTGQAVGFSYTDANKNKGITW    60
XP_031306477.1_[Camelus_dromedarius]             MGDVEKGKKIFVQKCAQCHTVEKGGKHKTGPNLHGLFGRKTGQAVGFSYTDANKNKGITW    60
P12831.2_[Sarcophaga_peregrina]                  AGDVEKGKKIFVQRCAQCHTVEAGGKHKVGPNLHGLFGRKTGQAPGFAYTDANKAKGITW    60
NP_001413453.1_[Helianthus_annuus]               AGNPTTGEKIFKTKCAQCHTVEKGAGHKQGPNLNGLFGRQSGTTAGYSYSAGNKNKAVIW    60
AAB72175.1_[Arabidopsis_thaliana]                PGNAKAGEKIFRTKCAQCHTVEAGAGHKQGPNLNGLFGRQSGTTAGYSYSAANKNKAVEW    60
XP_044353760.1_[Triticum_aestivum]               PGNPAAGEKIFKTKCAQCHTVDKGAGHKQGPNLNGLFGRQSGTTAGYSYSTANKSMAVVW    60
```

```
P00075_[Ulva_intestinalis]                        PGBPAKGAKIFKAKCAZCHTVBAGAGHKQGPNLNGAFGRTSGTAAGFSYSAABKBKTADW    60
XP_060957730.1_[Cannabis_sativa]                  PGNSKTGEKIFRTKCAQCHTVDKGAGHKQGPNLNGLFGRQSGTTAGYSYSAANKNMAVTW    60
P00077_[Strigomonas_oncopelti]                    PGDAAKGEKIFKGRAAQCHTGAKGGANGVGPNLFGIVNRHSGTVEGFAYSKANADSGVVW    60
P00067_[Tropaeolum_majus]                         AGDNKAGDKIFKNKCAQCHTVDKGAGHKQGPNLNGLFGRQSGTTAGYSYSAANKNKAVLW    60
XP_021848354.1_[Spinacia_oleracea]                PGNKDVGAKIFKTKCAQCHTVDQGAGHKQGPNLNGLFGRQSGTAASYSYSAANKNKAVIW    60
P00062_[Sambucus_nigra]                           PGNPKAGEKIFKTKCNQCHTVDKGAGHKQGPNLNGLFGRQSGTTAGYSYSAANKNMAVNW    60
NP_001363126.1_[Vigna_radiata]                    PGNSKSGEKIFKTKCAQCHTVDKGAGHKQGPNLNGLFGRQSGTTAGYSYSTANKNMAVIW    60
P00071_[Pastinaca_sativa]                         PGDKDVGGKIFKTKCAZCHTVZLGAGHKQGPNLNGLFGRQSGTTAGYSYSAANKNKAVLW    60
P00066_[Nigella_damascena]                        AGBSASGEKIFKTKCAZCHTVBZGAGHKZGPNLHGLFGRQSGTVAGYSYSAANKNKAVNW    60
P00058_[Gossypium_barbadense]                     PGBAKAGEKIFKTKCAQCHTVDKGAGHKQGPNLNGLFGRQSGTTAGYSYSAANKNMAVQW    60
XP_014694487_[Equus_asinus]                       MGDVEKGKKIFVQKCAQCHTVEKGGKHKTGPNLHGLFGRKTGQAPGFSYTDANKNKGITW    60
XP_058977350_[Musca_domestica]                    AGDVEKGKKIFVQRCAQCHTVEAGGKHKVGPNLHGLFGRKTGQAAGFAYTDANKAKGITW    60
P00043.3_[Debaryomyces_hansenii]                  KGSEKKGANLFKTRCLQCHTVEEGGPHKVGPNLHGVVGRTSGQAQGFSYTDANKKKGVEW    60
P19974_[Caenorhabditis_elegans]                   AGDYEKGKKVYKQRCLQCHVVDST-ATKTGPTLHGVIGRTSGTVSGFDYSAANKNKGVVW    59
P00078_[Crithidia_fasciculata]                    PGDAARGEKLFKGRAAQCHTANQGGANGVGPNLYGLVGRHSGTIEGYAYSKANAESGVVW    60
P00076_[Euglena_gracilis]                         -GDAERGKKLFESRAAQCHSAQKG-VNSTGPSLWGVYGRTSGSVPGYAYSNANKNAAIVW    58
P00041_[Pichia_kudriavzevii]                      QGSAKKGATLFKTRCAQCHTIEAGGPHKVGPNLHGIFSRHSGQAEGYSYTDANKRAGVEW    60
CAA43224.1_[Kluyveromyces_lactis]                 KGSEKKGATLFKTRCLQCHTVEAGGPHKVGPNLHGVFGRHSGKASGYSYTDANIKKNVLW    60
Q41346_[Stellaria_longipes]                       EGDAKKGANLFKTRCAQCHTLGEGEGNKIGPNLHGLFGRHTGSVEGFSYTDANKAKGIEW    60
                                                        .   :  :.  **          ** *    .*  *   .: *: .       *

P99999_[Homo_sapiens]                             GEDTLMEYLENPKKYIPGTKMIFVGIKKKEERADLIAYLKKATNE    105
NP_001157486.1_[Equus_caballus]                   KEETLMEYLENPKKYIPGTKMIFAGIKKKTEREDLIAYLKKATN-    104
KAF4003837.1_[Saccharomyces_cerevisiae_iso-1]     DENNMSEYLTNPKKYIPGTKMAFGGLKKEKDRNDLITYLKKAC--    103
KAF4000258.1_[Saccharomyces_cerevisiae_iso-2]     DEDSMSEYLTNPKKYIPGTKMAFAGLKKEKDRNDLITYMTKAAK-    104
P81459.1_[Thunnus_alalunga]                       NNDTLMEYLENPKKYIPGTKMIFAGIKKKGERQDLVAYLKSATS-    103
XP_015635265.1_[Oryza_sativa]                     EEGTLYDYLLNPKKYIPGTKMVFPGLKKPQERTDLIAYLKESTA-    104
XP_059584883.1_[Alligator_mississippiensis]       GEETLMEYLENPKKYIPGTKMIFAGIKKKPERADLIAYLKEATAN    105
XP_004553572.1_[Maylandia_zebra]                  GEDTLMEYLENPKKYIPGTKMIFAGIKKKGERQDLIAYLKSATS-    104
NP_001385227.1_[Gallus_gallus]                    GEDTLMEYLENPKKYIPGTKMIFAGIKKKSERVDLIAYLKDATSK    105
AAB86817.3_[Scheffersomyces_stipitis]             SEQTMSDYLENPKKYIPGTKMAFGGLKKPKDRNDLVTYLASATK-    104
AAB86850.1_[Fritillaria_agrestis]                 DENTLYDYLLNPKKYIPGTKMVFPGLKKPQDRADLIAYLKEATSS    105
XP_014694487.1_[Equus_asinus]                     KEETLMEYLENPKKYIPGTKMIFAGIKKKTEREDLIAYLKKATNE    105
```

```
C04604_[Cavia_porcellus]                    GEDTLMEYLENPKKYIPGTKMIFAGIKKKGERADLIAYLKKATNE    104
XP_042229284.1_[Homarus_americanus]         NTETLDVYLTNPKKYIPGTKMVFAGLKKKNERADLIAYLEESTK-    104
NP_001065289.1_[Pan_troglodytes]            GEDTLMEYLENPKKYIPGTKMIFVGIKKKEERADLIAYLKKATNE    105
XP_058977350.1_[Musca_domestica]            NEDTLFEYLENPKKYIPGTKMIFAGLKKPNERGDLIAYLKSATK-    104
XP_030040484.1_[Manduca_sexta]              NEDTLFEYLENPKKYIPGTKMVFAGLKKANERADLIAYLKQATK-    104
XP_008663765.1_[Zea_mays]                   EEGTLYDYLFNPKKYIPGTKMVFPGLKKPKERTDLIAYLKESTA-    104
CAB41053.1_[Schizosaccharomyces_pombe]      DEETLFAYLENPKKYIPGTKMAFAGFKKPADRNNVITYLKKATSE    105
XP_006027096.1_[Alligator_sinensis]         GEETLMEYLENPKKYIPGTKMIFAGIKKKPERADLIAYLKEATAN    105
AAB50255.1_[Aspergillus_nidulans]           DENSLFKYLENPKKYIPGTKMAFGGLKKTKERNDLITYLKESTA-    104
XP_004418964.1_[Ceratotherium_simum]        KEETLMEYLENPKKYIPGTKMIFAGIKKKAEREDLIAYLKKATNE    105
XP_054971771.1_[Pan_paniscus]               GEDTLMEYLENPKKYIPGTKMIFVGIKKKEERADLIAYLKKATNE    105
XP_028701782.1_[Macaca_mulatta]             GEDTLMEYLENPKKYIPGTKMIFVGIKKKEERADLIAYLKKATNE    105
AEP27214.1_[Ateles_paniscus]                GEDTLMEYLENPKKYIPGTKMIFVGIKKKGEREDLIAYLKKATNE    105
XP_046526693.1_[Equus_quagga]               KEETLMEYLENPKKYIPGTKMIFAGIKKKTEREDLIAYLKKATNE    105
XP_027828510.2_[_[Ovis_aries]               GEETLMEYLENPKKYIPGTKMIFAGIKKKGEGG-LDSLSQKATNE    104
XP_057589467.1_[Hippopotamus_amphibius]     GEETLMEYLENPKKYIPGTKMIFAGIKKKGERADLIAYLKQATNE    105
XP_017205665.2_[Oryctolagus_cuniculus]      GEDTLMEYLENPKKYIPGTKMIFVGIKKKNERADLIAYLKKATNE    105
AAA21711.1_[Rattus_norvegicus_somatic]      GEDTLMEYLENPKKYIPGTKMIFAGIKKKGERADLIAYLKKATNE    105
P68098.2_[Lama_guanicoe]                    GEETLMEYLENPKKYIPGTKMIFAGIKKKGERADLIAYLKKATNE    105
NP_001183974.1_[Canis_lupus_familiaris]     GEETLMEYLENPKKYIPGTKMIFAGIKKTGERADLIAYLKKATKE    105
XP_034876658.1_[Mirounga_leonina]           GEETLMEYLENPKKYIPGTKMIFAGIKKTGERADLIAYLKIATKE    105
P00013.2_[Miniopterus_schreibersii]         GEATLMEYLENPKKYIPGTKMIFAGIKKSAERADLIAYLKKATKE    105
P00014.2_[Macropus_giganteus]               GEDTLMEYLENPKKYIPGTKMIFAGIKKKGERADLIAYLKKATNE    105
CAA25899.1_[Mus_musculus]                   GEDTLMEYLENPKKYIPGTKMIFAGIKKKGERADLIAYLKKATNE    105
XP_010711091.1_[Meleagris_gallopavo]        GEDTLMEYLENPKKYIPGTKMIFAGIKKKSERVDLIAYLKDATSK    105
P00017.2_[Aptenodytes_patagonicus]          GEDTLMEYLENPKKYIPGTKMIFAGIKKKSERADLIAYLKDATSK    105
XP_025959630.2_[Dromaius_novaehollandiae]   GEDTLMEYLENPKKYIPGTKMIFAGIKKKSERADLIAYLKDATSK    105
XP_068791281.1_[Struthio_camelus]           GEDTLMEYLENPKKYIPGTKMIFAGIKKKSERADLIAYLKDATSK    105
XP_027303674.1_[Anas_platyrhynchos]         GEDTLMEYLENPKKYIPGTKMIFAGIKKKSERADLIAYLKDATAK    105
XP_064907422.1_[Columba_livia]              GEDTLMEYLENPKKYIPGTKMIFAGIKKKAERADLIAYLKQATAK    105
P00022_[Chelydra_serpentina]                GEETLMEYLENPKKYIPGTKMIFAGIKKKAERADLIAYLKDATSK    105
AFJ49828.1_[Crotalus_adamanteus]            GDDTLMEYLENPKKYIPGTKMVFTGLKSKKERTDLIAYLKEATAK    105
```

| | | |
|---|---|---|
| ACO51922.1_[Aquarana_catesbeiana] | NEDTLMEYLENPKKYIPGTKMIFAGIKKQGERKDLIAYLKSACSC | 105 |
| P00025_[Katsuwonusa_pelamis] | NENTLMEYLENPKKYIPGTKMIFAGIKKKGERQDLVAYLKSATS- | 104 |
| XP_042613797.1_[Cyprinus_carpio] | SEETLMEYLENPKKYIPGTKMIFAGIKKKGERADLIAYLKSATS- | 104 |
| P00027_[Squalus_suckleyi] | QQETLRIYLENPKKYIPGTKMIFAGLKKKSERQDLIAYLKKTAAS | 105 |
| P00028_[Entosphenus_tridentatus] | NQETLFVYLENPKKYIPGTKMIFAGIKKEGERKDLIAYLKKSTSE | 105 |
| P00029_[Asterias_rubens] | KNETLFEYLENPKKYIPGTKMVFAGLKKQKERQDLIAYLEAATK- | 104 |
| P00030_[Eisenia_fetida] | TKDTLYEYLENPKKYIPGTKMVFAGLKNEKQRANLIAYLEQETK- | 104 |
| P00031_[Macrobrachium_malcolmsonii] | QADTLDVYLENPKKYIPGTKMVFAGLKKANERADLIAYLKQATNL | 105 |
| P00032_[Cornu_aspersum] | KNQTLFEYLENPKKYIPGTKMVFAGLKBZTERVHLIAYLZZATKK | 104 |
| XP_004536252.1_[Ceratitis_capitata] | TDDALFEYLENPKKFIPGTKMVFAGLKKAAERADIIAYLKTAK-- | 103 |
| P00035_[Haematobia_irritans] | QDDTLFEYLENPKKYIPGTKMIFAGLKKPNERGDLIAYLKSATK- | 104 |
| XP_023306578.1_[Lucilia_cuprina] | NEDTLFEYLENPKKYIPGTKMIFAGLKKPNERGDLIAYLKSATK- | 104 |
| P00037_[Samia_cynthia] | GDDTLFEYLENPKKYIPGTKMVFAGLKKANERADLIAYLKESTK- | 104 |
| NP_001170961.1_[Apis_mellifera] | NKETLFEYLENPKKYIPGTKMVFAGLKKPQERADLIAYIEQASK- | 104 |
| XP_030040484_[Manduca_sexta] | NEDTLFEYLENPKKYIPGTKMVFAGLKKANERADLIAYLKQATK- | 104 |
| XP_049857057.1_[Schistocerca_gregaria] | NEDTLFIYLENPKKYIPGTKMVFAGLKKPQERADLIAYLKESTK- | 104 |
| P00042_[Wickerhamomyces_anomalus] | SEQTMSDYLENPKKYIPGTKMAFGGLKKEKDRNDLVTYLANATK- | 104 |
| P00043_[Debaryomyces_hansenii] | SEQNLSDYLENPKKYIPGTKMAFGGLKKAKDRNDLISYLVKATK- | 104 |
| P22342_[Euglena_viridis] | EDESLNKFLENPKKYVPGTKMAFAGIKAKKDRLDIIAYMKTLKD- | 102 |
| P00047_[Thermomyces_lanuginosus] | NEDTLFEYLENPKKFIPGTKMAFGGLKKNKDRNDLITYLKEATK- | 104 |
| KAK3488365.1_[Neurospora_crassa] | DENTLFEYLENPKKYIPGTKMAFGGLKKDKDRNDIITYMKEATA- | 104 |
| P00049_[Ustilago_sphaerogena] | EEETFLEYLENPKKYIPGTKMAFGGLKKEKDRNDLVTYLREETK- | 104 |
| XP_013707149.1_[Brassica_napus] | EEKTLYDYLLNPKKYIPGTKMVFPGLKKPQDRADLIAYLKEATA- | 104 |
| XP_022994612.1_[Cucurbita_maxima] | EEKTLYDYLLNPKKYIPGTKMVFPGLKKPQDRADLIAYLKESTA- | 104 |
| XP_011095631.1_[Sesamum_indicum] | EEKTLYDYLLNPKKYIPGTKMVFPGLKKPQERADLIAYLKESTAN | 105 |
| XP_002528842.1_[Ricinus_communis] | GENTLYDYLLNPKKYIPGTKMVFPGLKKPQDRADLIAYLKQATA- | 104 |
| P00059_[Abutilon_theophrasti] | GENTLYDYLLNPKKYIPGTKMVFPGLKKPQDRADLIAYLKZSTA- | 104 |
| P00064_[Allium_porrum] | ZZBTLYDYLLNPKKYIPGTKMVFPGLKKPQDRADLIAYLKESTA- | 104 |
| KAE9612850.1_[Lupinus_albus] | EEKTLYDYLLNPKKYIPGTKMVFPGLKKPQDRADLIAYLKESTAQ | 105 |
| P00069_[Guizotia_abyssinica] | ZZBSLYDYLLNPKKYIPGTKMVFPGLKKPZZRADLIAYLKASTA- | 104 |
| P00072_[Fagopyrum_esculentum] | GEDTLYEYLLNPKKYIPGTKMVFPGLKKPQERADLIAYLKBSTZ- | 104 |
| P00074_[Ginkgo_biloba] | GZZTLYEYLLNPKKYIPGTKMVFPGLKKPZZRADLISYLKQATSQ | 105 |

| | | |
|---|---|---|
| AAA28437.1_[Drosophila_melanogaster] | NEDTLFEYLENPKKYIPGTKMIFAGLKKPNERGDLIAYLKSATK- | 104 |
| NP_036972.1_[Rattus_norvegicus_testis-specific] | TEETLMEYLENPKKYIPGTKMIFAGIKKKSEREDLIQYLKEATSS | 105 |
| P12831_[Sarcophaga_peregrina] | NEDTLFEYLENPKKYIPGTKMIFAGLKKPNERGDLIAYLKSATK- | 104 |
| CAB16954.1_[Chlamydomonas_reinhardtii] | GESTLYEYLLNPKKYMPGNKMVFAGLKKPEERADLIAYLKQATA- | 104 |
| P19681.2_[Schwanniomyces_occidentalis] | TEQTMSDYLENPKKYIPGTKMAFGGLKKPKDRNDLITYLANATK- | 104 |
| P21665_[Varanus_varius] | GEDTLFEYLENPKKYIPGTKMIFAGIKKKTERDDLIAYLKEATAK | 105 |
| XDT51687.1_[Nakaseomyces_glabratus] | DEDNMSDYLTNPKKYIPGTKMAFGGLKKEKDRKDLIAYLKKATSD | 104 |
| AAB50255_[Aspergillus_nidulans] | DENSLFKYLENPKKYIPGTKMAFGGLKKTKERNDLITYLKESTA- | 104 |
| KAL1579288.1_[Candida_albicans] | TEQTMSDYLENPKKYIPGTKMAFGGLKKPKDRNDLVTYLKKATS- | 104 |
| P56205_[Aspergillus_niger] | DENTLFSYLENPKKFIPGTKMAFGGLKKGKERNDLITYLKESTA- | 104 |
| NP_001039526.1_[Bos_taurus] | GEETLMEYLENPKKYIPGTKMIFAGIKKKGEREDLIAYLKKATNE | 105 |
| NP_001123442.1_[Sus_scrofa] | GEETLMEYLENPKKYIPGTKMIFAGIKKKGEREDLIAYLKKATNE | 105 |
| XP_068381675.1_[Eschrichtius_robustus] | GEETLMEYLENPKKYIPGTKMIFAGIKKKGERADLIAYLKKATNE | 105 |
| XP_031306477.1_[Camelus_dromedarius] | GEETLMEYLENPKKYIPGTKMIFAGIKKKGERADLIAYLKKATNE | 105 |
| P12831.2_[Sarcophaga_peregrina] | NEDTLFEYLENPKKYIPGTKMIFAGLKKPNERGDLIAYLKSATK- | 104 |
| NP_001413453.1_[Helianthus_annuus] | EENTLYDYLLNPKKYIPGTKMVFPGLKKPQERADLIAYLKTSTA- | 104 |
| AAB72175.1_[Arabidopsis_thaliana] | EEKALYDYLLNPKKYIPGTKMVFPGLKKPQDRADLIAYLKESTAP | 105 |
| XP_044353760.1_[Triticum_aestivum] | EEKTLYDYLLNPKKYIPGTKMVFPGLKKPQERADLIAYLKSATA- | 104 |
| P00075_[Ulva_intestinalis] | BZBTLYDYLLNPKKYIPGTKMVFAGLKKPZBRADLIAFLKDATA- | 104 |
| XP_060957730.1_[Cannabis_sativa] | EEKTLYDYLLNPKKYIPGTKMVFPGLKKPQDRADLIAYLKESTA- | 104 |
| P00077_[Strigomonas_oncopelti] | TPEVLDVYLENPKKFMPGTKMSFAGIKKPQERADLIAYLENLK-- | 103 |
| P00067_[Tropaeolum_majus] | ZZATLYDYLLNPKKYIPGTKMVFPGLKKPQDRADLIAYLKESTA- | 104 |
| XP_021848354.1_[Spinacia_oleracea] | SEDTLYEYLLNPKKYIPGTKMVFPGLKKPQDRADLIAYLKDSTQ- | 104 |
| P00062_[Sambucus_nigra] | EEKTLYDYLLNPKKYIPGTKMVFPGLKKPQDRADLIAYLKQSTA- | 104 |
| NP_001363126.1_[Vigna_radiata] | EEKTLYDYLLNPKKYIPGTKMVFPGLKKPQDRADLIAYLKESTA- | 104 |
| P00071_[Pastinaca_sativa] | ABBTLYDYLLNPKKYIPGTKMVFPGLKKPQDRADLIAYLKHATA- | 104 |
| P00066_[Nigella_damascena] | EEKTLYDYLLNPKKYIPGTKMVFPGLKKPZZRABLLAYLKESTA- | 104 |
| P00058_[Gossypium_barbadense] | GENTLYDYLLNPKKYIPGTKMVFPGLKKPQDRADLIAYLKZSTA- | 104 |
| XP_014694487_[Equus_asinus] | KEETLMEYLENPKKYIPGTKMIFAGIKKKTEREDLIAYLKKATNE | 105 |
| XP_058977350_[Musca_domestica] | NEDTLFEYLENPKKYIPGTKMIFAGLKKPNERGDLIAYLKSATK- | 104 |
| P00043.3_[Debaryomyces_hansenii] | SEQNLSDYLENPKKYIPGTKMAFGGLKKAKDRNDLISYLVKATK- | 104 |
| P19974_[Caenorhabditis_elegans] | TKETLFEYLLNPKKYIPGTKMVFAGLKKADERADLIKYIEVESAK | 104 |

```
P00078_[Crithidia_fasciculata]          TPDVLDVYLENPKKFMPGTKMSFAGMKKPQERADVIAYLETLKG-      104
P00076_[Euglena_gracilis]               EEETLHKFLENPKKYVPGTKMAFAGIKAKKDRQDIIAYMKTLKD-      102
P00041_[Pichia_kudriavzevii]            AEPTMSDYLENPKKYIPGTKMAFGGLKKAKDRNDLVTYMLEASK-      104
CAA43224.1_[Kluyveromyces_lactis]       DEQTMSDYLENPKKYIPGTKMAFGGLKKEKDRNDIVTYMLKACK-      104
Q41346_[Stellaria_longipes]             NKDTLFEYLENPKKYIPGTKMAFGGLKKDKDRNDLITFLQDSTK-      104
                                            :  :* ****::**.** * *:*       :
```

**s**

**Table S2.** Molecular effects of Cyt-c mutations with their MutPred2 score, probability score and p-values.

| Mutation (MutPred2 score) | Molecular effect | Probability | p-values (<= 0.05) |
|---|---|---|---|
| T20I (0.719) | Loss of Intrinsic disorder | 0.37 | 0.04 |
| | Altered Metal binding | 0.33 | 0.01 |
| | Altered Ordered interface | 0.28 | 0.03 |
| | Gain of Strand | 0.27 | 0.02 |
| | Gain of Catalytic site at C18 | 0.16 | 0.02 |
| | Altered DNA binding | 0.14 | 0.05 |
| | Gain of Pyrrolidone carboxylic acid at Q17 | 0.06 | 0.03 |
| V21G (0.858) | Gain of Intrinsic disorder | 0.48 | 2.50E-03 |
| | Altered Metal binding | 0.3 | 0.01 |
| | Gain of B-factor | 0.3 | 2.80E-03 |
| | Altered Stability | 0.26 | 7.40E-03 |
| | Loss of Catalytic site at C18 | 0.18 | 0.02 |
| | Loss of Pyrrolidone carboxylic acid at Q17 | 0.05 | 0.04 |
| H27Y (0.863) | Altered Ordered interface | 0.45 | 1.90E-04 |
| | Loss of Intrinsic disorder | 0.38 | 0.03 |
| | Gain of Acetylation at K26 | 0.3 | 4.00E-03 |
| | Loss of B-factor | 0.26 | 0.03 |
| | Altered Metal binding | 0.24 | 0.04 |
| | Gain of Methylation at K28 | 0.09 | 0.04 |
| | Gain of Catalytic site at K28 | 0.08 | 0.05 |
| N32H (0.926) | Altered Ordered interface | 0.29 | 0.03 |
| | Altered Metal binding | 0.27 | 0.02 |
| | Loss of Catalytic site at K28 | 0.08 | 0.05 |
| L33V (0.863) | Altered Ordered interface | 0.28 | 0.04 |
| | Gain of Allosteric site at H34 | 0.19 | 0.04 |
| | Gain of Catalytic site at N32 | 0.08 | 0.05 |
| G42S (0.917) | Altered Ordered interface | 0.28 | 0.03 |
| | Loss of Relative solvent accessibility | 0.25 | 0.04 |
| | Loss of Methylation at K40 | 0.16 | 0.01 |
| | Gain of Catalytic site at R39 | 0.15 | 0.02 |
| | Altered Transmembrane protein | 0.14 | 0.02 |
| | Altered Metal binding | 0.12 | 0.03 |
| | Loss of Pyrrolidone carboxylic acid at Q43 | 0.07 | 0.02 |
| Y49H (0.924) | Altered Ordered interface | 0.47 | 1.30E-04 |
| | Altered Disordered interface | 0.29 | 0.03 |
| | Loss of Phosphorylation at Y49 | 0.28 | 0.02 |
| | Loss of Relative solvent accessibility | 0.26 | 0.03 |
| | Altered Metal binding | 0.25 | 2.90E-03 |

| | | | |
|---|---|---|---|
| | Loss of Allosteric site at Y49 | 0.25 | 0.02 |
| | Altered Transmembrane protein | 0.21 | 4.10E-03 |
| | Loss of Methylation at K54 | 0.2 | 6.70E-03 |
| | Altered DNA binding | 0.19 | 0.02 |
| | Gain of Catalytic site at K54 | 0.1 | 0.04 |
| A52T (0.906) | Loss of Relative solvent accessibility | 0.27 | 0.02 |
| | Altered Metal binding | 0.25 | 2.30E-03 |
| | Altered Transmembrane protein | 0.24 | 1.50E-03 |
| | Altered Ordered interface | 0.24 | 0.03 |
| | Loss of Allosteric site at Y49 | 0.24 | 0.02 |
| | Gain of Acetylation at K56 | 0.2 | 0.04 |
| | Loss of Methylation at K54 | 0.2 | 6.60E-03 |
| | Altered DNA binding | 0.16 | 0.04 |
| | Gain of Catalytic site at K56 | 0.1 | 0.04 |
| A52V (0.911) | Gain of Allosteric site at Y49 | 0.32 | 9.30E-04 |
| | Altered Ordered interface | 0.29 | 0.02 |
| | Gain of Relative solvent accessibility | 0.28 | 0.02 |
| | Altered Metal binding | 0.25 | 2.40E-03 |
| | Altered Transmembrane protein | 0.24 | 1.70E-03 |
| | Gain of Acetylation at K56 | 0.21 | 0.03 |
| | Loss of Methylation at K54 | 0.2 | 6.20E-03 |
| | Altered DNA binding | 0.19 | 0.02 |
| | Gain of Catalytic site at K56 | 0.1 | 0.04 |
| R92G (0.888) | Altered Coiled coil | 0.46 | 8.10E-03 |
| | Altered Metal binding | 0.39 | 9.00E-03 |
| | Loss of Allosteric site at R92 | 0.31 | 3.70E-03 |
| | Gain of Methylation at K87 | 0.31 | 1.10E-04 |
| | Altered Disordered interface | 0.3 | 0.02 |
| | Loss of Relative solvent accessibility | 0.3 | 0.01 |
| | Altered DNA binding | 0.29 | 3.80E-03 |
| | Altered Ordered interface | 0.28 | 0.04 |
| | Gain of B-factor | 0.26 | 0.02 |
| | Gain of Acetylation at K87 | 0.26 | 1.00E-02 |
| | Loss of SUMOylation at K89 | 0.23 | 0.02 |
| A97D (0.947) | Altered Coiled coil | 0.37 | 0.01 |
| | Gain of Relative solvent accessibility | 0.34 | 1.90E-03 |
| | Altered Disordered interface | 0.31 | 0.02 |
| | Gain of Acetylation at K100 | 0.28 | 5.40E-03 |
| | Altered Ordered interface | 0.27 | 0.04 |
| | Loss of Methylation at K100 | 0.26 | 1.70E-03 |
| | Gain of Phosphorylation at Y98 | 0.26 | 0.03 |
| | Loss of Allosteric site at R92 | 0.24 | 0.02 |
| | Altered Metal binding | 0.23 | 0.02 |

| | | | |
|---|---|---|---|
| | Altered DNA binding | 0.19 | 0.02 |
| | Gain of Ubiquitylation at K101 | 0.16 | 0.04 |
| Y98H (0.908) | Altered Coiled coil | 0.75 | 1.60E-03 |
| | Altered Ordered interface | 0.44 | 4.40E-04 |
| | Altered Disordered interface | 0.41 | 3.70E-03 |
| | Altered Metal binding | 0.33 | 3.90E-03 |
| | Gain of Intrinsic disorder | 0.3 | 0.04 |
| | Loss of Relative solvent accessibility | 0.28 | 0.02 |
| | Loss of Acetylation at K100 | 0.28 | 5.90E-03 |
| | Loss of Allosteric site at Y98 | 0.26 | 0.01 |
| | Loss of Methylation at K100 | 0.26 | 1.70E-03 |
| | Loss of Phosphorylation at Y98 | 0.26 | 0.03 |
| | Altered Stability | 0.19 | 0.01 |
| | Altered DNA binding | 0.15 | 0.04 |
| | Loss of Ubiquitylation at K101 | 0.15 | 0.05 |
| L99V (0.808) | Altered Coiled coil | 0.43 | 9.10E-03 |
| | Altered Disordered interface | 0.38 | 6.10E-03 |
| | Loss of Relative solvent accessibility | 0.26 | 0.03 |
| | Gain of Allosteric site at Y98 | 0.25 | 0.01 |
| | Gain of Acetylation at K100 | 0.24 | 0.02 |
| | Gain of Methylation at K100 | 0.24 | 1.40E-03 |
| | Altered Metal binding | 0.23 | 0.04 |
| | Gain of Ubiquitylation at K101 | 0.16 | 0.04 |
| T103I (0.749) | Altered Disordered interface | 0.45 | 1.60E-03 |
| | Altered Coiled coil | 0.4 | 0.01 |
| | Altered Ordered interface | 0.3 | 0.02 |
| | Loss of Relative solvent accessibility | 0.27 | 0.02 |
| | Loss of Methylation at K100 | 0.25 | 1.80E-03 |
| | Loss of Allosteric site at Y98 | 0.24 | 0.02 |
| | Loss of Acetylation at K100 | 0.24 | 0.02 |
| | Loss of Ubiquitylation at K101 | 0.15 | 0.05 |

**Table S3.** Cyt-c interaction with ten other proteins along with confidence scores (out of 1.000), analyzed by STRING database.

| Protein | Description | Score |
|---|---|---|
| QCRFS1 | Cytochrome b-c1 complex subunit Rieske | 0.999 |
| CASP9 | Caspase-9 subunit p10 | 0.999 |
| APAF1 | Apoptotic protease-activating factor 1 | 0.999 |
| MT-CYB | Cytochrome b | 0.998 |
| BCL2L1 | Bcl-2-like protein 1 | 0.998 |
| CASP3 | Caspase-3 subunit p12 | 0.997 |
| BCL2 | Apoptosis regulator Bcl-2 | 0.997 |
| CYB5B | Cytochrome b5 type B | 0.996 |
| CYB5A | Cytochrome b5 | 0.996 |

**Table S4.** Overall percentage of secondary structure distribution of WT and mutants during 500 ns simulation.

| Protein | Parallel β-sheet | Anti-parallel β-sheet | 3-10 Helix | α-helix | π-helix | Turn | Bend | Coil |
|---|---|---|---|---|---|---|---|---|
| WT | 0.779 | 2.316 | 1.479 | 42.672 | 0.000 | 13.784 | 10.425 | 28.545 |
| T20I | 0.905 | 2.212 | 1.697 | 42.274 | 0.000 | 12.938 | 11.857 | 28.118 |
| V21G | 0.855 | 2.379 | 1.658 | 42.718 | 0.000 | 14.224 | 9.608 | 28.557 |
| H27Y | 1.267 | 2.075 | 1.808 | 42.679 | 0.000 | 13.730 | 10.946 | 27.494 |
| N32H | 0.205 | 2.046 | 1.037 | 42.842 | 0.000 | 13.812 | 11.288 | 28.771 |
| L33V | 0.721 | 2.091 | 1.201 | 42.443 | 0.000 | 13.530 | 11.384 | 28.632 |
| Y49H | 0.835 | 2.321 | 1.595 | 43.535 | 0.000 | 13.516 | 10.021 | 28.177 |
| G42S | 1.120 | 2.409 | 1.809 | 42.572 | 0.000 | 13.513 | 11.260 | 27.317 |
| A52T | 0.871 | 2.195 | 1.667 | 42.523 | 0.000 | 14.566 | 9.450 | 28.729 |
| A52V | 1.137 | 2.201 | 1.849 | 43.330 | 0.000 | 13.005 | 11.004 | 27.474 |
| R92G | 0.517 | 2.134 | 1.228 | 43.295 | 0.000 | 13.801 | 10.513 | 28.511 |
| A97D | 0.802 | 2.130 | 0.631 | 41.403 | 0.000 | 14.329 | 11.050 | 29.656 |
| Y98H | 1.045 | 2.099 | 1.565 | 42.623 | 0.000 | 14.899 | 9.511 | 28.259 |
| L99V | 1.174 | 1.833 | 2.085 | 42.714 | 0.000 | 13.224 | 11.816 | 27.154 |
| T103I | 0.814 | 2.186 | 1.551 | 43.790 | 0.000 | 14.228 | 9.980 | 27.450 |

**Table S5.** The RMSD analysis of WT and mutant Cyt-c (whole protein and Ω loops) over a 500 ns simulation, presented as median (95% CI) and mean ± SD. Statistical comparisons with WT were performed using the Kruskal-Wallis test followed by Dunn's multiple comparisons to calculate p-values.

| Protein | Cα | | | Proximal Ω loop | | | Central Ω loop | | | Distal Ω loop | | |
|---|---|---|---|---|---|---|---|---|---|---|---|---|
| | Median (95% CI) | Mean ± SD | p-value | Median (95% CI) | Mean ± SD | p-value | Median (95% CI) | Mean ± SD | p-value | Median (95% CI) | Mean ± SD | p-value |
| WT | 1.436 (1.476-1.483) | 1.480 ± 0.405 | | 1.542 (1.558-1.568) | 1.563 ± 0.524 | | 0.856 (0.89-0.893) | 0.892 ± 0.206 | | 0.600 (0.617-0.619) | 0.618 ± 0.127 | |
| T20I | 2.076 (1.958-1.968) | 1.963 ± 0.576 | <0.0001 | 2.299 (2.097-2.108) | 2.102 ± 0.657 | <0.0001 | 0.997 (1.256-1.265) | 1.260 ± 0.537 | <0.0001 | 0.79 (0.792-0.794) | 0.793 ± 0.138 | <0.0001 |
| V21G | 1.849 (1.704-1.715) | 1.709 ± 0.604 | <0.0001 | 1.611 (1.438-1.446) | 1.442 ± 0.454 | <0.0001 | 1.314 (1.283-1.292) | 1.287 ± 0.534 | <0.0001 | 0.753 (0.761-0.764) | 0.762 ± 0.142 | <0.0001 |
| H27Y | 1.340 (1.397-1.401) | 1.399 ± 0.267 | <0.0001 | 1.520 (1.484-1.488) | 1.486 ± 0.217 | <0.0001 | 0.814 (0.827-0.829) | 0.828 ± 0.124 | <0.0001 | 0.689 (0.697-0.699) | 0.698 ± 0.141 | <0.0001 |
| N32H | 1.923 (1.885-1.893) | 1.889 ± 0.439 | <0.0001 | 2.471 (2.354-2.364) | 2.359 ± 0.599 | <0.0001 | 0.775 (0.826-0.829) | 0.828 ± 0.193 | <0.0001 | 0.820 (0.875-0.878) | 0.876 ± 0.206 | <0.0001 |
| L33V | 1.764 (1.749-1.756) | 1.752 ± 0.390 | <0.0001 | 1.743 (1.843-1.853) | 1.848 ± 0.612 | <0.0001 | 0.738 (0.751-0.753) | 0.752 ± 0.130 | <0.0001 | 0.693 (0.691-0.693) | 0.692 ± 0.099 | <0.0001 |
| G42S | 1.550 (1.610-1.621) | 1.616 ± 0.640 | <0.0001 | 1.454 (1.332-1.340) | 1.336 ± 0.453 | <0.0001 | 1.454 (1.332-1.340) | 1.336 ± 0.453 | <0.0001 | 0.778 (0.802-0.805) | 0.804 ± 0.167 | <0.0001 |
| Y49H | 1.501 (1.502-1.507) | 1.504 ± 0.319 | <0.0001 | 1.702 (1.652-1.659) | 1.655 ± 0.404 | <0.0001 | 1.702 (1.652-1.659) | 1.655 ± 0.404 | <0.0001 | 0.616 (0.644-0.647) | 0.645 ± 0.155 | <0.0001 |
| A52T | 1.268 (1.690-1.707) | 1.698 ± 0.952 | <0.0001 | 1.028 (1.144-1.151) | 1.147 ± 0.427 | <0.0001 | 1.028 (1.144-1.151) | 1.147 ± 0.427 | <0.0001 | 0.628 (0.654-0.656) | 0.655 ± 0.150 | <0.0001 |
| A52V | 1.726 (1.583-1.591) | 1.587 ± 0.452 | <0.0001 | 1.903 (1.710-1.722) | 1.716 ± 0.681 | <0.0001 | 1.903 (1.710-1.722) | 1.716 ± 0.681 | <0.0001 | 0.676 (0.683-0.685) | 0.684 ± 0.114 | <0.0001 |
| R92G | 1.503 (1.517-1.523) | 1.520 ± 0.318 | <0.0001 | 1.396 (1.378-1.387) | 1.382 ± 0.468 | <0.0001 | 1.396 (1.378-1.387) | 1.382 ± 0.468 | <0.0001 | 0.603 (0.612-0.614) | 0.613 ± 0.100 | >0.9999 |
| A97D | 2.640 (2.463-2.475) | 2.469 ± 0.661 | <0.0001 | 1.769 (1.645-1.653) | 1.649 ± 0.423 | <0.0001 | 1.214 (1.465-1.475) | 1.470 ± 0.621 | <0.0001 | 0.852 (0.832-0.835) | 0.833 ± 0.166 | <0.0001 |
| Y98H | 1.303 (1.331-1.336) | 1.333 ± 0.310 | <0.0001 | 1.068 (1.046-1.049) | 1.047 ± 0.171 | <0.0001 | 1.060 (1.047-1.052) | 1.049 ± 0.285 | <0.0001 | 0.694 (0.692-0.694) | 0.693 ± 0.120 | <0.0001 |
| L99V | 1.670 (1.665-1.672) | 1.668 ± 0.389 | <0.0001 | 1.790 (1.801-1.810) | 1.806 ± 0.559 | <0.0001 | 1.003 (0.994-0.999) | 0.996 ± 0.291 | <0.0001 | 0.737 (0.739-0.741) | 0.740 ± 0.110 | <0.0001 |
| T103I | 1.581 (1.641-1.647) | 1.644 ± 0.344 | <0.0001 | 1.494 (1.535-1.543) | 1.539 ± 0.488 | <0.0001 | 0.972 (1.021-1.026) | 1.023 ± 0.283 | <0.0001 | 0.675 (0.673-0.675) | 0.674 ± 0.120 | <0.0001 |

**Table S6.** The Rg analysis of WT and mutant Cyt-c (whole protein and Ω loops) over a 500 ns simulation, presented as median (95% CI) and mean ± SD. Statistical comparisons with WT were performed using the Kruskal-Wallis test followed by Dunn's multiple comparisons to calculate p-values.

| Protein | Cα (Å) | | | Proximal loop (Å) | | | Central loop (Å) | | | Distal loop (Å) | | |
|---|---|---|---|---|---|---|---|---|---|---|---|---|
| | Median (95% CI) | Mean ± SD | p-value | Median (95% CI) | Mean ± SD | p-value | Median (95% CI) | Mean ± SD | p-value | Median (95% CI) | Mean ± SD | p-value |
| WT | 12.820 (12.851-12.853) | 12.852 ± 0.123 | | 9.349 (9.36-9.363) | 9.362 ± 0.178 | | 7.222 (7.233-7.235) | 7.234 ± 0.094 | | 6.710 (6.709-6.71) | 6.709 ± 0.071 | |
| T20I | 12.980 (12.966-12.969) | 12.968 ± 0.168 | <0.0001 | 9.434 (9.416-9.419) | 9.418 ± 0.156 | <0.0001 | 7.352 (7.4-7.404) | 7.402 ± 0.200 | <0.0001 | 6.730 (6.75-6.752) | 6.751 ± 0.13 | <0.0001 |
| V21G | 13.011 (12.974-12.977) | 12.975 ± 0.184 | <0.0001 | 9.259 (9.262-9.264) | 9.263 ± 0.131 | <0.0001 | 7.374 (7.366-7.369) | 7.368 ± 0.194 | <0.0001 | 6.701 (6.704-6.705) | 6.704 ± 0.085 | <0.0001 |
| H27Y | 12.784 (12.801-12.803) | 12.802 ± 0.095 | <0.0001 | 9.193 (9.205-9.207) | 9.206 ± 0.095 | <0.0001 | 7.212 (7.222-7.224) | 7.223 ± 0.094 | <0.0001 | 6.710 (6.715-6.717) | 6.716 ± 0.093 | 0.0024 |
| N32H | 12.941 (12.936-12.938) | 12.937 ± 0.095 | <0.0001 | 9.584 (9.547-9.551) | 9.549 ± 0.224 | <0.0001 | 7.130 (7.133-7.134) | 7.133 ± 0.072 | <0.0001 | 6.692 (6.708-6.710) | 6.709 ± 0.133 | <0.0001 |
| L33V | 12.836 (12.845-12.847) | 12.846 ± 0.102 | 0.0396 | 9.411 (9.454-9.458) | 9.456 ± 0.25 | <0.0001 | 7.160 (7.16-7.162) | 7.161 ± 0.076 | <0.0001 | 6.642 (6.643-6.644) | 6.643 ± 0.087 | <0.0001 |
| G42S | 12.839 (12.893-12.896) | 12.894 ± 0.185 | <0.0001 | 9.204 (9.224-9.226) | 9.225 ± 0.132 | <0.0001 | 7.422 (7.437-7.439) | 7.438 ± 0.118 | <0.0001 | 6.740 (6.752-6.753) | 6.753 ± 0.101 | <0.0001 |
| Y49H | 12.778 (12.800-12.801) | 12.801 ± 0.097 | <0.0001 | 9.288 (9.294-9.296) | 9.295 ± 0.139 | <0.0001 | 7.259 (7.262-7.263) | 7.263 ± 0.077 | <0.0001 | 6.671 (6.676-6.678) | 6.677 ± 0.093 | <0.0001 |
| A52T | 12.825 (12.912-12.916) | 12.914 ± 0.202 | <0.0001 | 9.200 (9.228-9.231) | 9.229 ± 0.131 | <0.0001 | 7.198 (7.254-7.257) | 7.255 ± 0.19 | 0.0002 | 6.771 (6.793-6.795) | 6.794 ± 0.131 | <0.0001 |
| A52V | 12.903 (12.894-12.896) | 12.895 ± 0.132 | <0.0001 | 9.345 (9.353-9.356) | 9.354 ± 0.222 | <0.0001 | 7.282 (7.289-7.291) | 7.29 ± 0.131 | <0.0001 | 6.661 (6.664-6.665) | 6.664 ± 0.085 | <0.0001 |
| R92G | 12.827 (12.831-12.832) | 12.832 ± 0.082 | <0.0001 | 9.401 (9.44-9.444) | 9.442 ± 0.238 | <0.0001 | 7.179 (7.181-7.183) | 7.182 ± 0.076 | <0.0001 | 6.797 (6.792-6.793) | 6.793 ± 0.083 | <0.0001 |
| A97D | 13.247 (13.187-13.191) | 13.189 ± 0.234 | <0.0001 | 9.320 (9.321-9.323) | 9.322 ± 0.149 | <0.0001 | 7.31 (7.366-7.371) | 7.368 ± 0.26 | <0.0001 | 6.848 (6.834-6.837) | 6.835 ± 0.133 | <0.0001 |
| Y98H | 12.751 (12.759-12.761) | 12.760 ± 0.069 | <0.0001 | 9.157 (9.157-9.158) | 9.157 ± 0.065 | <0.0001 | 7.296 (7.308-7.311) | 7.309 ± 0.13 | <0.0001 | 6.718 (6.722-6.723) | 6.723 ± 0.086 | <0.0001 |
| L99V | 12.931 (12.917-12.919) | 12.918 ± 0.100 | <0.0001 | 9.345 (9.351-9.354) | 9.352 ± 0.169 | 0.0006 | 7.329 (7.351-7.355) | 7.353 ± 0.222 | <0.0001 | 6.651 (6.661-6.663) | 6.662 ± 0.106 | <0.0001 |
| T103I | 12.871 (12.880-12.882) | 12.881 ± 0.112 | <0.0001 | 9.277 (9.28-9.282) | 9.281 ± 0.153 | <0.0001 | 7.189 (7.221-7.224) | 7.223 ± 0.144 | <0.0001 | 6.695 (6.704-6.706) | 6.705 ± 0.114 | <0.0001 |

**Table S7.** The RMSF analysis of WT and mutant Cyt-c (whole protein and Ω loops) over a 500 ns simulation, presented as median (95% CI) and mean ± SD. Statistical comparisons with WT were performed using the Kruskal-Wallis test followed by Dunn's multiple comparisons to calculate p-values.

| Protein | Cα (Å) | | | Proximal Ω loop (Å) | | | Central Ω loop (Å) | | | Distal Ω loop (Å) | | |
|---|---|---|---|---|---|---|---|---|---|---|---|---|
| | Median (95% CI) | Mean ± SD | p-value | Median (95% CI) | Mean ± SD | p-value | Median (95% CI) | Mean ± SD | p-value | Median (95% CI) | Mean ± SD | p-value |
| WT | 0.758 (0.888-1.138) | 1.013 ± 0.642 | | 0.770 (0.83-1.317) | 1.074 ± 0.627 | | 1.439 (1.236-1.582) | 1.409 ± 0.348 | | 0.757 (0.701-0.898) | 0.800 ± 0.177 | |
| T20I | 0.812 (0.959-1.221) | 1.090 ± 0.675 | 0.8171 | 0.812 (0.911-1.634) | 1.272 ± 0.932 | >0.9999 | 1.616 (1.377-1.832) | 1.604 ± 0.458 | >0.9999 | 0.819 (0.787-0.948) | 0.867 ± 0.145 | >0.9999 |
| V21G | 0.862 (1.003-1.273) | 1.138 ± 0.695 | 0.5688 | 0.882 (0.944-1.343) | 1.143 ± 0.515 | 0.6645 | 2.299 (1.82-2.529) | 2.175 ± 0.713 | 0.1898 | 0.911 (0.812-1.016) | 0.914 ± 0.184 | 0.683 |
| H27Y | 0.659 (0.723-0.93) | 0.826 ± 0.532 | 0.0046 | 0.647 (0.68-0.976) | 0.828 ± 0.381 | 0.4784 | 0.792 (0.725-0.944) | 0.834 ± 0.22 | 0.0012 | 0.661 (0.619-0.765) | 0.692 ± 0.132 | 0.3569 |
| N32H | 0.816 (0.931-1.209) | 1.070 ± 0.714 | >0.9999 | 0.967 (1.081-1.808) | 1.445 ± 0.938 | 0.1379 | 0.801 (0.759-1.095) | 0.927 ± 0.338 | 0.0093 | 0.747 (0.674-0.847) | 0.76 ± 0.156 | >0.9999 |
| L33V | 0.723 (0.839-1.126) | 0.983 ± 0.739 | 0.9846 | 1.018 (1.052-1.756) | 1.404 ± 0.909 | 0.3087 | 0.846 (0.743-0.994) | 0.868 ± 0.253 | 0.0034 | 0.611 (0.563-0.702) | 0.633 ± 0.125 | 0.0289 |
| G42S | 0.821 (0.944-1.196) | 1.070 ± 0.647 | >0.9999 | 0.923 (0.848-1.25) | 1.049 ± 0.518 | >0.9999 | 1.557 (1.435-1.885) | 1.66 ± 0.453 | >0.9999 | 0.863 (0.789-1.007) | 0.898 ± 0.197 | >0.9999 |
| Y49H | 0.677 (0.774-1.003) | 0.889 ± 0.587 | 0.086 | 0.734 (0.817-1.429) | 1.123 ± 0.79 | >0.9999 | 1.015 (0.904-1.20) | 1.052 ± 0.298 | 0.0796 | 0.628 (0.601-0.746) | 0.674 ± 0.131 | 0.3911 |
| A52T | 0.881 (1.015-1.267) | 1.141 ± 0.647 | 0.1422 | 0.86 (0.906-1.298) | 1.102 ± 0.506 | 0.9572 | 2.134 (1.72-2.384) | 2.052 ± 0.667 | 0.1186 | 0.985 (0.922-1.069) | 0.996 ± 0.133 | 0.0506 |
| A52V | 0.772 (0.871-1.117) | 0.994 ± 0.634 | >0.9999 | 0.915 (0.983-1.738) | 1.**361** ± 0.975 | 0.865 | 1.325 (1.158-1.436) | 1.297 ± 0.28 | >0.9999 | 0.705 (0.673-0.826) | 0.750 ± 0.138 | >0.9999 |
| R92G | 0.655 (0.775-1.028) | 0.902 ± 0.651 | 0.0379 | 0.828 (0.946-1.619) | 1.283 ± 0.867 | >0.9999 | 0.741 (0.688-0.922) | 0.805 ± 0.235 | 0.0002 | 0.557 (0.529-0.641) | 0.585 ± 0.100 | 0.0068 |
| A97D | 0.926 (1.091-1.42) | 1.255 ± 0.846 | <0.0001 | 0.894 (0.957-1.313) | 1.135 ± 0.459 | 0.2173 | 1.684 (1.413-1.872) | 1.642 ± 0.462 | 0.5157 | 0.922 (0.904-1.062) | 0.983 ± 0.143 | 0.0162 |
| Y98H | 0.681 (0.759-0.938) | 0.848 ± 0.461 | 0.0898 | 0.782 (0.692-0.88) | 0.786 ± 0.242 | >0.9999 | 1.347 (1.231-1.538) | 1.384 ± 0.309 | >0.9999 | 0.671 (0.627-0.792) | 0.710 ± 0.149 | 0.4384 |
| L99V | 0.79 (0.921-1.169) | 1.045 ± 0.639 | >0.9999 | 0.868 (0.962-1.593) | 1.277 ± 0.814 | 0.9165 | 1.328 (1.124-1.424) | 1.274 ± 0.301 | 0.9415 | 0.762 (0.710-0.863) | 0.786 ± 0.138 | >0.9999 |
| T103I | 0.772 (0.847-1.063) | 0.955 ± 0.556 | >0.9999 | 0.770 (0.856-1.453) | 1.154 ± 0.769 | >0.9999 | 1.217 (1.03-1.282) | 1.156 ± 0.254 | 0.1092 | 0.706 (0.672-0.832) | 0.752 ± 0.145 | >0.9999 |

**Table S8.** The distance analysis for proximal, central, and distal Ω-loop to heme Fe for WT and mutant Cyt-c ) over a 500 ns simulation, presented as median (95% CI) and mean ± SD. Statistical comparisons with WT were performed using the Kruskal-Wallis test followed by Dunn's multiple comparisons to calculate p-values.

| Protein | Proximal loop - Fe (Å) | | Central loop - Fe (Å) | | Distal loop - Fe (Å) | |
|---|---|---|---|---|---|---|
| | Median (95% CI) | Mean ± SD | Median (95% CI) | Mean ± SD | Median (95% CI) | Mean ± SD |
| WT | 7.797 (7.858-7.862) | 7.860 ± 0.260 | 11.868 (11.915-11.92) | 11.917 ± 0.300 | 8.149 (8.148-8.151) | 8.150 ± 0.171 |
| T20I | 7.971 (7.973-7.977) | 7.975 ± 0.217 | 12.066 (12.216-12.228) | 12.222 ± 0.710 | 8.174 (8.17-8.173) | 8.172 ± 0.194 |
| V21G | 7.801 (7.824-7.828) | 7.826 ± 0.223 | 13.064 (13.023-13.043) | 13.033 ± 1.100 | 8.22 (8.218-8.221) | 8.219 ± 0.172 |
| H27Y | 7.808 (7.808-7.812) | 7.81 ± 0.185 | 11.776 (11.769-11.773) | 11.771 ± 0.202 | 8.101 (8.103-8.106) | 8.104 ± 0.179 |
| N32H | 7.922 (7.929-7.933) | 7.931 ± 0.225 | 11.755 (11.752-11.755) | 11.753 ± 0.166 | 8.262 (8.26-8.263) | 8.261 ± 0.184 |
| L33V | 7.674 (7.719-7.724) | 7.722 ± 0.294 | 11.746 (11.736-11.740) | 11.738 ± 0.225 | 8.245 (8.268-8.272) | 8.27 ± 0.263 |
| G42S | 7.832 (7.837-7.840) | 7.838 ± 0.162 | 11.990 (12.219-12.235) | 12.227 ± 0.879 | 8.181 (8.184-8.187) | 8.186 ± 0.183 |
| Y49H | 7.946 (7.969-7.973) | 7.971 ± 0.245 | 11.721 (11.746-11.752) | 11.749 ± 0.300 | 8.194 (8.196-8.198) | 8.197 ± 0.145 |
| A52T | 7.764 (7.860-7.866) | 7.863 ± 0.315 | 12.065 (12.603-12.622) | 12.613 ± 1.082 | 8.080 (8.089-8.092) | 8.091 ± 0.178 |
| A52V | 8.073 (8.060-8.066) | 8.063 ± 0.31 | 12.138 (12.153-12.159) | 12.156 ± 0.350 | 8.206 (8.206-8.209) | 8.207 ± 0.161 |
| R92G | 7.829 (7.858-7.862) | 7.860 ± 0.245 | 11.781 (11.780-11.783) | 11.782 ± 0.169 | 8.070 (8.075-8.077) | 8.076 ± 0.157 |
| A97D | 7.956 (7.947-7.951) | 7.949 ± 0.234 | 12.420 (12.395-12.405) | 12.400 ± 0.614 | 8.150 (8.157-8.161) | 8.159 ± 0.204 |
| Y98H | 7.751 (7.753-7.755) | 7.754 ± 0.145 | 11.99 (12.136-12.145) | 12.140 ± 0.535 | 8.120 (8.119-8.122) | 8.12 ± 0.165 |
| L99V | 8.214 (8.172-8.177) | 8.175 ± 0.304 | 11.861 (11.902-11.906) | 11.904 ± 0.263 | 8.222 (8.217-8.219) | 8.218 ± 0.162 |
| T103I | 7.777 (7.780-7.783) | 7.781 ± 0.163 | 11.793 (11.906-11.913) | 11.909 ± 0.429 | 8.183 (8.178-8.182) | 8.180 ± 0.201 |

**Table S9.** The median (95% CI) and mean ± SD of distances between H-bonded residues between Tyr68-Asn53, Tyr68-Met81, and Tyr68-Thr79 of WT and mutant Cyt-c over 500 ns MD simulation. Note that amino acid numbering differs by +1 from that of the PDB file (3ZCF).

| Protein | Tyr68 - Asn53 (Å) | | Tyr68 - Met81 (Å) | | Tyr68 - Thr79 (Å) | |
|---|---|---|---|---|---|---|
| | Median (95% CI) | Mean ± SD | Median (95% CI) | Mean ± SD | Median (95% CI) | Mean ± SD |
| WT | 11.213 (11.588-11.604) | 11.596 ± 0.927 | 9.969 (9.985-9.989) | 9.987 ± 0.227 | 10.260 (10.285-10.289) | 10.287 ± 0.194 |
| T20I | 12.370 (12.758-12.788) | 12.773 ± 1.674 | 9.924 (9.946-9.951) | 9.949 ± 0.292 | 10.248 (10.283-10.287) | 10.285 ± 0.252 |
| V21G | 13.684 (13.448-13.482) | 13.465 ± 1.967 | 10.029 (10.053-10.058) | 10.055 ± 0.277 | 10.214 (10.241-10.245) | 10.243 ± 0.235 |
| H27Y | 11.145 (11.149-11.153) | 11.151 ± 0.276 | 9.960 (9.985-9.991) | 9.988 ± 0.308 | 10.277 (10.325-10.329) | 10.327 ± 0.272 |
| N32H | 11.126 (11.13-11.134) | 11.132 ± 0.239 | 9.705 (9.715-9.720) | 9.717 ± 0.268 | 10.224 (10.244-10.248) | 10.246 ± 0.204 |
| L33V | 11.309 (11.335-11.342) | 11.338 ± 0.367 | 10.123 (10.188-10.196) | 10.192 ± 0.428 | 10.572 (10.657-10.666) | 10.662 ± 0.494 |
| G42S | 11.944 (12.422-12.449) | 12.435 ± 1.558 | 10.003 (10.031-10.036) | 10.033 ± 0.307 | 10.340 (10.393-10.398) | 10.395 ± 0.310 |
| Y49H | 11.098 (11.330-11.342) | 11.336 ± 0.66 | 9.844 (9.853-9.857) | 9.855 ± 0.231 | 10.205 (10.212-10.215) | 10.214 ± 0.183 |
| A52T | 11.705 (12.724-12.758) | 12.741 ± 1.959 | 10.163 (10.193-10.199) | 10.196 ± 0.365 | 10.401 (10.465-10.471) | 10.468 ± 0.340 |
| A52V | 12.105 (12.245-12.262) | 12.253 ± 0.948 | 9.917 (9.920-9.924) | 9.922 ± 0.230 | 10.193 (10.203-10.206) | 10.205 ± 0.194 |
| R92G | 10.936 (10.942-10.946) | 10.944 ± 0.228 | 9.760 (9.773-9.777) | 9.775 ± 0.253 | 10.177 (10.19-10.194) | 10.192 ± 0.183 |
| A97D | 12.139 (12.361-12.383) | 12.372 ± 1.283 | 9.963 (9.982-9.988) | 9.985 ± 0.321 | 10.277 (10.316-10.321) | 10.319 ± 0.267 |
| Y98H | 11.276 (11.691-11.71) | 11.700 ± 1.101 | 9.932 (9.938-9.942) | 9.940 ± 0.229 | 10.234 (10.245-10.249) | 10.247 ± 0.191 |
| L99V | 12.029 (12.079-12.096) | 12.087 ± 1.003 | 10.046 (10.05-10.055) | 10.052 ± 0.261 | 10.217 (10.225-10.229) | 10.227 ± 0.200 |
| T103I | 11.164 (11.638-11.658) | 11.648 ± 1.127 | 9.851 (9.851-9.856) | 9.854 ± 0.270 | 10.238 (10.247-10.25) | 10.249 ± 0.185 |

**Table S10.** The median (95% CI) and mean ± SD calculations for cavity A (Ala51–Gly78 Cα) and cavity B (Asn32–Ala44 Cα) of WT and mutant Cyt-c over 500 ns MD simulation. Note that amino acid numbering differs by +1 from that of the PDB file (3ZCF).

| Protein | Cavity A (Ala51–Gly78 Cα) (Å) | | Cavity B (Asn32–Ala44 Cα) (Å) | |
|---|---|---|---|---|
| | Median (95% CI) | Mean ± SD | Median (95% CI) | Mean ± SD |
| WT | 8.265 (8.618-8.636) | 8.627 ± 1.021 | 7.957 (7.874-7.889) | 7.881 ± 0.857 |
| T20I | 9.117 (9.399-9.425) | 9.412 ± 1.509 | 7.077 (7.185-7.199) | 7.192 ± 0.83 |
| V21G | 10.265 (10.467-10.509) | 10.488 ± 2.374 | 7.393 (7.605-7.623) | 7.614 ± 1.001 |
| H27Y | 8.23 (8.292-8.3) | 8.296 ± 0.449 | 8.133 (8.1-8.113) | 8.107 ± 0.696 |
| N32H | 8.143 (8.163-8.169) | 8.166 ± 0.344 | 7.732 (7.694-7.704) | 7.699 ± 0.556 |
| L33V | 8.08 (8.106-8.112) | 8.109 ± 0.328 | 8.378 (8.192-8.208) | 8.2 ± 0.915 |
| G42S | 9.077 (9.67-9.707) | 9.688 ± 2.105 | 7.597 (7.714-7.727) | 7.721 ± 0.704 |
| Y49H | 8.23 (8.432-8.444) | 8.438 ± 0.713 | 6.82 (6.738-6.753) | 6.746 ± 0.848 |
| A52T | 8.986 (10.439-10.489) | 10.464 ± 2.864 | 8.058 (8.069-8.084) | 8.076 ± 0.833 |
| A52V | 9.825 (9.886-9.905) | 9.895 ± 1.096 | 7.413 (7.498-7.513) | 7.506 ± 0.863 |
| R92G | 8.125 (8.14-8.146) | 8.143 ± 0.321 | 8.073 (8.03-8.043) | 8.036 ± 0.722 |
| A97D | 8.409 (8.687-8.704) | 8.695 ± 0.983 | 8.035 (8.175-8.191) | 8.183 ± 0.899 |
| Y98H | 8.192 (8.683-8.707) | 8.695 ± 1.387 | 7.954 (8.033-8.051) | 8.042 ± 1.006 |
| L99V | 9.022 (9.159-9.181) | 9.17 ± 1.265 | 7.518 (7.506-7.526) | 7.516 ± 1.152 |
| T103I | 8.209 (8.609-8.628) | 8.618 ± 1.078 | 8.114 (8.139-8.152) | 8.145 ± 0.781 |

**Table S11.** The median (95% CI) and mean ± SD of the distance between His19 (NE2) - heme Fe and Met81 (SD) - heme Fe of WT and mutant Cyt-c over 500 ns MD simulation. Note that amino acid numbering differs by +1 from that of the PDB file (3ZCF).

| Protein | His19 - Heme Fe (Å) | | Met81 - Heme Fe (Å) | |
|---|---|---|---|---|
| | Median (95% CI) | Mean ± SD | Median (95% CI) | Mean ± SD |
| WT | 2.186 (2.186-2.187) | 2.186 ± 0.037 | 2.385 (2.385-2.386) | 2.385 ± 0.018 |
| T20I | 2.185 (2.185-2.186) | 2.185 ± 0.044 | 2.385 (2.385-2.386) | 2.385 ± 0.022 |
| V21G | 2.187 (2.186-2.187) | 2.187 ± 0.044 | 2.385 (2.384-2.385) | 2.385 ± 0.022 |
| H27Y | 2.184 (2.185-2.185) | 2.185 ± 0.044 | 2.385 (2.385-2.385) | 2.385 ± 0.022 |
| N32H | 2.208 (2.207-2.208) | 2.208 ± 0.046 | 2.387 (2.387-2.387) | 2.387 ± 0.022 |
| L33V | 2.200 (2.200-2.201) | 2.200 ± 0.045 | 2.386 (2.386-2.387) | 2.386 ± 0.022 |
| G42S | 2.177 (2.177-2.177) | 2.177 ± 0.044 | 2.385 (2.384-2.385) | 2.385 ± 0.022 |
| Y49H | 2.179 (2.179-2.179) | 2.179 ± 0.044 | 2.385 (2.385-2.385) | 2.385 ± 0.022 |
| A52T | 2.184 (2.184-2.185) | 2.184 ± 0.044 | 2.386 (2.386-2.386) | 2.386 ± 0.022 |
| A52V | 2.186 (2.186-2.187) | 2.186 ± 0.045 | 2.385 (2.385-2.385) | 2.385 ± 0.022 |
| R92G | 2.192 (2.192-2.193) | 2.192 ± 0.046 | 2.387 (2.387-2.388) | 2.387 ± 0.022 |
| A97D | 2.190 (2.189-2.190) | 2.190 ± 0.045 | 2.385 (2.385-2.385) | 2.385 ± 0.022 |
| Y98H | 2.178 (2.178-2.178) | 2.178 ± 0.044 | 2.385 (2.385-2.386) | 2.385 ± 0.022 |
| L99V | 2.187 (2.187-2.187) | 2.187 ± 0.044 | 2.385 (2.385-2.385) | 2.385 ± 0.022 |
| T103I | 2.181 (2.180-2.182) | 2.181 ± 0.062 | 2.385 (2.384-2.385) | 2.385 ± 0.031 |
